# A systematic comparison reveals substantial differences in chromosomal versus episomal encoding of enhancer activity

**DOI:** 10.1101/061606

**Authors:** Fumitaka Inoue, Martin Kircher, Beth Martin, Gregory M. Cooper, Daniela M. Witten, Michael T. McManus, Nadav Ahituv, Jay Shendure

**Author notes:** These authors contributed equally to this work (Fumitaka Inoue, Martin Kircher).

## Abstract

Candidate enhancers can be identified on the basis of chromatin modifications, the binding of chromatin modifiers and transcription factors and cofactors, or chromatin accessibility. However, validating such candidates as bona fide enhancers requires functional characterization, typically achieved through reporter assays that test whether a sequence can drive expression of a transcriptional reporter via a minimal promoter. A longstanding concern is that reporter assays are mainly implemented on episomes, which are thought to lack physiological chromatin. However, the magnitude and determinants of differences in *cis*-regulation for regulatory sequences residing in episomes versus chromosomes remain almost completely unknown. To address this question in a systematic manner, we developed and applied a novel lentivirus-based massively parallel reporter assay (lentiMPRA) to directly compare the functional activities of 2,236 candidate liver enhancers in an episomal versus a chromosomally integrated context. We find that the activities of chromosomally integrated sequences are substantially different from the activities of the identical sequences assayed on episomes, and furthermore are correlated with different subsets of ENCODE annotations. The results of chromosomally-based reporter assays are also more reproducible and more strongly predictable by both ENCODE annotations and sequence-based models. With a linear model that combines chromatin annotations and sequence information, we achieve a Pearson’s R^2^ of 0.347 for predicting the results of chromosomally integrated reporter assays. This level of prediction is better than with either chromatin annotations or sequence information alone and also outperforms predictive models of episomal assays. Our results have broad implications for how *cis*-regulatory elements are identified, prioritized and functionally validated.

## Introduction

An enhancer is defined as a short region of DNA that can potentiate the expression of a gene, independent of its orientation and flexible with respect to its position relative to the transcriptional start site (Banerji et al. 1981; Moreau et al. 1981). Enhancers are thought to be modular, in the sense that they are active in heterologous sequence contexts and in that multiple enhancers may additively dictate the overall expression pattern of a gene (Shlyueva et al. 2014). They act through the binding of transcription factors, which recruit histone modifying factors, such as histone acetyltransferase (HAT) or histone methyltransferase (HMT). Enhancers are also associated with chromatin remodeling factors (e.g. SWI/SNF) and the cohesin complex, which are involved in regulating chromatin structure and accessibility (Schmidt et al. 2010; Euskirchen et al. 2011; Faure et al. 2012).

Antibodies against specific transcription factors (TFs), histone modifications or transcriptional co-activators are commonly used for chromatin immunoprecipitation followed by massively parallel sequencing (ChIP-seq) to identify candidate enhancers in a genome-wide manner. For example, the ENCODE Consortium and other efforts have identified thousands of candidate enhancers in mammalian genomes on the basis of such marks or their correlates (e.g. p300 ChIP-seq; H3K27ac ChIP-seq; DNaseI hypersensitivity) in diverse cell lines and tissues (Visel et al. 2009; Dunham I et al. 2012). However, a major limitation of such assays is that they reflect biochemical marks that are correlated with enhancer activity, rather than directly showing that any particular sequence actually functions as an enhancer. In other words, while such assays yield genome-wide catalogs of potential enhancers, they do not definitively predict bona fide enhancers nor precisely define their boundaries.

For decades, the primary means of functionally validating enhancers has been the episomal reporter assay. The standard approach is to relocate the candidate enhancer sequence to an episomal vector, adjacent to a minimal promoter driving expression of a reporter gene, e.g. luciferase or others. More recently, massively parallel reporter assays (MPRAs) have enabled the functional characterization of *cis*-regulatory elements including enhancers in a high-throughput manner. MPRAs use sequencing-based quantification of reporter barcodes to enable multiplexing of the reporter assay (Patwardhan et al. 2009). MPRAs have been used primarily in an episomal manner for the saturation mutagenesis of promoters and enhancers (Patwardhan et al. 2009; Kinney et al. 2010; Melnikov et al. 2012; Patwardhan et al. 2012), for exploring the grammatical rules of promoters and enhancers (Smith et al. 2013; Sharon et al. 2014), and for testing of thousands of enhancer candidates in different cells or tissues (Kwasnieski et al. 2012; Arnold et al. 2013; Kheradpour et al. 2013; Arnold et al. 2014; Shlyueva et al. 2014; Savic et al. 2015; White 2015). Adeno-associated virus (AAV) MPRAs have also been developed, allowing these assays to be carried out *in vivo* and to perform reporter assays within target cells and tissues that are difficult to transduce, such as the brain (Shen et al. 2016), although these do not involve genomic integration.

Despite their widespread use to validate enhancers and other cis-regulatory elements, a longstanding concern about reporter assays is that they are almost always carried out via transient transfection of non-integrating episomes. It is unknown whether transiently transfected sequences are chromatinized in a way that makes them appropriate models for endogenous gene expression from chromosomes (Smith and Hager 1997); but to the extent that this question has been explored, there are differences. For example, work from Archer, Hager and colleagues using the mouse mammary tumor virus (*MMTV*) promoter as a model shows differences in histone H1 stoichiometry and nucleosome positioning resulting in an inability of episomal assays to reliably assay cooperative TF binding (Archer et al. 1992; Smith and Hager 1997; Hebbar and Archer 2007; Hebbar and Archer 2008). In another work, the chromatin structure of transiently transfected non-replicating plasmid DNA was observed to be differently fragmented than endogenous chromatin by micrococcal nuclease, and along with other data supports a model in which atypical chromatin might be induced by association of episomes with nuclear structures (Jeong and Stein 1994). However, the extent to which these factors operate to confound the results of enhancer reporter assays more broadly, for categorical (i.e. is a particular sequence an enhancer?), qualitative (i.e. in what tissues is an element an enhancer?), and quantitative validation (i.e. what level of activation does a particular sequence confer?), has yet to be systematically investigated.

To address these questions, we developed lentiviral MPRA (lentiMPRA), a technology that uses lentivirus to integrate enhancer MPRA libraries into the genome. To overcome the substantial position-effect variegation observed by others in attempting to use lentiviral infection for MPRA (Murtha et al. 2014), we employed a flanking antirepressor element (#40) and a scaffold-attached region (SAR) (Klehr et al. 1991; Kwaks et al. 2003) on either side of our construct. In addition, we relied on as many as 100 independent reporter barcode sequences per assayed candidate enhancer sequence, integrated at diverse sites. The resulting system allows for high-throughput, highly reproducible and quantitative measurement of the regulatory potential of candidate enhancers in a chromosomally integrated context. Furthermore, the cell-type range of lentivirus transduction is much broader than transfection, e.g. permitting MPRAs to be conducted in neurons, primary cells or organoids.

By using integration-competent vs. integration-defective components of the lentiviral system, we directly compared the functional activities of 2,236 candidate liver enhancers in a chromosomally integrated versus an episomal context in the human liver hepatocellular carcinoma cell line, HepG2. We find that the activities of chromosomally integrated sequences are substantially different from the activities of the identical sequences assayed episomally, and are correlated with different subsets of ENCODE annotations. We also find that the results of chromosomally-based reporter assays are more reproducible and more strongly predicted by ENCODE annotations and sequence-based models.

## Results

### Construction and validation of the lentiMPRA vector

The potential for confounding of lentiviral assays by site-of-integration effects was demonstrated by a recent MPRA study (Functional Identification of Regulatory Elements Within Accessible Chromatin or FIREWACh) that used lentiviral infection and found that 26% of positive controls did not show activated GFP expression, while other measures estimated a false positive rate of 22% (Murtha et al. 2014). We therefore constructed a lentiviral vector (pLS-mP) that contains a minimal promoter (mP) and the enhanced green fluorescent protein (EGFP) gene flanked on one side by the antirepressor element #40 and the other by a SAR that reduce site-of-integration effects and provide consistent transgene expression (Fig. 1A, Supplementary File 1) (Klehr et al. 1991; Kwaks et al. 2003; Kissler et al. 2006). In experiments involving chromosomal integration of this enhancer reporter, we confirmed that EGFP is not expressed in the absence of an enhancer, while abundantly expressed under the control SV40 enhancer across a panel of cell lines representing diverse tissues-of-origin. These include: K562 (lymphoblasts), H1-ESC (embryonic stem cells), HeLa-S3 (cervix), HepG2 (hepatocytes), T-47D (epithelial) and Sk-n-sh retinoic acid treated (neuronal) cells (Fig. S1). Furthermore, when SV40 and the *Ltv1* liver enhancer (Patwardhan et al. 2012) are tested without the flanking antirepressor sequences, we observed much lower levels of EGFP expression in HepG2 cells, consistent with our expectation that the antirepressors facilitate robust enhancer-mediated expression from the integrated reporter (Fig. 1B).

**Figure 1.**
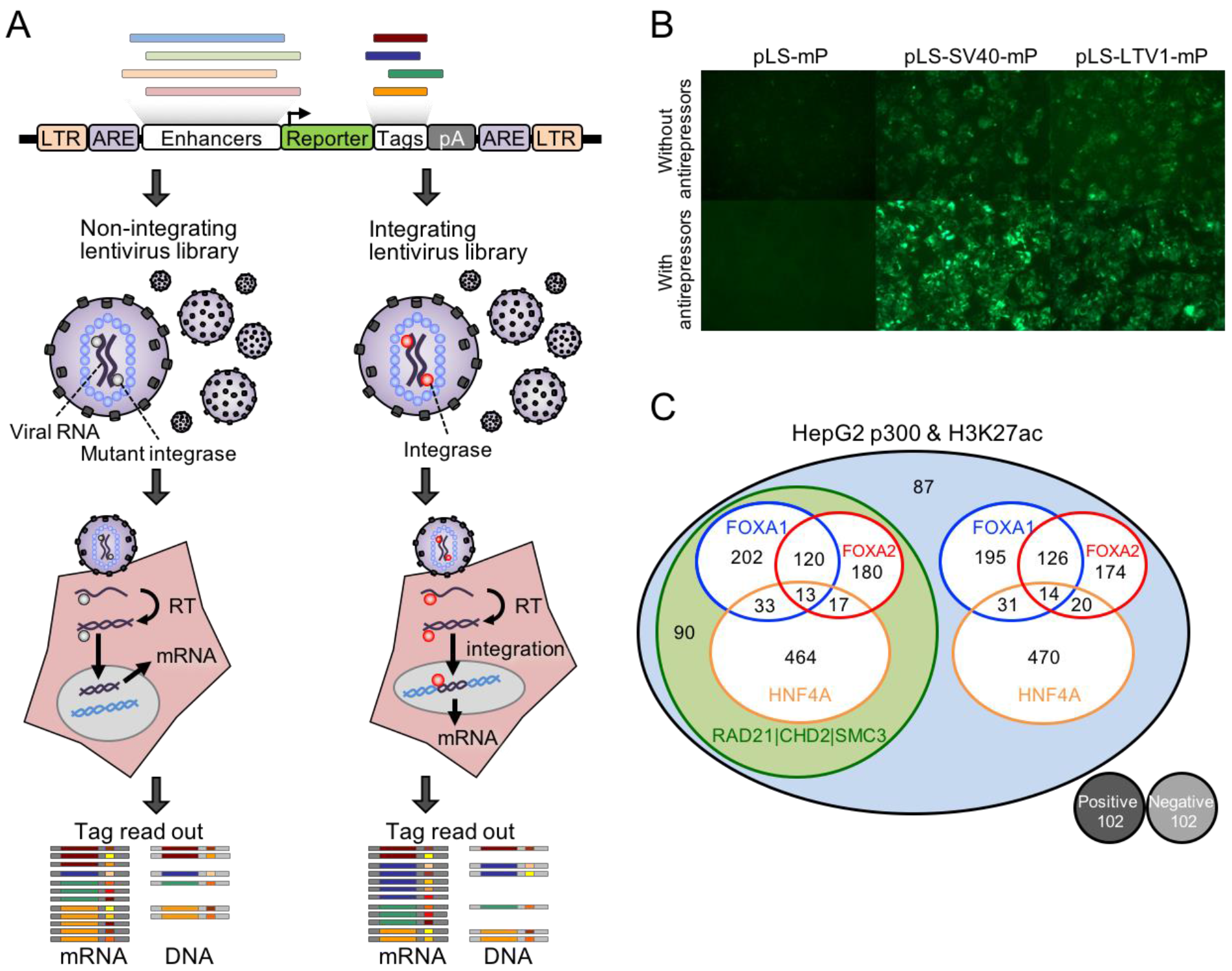
Study design for LentiMPRA. Study design for LentiMPRA. (*A*) Schematic diagram of lentiMPRA. Candidate enhancers and barcode tags were synthesized in tandem as a microarray-derived oligonucleotide library, and cloned into the pLS-mP vector, followed by cloning of a minimal promoter (mP) and reporter (EGFP) between them. The resulting lentiMPRA library was packaged with either wildtype or mutant integrase, and infected into HepG2 cells. Both DNA and mRNA were extracted, and barcode tags were sequenced to test their enhancer activities in an episomal vs. genome integrating manner. (*B*) HepG2 cells infected with lentiviral reporter construct bearing no enhancer (pLS-mP), an SV40 enhancer (pLS-SV40-mP), and LTV1 (pLS-LTV1-mP), a known liver enhancer (Patwardhan et al. 2012), with or without antirepressors. The inclusion of antirepressors results in stronger and more consistent expression, but is still dependent on the presence of an enhancer. (*C*) Venn diagram showing the composition of the lentiMPRA library. 2,236 enhancer candidate sequences were chosen on the basis of having ENCODE HepG2 ChIP-seq peaks for EP300 and H3K27ac marks. The candidates overlapped with or without ChIP-seq peaks for FOXA1, FOXA2, or HNF4A. Half ofthe candidates overlapped with ChIP-seq peaks for RAD21, SMC3, and CHD2. The library included 102 positive and 102 negative controls.

### Design and construction of a library of candidate liver enhancers

To evaluate lentiMPRA, we designed a liver enhancer library that comprises 2,236 candidate sequences and 204 control sequences (Fig. 1C, Supplementary File 2), each 171 bp in length. All enhancer candidate sequences were chosen on the basis of having ENCODE HepG2 ChIP-seq peaks for EP300 and H3K27ac, which are generally indicative of enhancer function (Heintzman et al. 2007; Visel et al. 2009). A subset of candidates (“type 1”) are centered at ChIP-seq peaks for forkhead box A1 (FOXA1) or FOXA2, known liver pioneer transcription factors (Lupien et al. 2008) or hepatocyte nuclear factor 4 alpha (HNF4A), a nuclear receptor involved in lipid metabolism and gluconeogenesis (Watt et al. 2003), while also overlapping with ENCODE-derived ChIP-seq peaks for the cohesin complex (RAD21 and SMC3) or chromodomain helicase DNA binding protein 2 (CHD2), a chromatin remodeler that is part of the SWI/SNF complex. Other subsets of candidates were required to overlap only a liver transcription factor peak (“type 2”), only a chromatin remodeler peak (“type 3”), or neither (“type 4”). The 204 control sequences comprised 200 synthetically designed controls from a previous study (synthetic regulatory element sequences (SRESs); 100 positive & 100 negative) (Smith et al. 2013) and an additional 2 positive (pos1 and pos2) and 2 negative endogenous controls (neg1 and neg2). We confirmed by standard luciferase reporter assay that pos1 and pos2 showed weak and strong enhancer activity, respectively, while neg1 and neg2 showed no activity (Fig. S2).

Each of the 2,440 enhancer candidates or controls was synthesized in *cis* with 100 unique reporter barcodes on a 244,000-feature microarray (Agilent OLS; 15 bp primer + 171 bp enhancer candidate or control + 14 bp spacer + 15 bp barcode + 15 bp primer = 230-mers). The purpose of encoding a large number of barcodes per assayed sequence was to facilitate reproducible and quantitative measurements of regulatory activity, as well as to mitigate against non-uniformity in oligonucleotide synthesis. We cloned these oligonucleotides to a version of the lentiMPRA vector that lacked mP and EGFP reporter. Subsequently, a restriction site in the spacer was used to reinsert the mP + EGFP cassette between the candidate enhancer and barcode, thus positioning the barcode in the 3’ UTR of EGFP (Fig. S3).

To evaluate the quality of the designed oligonucleotides and the representation of individual barcodes, we sequenced the cloned oligonucleotide library (i.e. prior to reinsertion of the mP + EGFP cassette) to a depth of 19.2 million paired-end consensus sequences, 52.6% of which had the expected length. Analysis of these data showed that most molecular copies of a given oligonucleotide are correct, that synthesis errors are distributed evenly along the designed insert sequence, and that single base deletions dominate the observed errors (Fig. S4A). Nonetheless, there was substantial non-uniformity in the library (Fig. S4B). While 90.5% of the 244,000 designed barcodes were observed at least once amongst 11.0 million full-length barcodes sequenced, their abundance is sufficiently dispersed that we estimated that a subset of 56-67% of the designed oligonucleotides would be propagated when maintaining a library complexity of 350,000-600,000 clones.

### Chromosomally integrated versus episomal lentiMPRA

We next sought to directly compare the functional activities of the 2,236 candidate liver enhancer sequences in a chromosomally integrated versus an episomal context. To this end, we packaged the lentiMPRA library with either a wild-type integrase (WT-IN) or a mutant integrase (MT-IN), with the latter allowing for the production of non-integrating lentivirus and transient transgene expression from non-integrated DNA (Leavitt et al. 1996; Nightingale et al. 2006) (Fig. 1A). Because the integrase is not encoded by the lentiMPRA library, this experimental design allows us to test the same exact library in both integrated and non-integrated contexts.

To optimize conditions and reduce background of unintegrated lentivirus in the integrating lentivirus prep, we utilized our positive control virus (pLS-SV40-mP) that was packaged with WT-IN and MT-IN, and examined the viral titer by qPCR for three different volumes (1, 5 and 25 μl per well of a 24-well plate) at four different time points (2-5 days post infection). For the lower volumes (1 and 5 ul), we observed a substantial reduction in total virus amounts at day 4 for both MT-IN and WT-IN that stabilized in the WT-IN only (Fig. S5A). This suggests that the nonintegrated virus declines at this time point, similar to what was previously reported (Butler et al. 2001). For the high volume (25 ul), we did not observe a substantial reduction or stabilization for MT-IN and WT-IN respectively until day 5 (Fig. S5A), suggesting that high amounts of virus would make it difficult to distinguish between integrated and non-integrated virus. We thus decided to obtain DNA/RNA from the cells with the WT-IN liver enhancer library at day 4 when they have an estimated 50 viral particles/cell and the MT-IN library at day 3 when they had an estimated 100 viral particles/cell. The total copy number of viral DNA in the cells infected with the liver enhancer libraries was validated by qPCR (Fig. S5B). During human immunodeficiency virus (HIV) infection, non-integrating virus represents a major portion of the virus at early infection time points and includes linear DNA that is rapidly degraded along with circular DNA containing terminal repeats (1-LTRc and 2-LTRc) (Munir et al. 2013). We further confirmed the copy number of non-integrated virus at our assayed time points by carrying out a qPCR on 2- LTRc, observing the expected low and high amounts of non-integrated virus with WT-IN and MT-IN, respectively (Fig. S5B).

### lentiMPRA on 2,236 candidate liver enhancer sequences

We recovered RNA and DNA from both WT-IN and MT-IN infections (three replicates each consisting of independent infections with the same library), amplified barcodes, and performed sequencing (Illumina NextSeq). We used both the forward and reverse reads to sequence the 15 bp reporter barcodes and obtain consensus sequences. We obtained an average of ~4.1 million raw barcode counts for DNA and an average of ~26 million raw barcode counts for RNA. Across replicates and sample types, 97% of barcodes were the correct length of 15bp. The number of unique sequences was on average ~450,000 for DNA and ~1.2 million for RNA. When clustering sequences with up to one substitution relative to a programmed barcode, the average number of unique sequences reduced to ~280,000 for DNA and ~700,000 for RNA. We speculate that our RNA readouts are impacted by sequence errors to a greater extent due to the reverse transcriptase (RT) step.

We matched the observed barcodes against the designed barcodes and normalized RNA and DNA for different sequencing depths in each sample by dividing counts by the sum of all observed counts and reporting them as counts per million. Only barcodes observed at least once in both RNA and DNA of the same sample were considered. Subsequently, RNA/DNA ratios were calculated. The average Spearman’s rho for DNA counts of the three integrase mutant (MT) experiments was 0.907, and for RNA counts of the MT experiments was 0.982. The average Spearman’s rho values for the wild-type integrase (WT) experiments were 0.864 and 0.979 for DNA and RNA, respectively. These correlations were determined for barcodes observed in pairs of replicates. Scatter plots for the MT and WT experiments are shown in Fig. S6 and Fig. S7, respectively.

While the DNA and RNA counts for individual barcodes are highly correlated between experiments, the noise of each measure results in a poor correlation of RNA/DNA ratios (Fig. S6, Fig. S7). However, there are on average 59-62 barcodes per candidate enhancer sequence (insert) in each replicate (out of 100 barcodes programmed on the array, with ~40% lost during cloning as discussed above) (Fig. S8). To reduce noise, we summed up the RNA or DNA counts across all associated barcodes for each insert observed in a given experiment and recalculated RNA/DNA ratios (Fig. S9). Pairwise-correlations of DNA and RNA counts of replicates are very high (average Spearman’s rho MT-RNA 0.996, MT-DNA 0.994, WT-RNA 0.997 and WT-DNA 0.991). Fig. 2 shows scatter plots and correlation values for per-insert RNA/DNA ratios for the MT and WT experiments. RNA/DNA ratios show markedly improved reproducibility after summing across barcodes, with an average Spearman’s rho of 0.908 (MT) and 0.944 (WT). In all pairwise comparisons of replicates, the integrated (WT) MPRA experiments exhibit a broader dynamic range and greater reproducibility than the episomal (MT) MPRA experiments. We also explored how stable the correlation of RNA/DNA ratios is between replicates by down-sampling the number of barcodes per insert or specifying an exact number of barcodes per insert (Fig. S10). Again, the WT experiments show greater reproducibility, especially for inserts represented by fewer independent barcodes.

**Figure 2.**
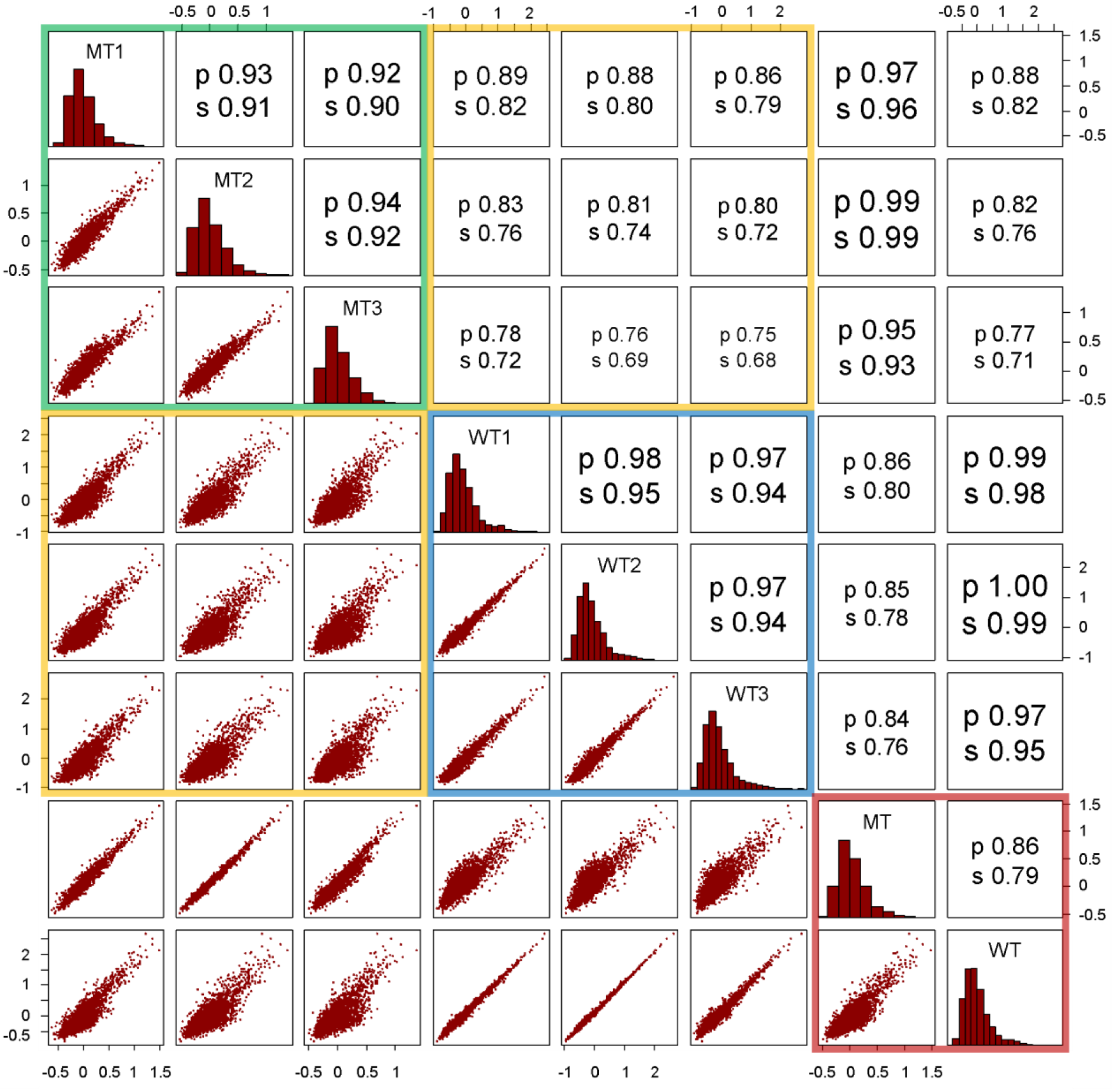
Pairwise correlation of per-insert RNA/DNA ratios between replicates, within and between MT vs. WT experiments. The lower left triangle shows pair-wise scatter plots. The diagonal provides replicate names and the respective histogram of the RNA/DNA ratios for that replicate. The upper triangle provides Pearson (p) and Spearman (s) correlation coefficients. MT vs. MT (green box) or WT vs. WT (blue box) comparisons are substantially more correlated than MT vs. WT (yellow boxes) comparisons, consistent with systematic differences between the episomal vs. integrated contexts for reporter assays that exceed technical noise. The two right-most columns and two bottom-most rows correspond to MT and WT after combining across the three replicates, with the combined MT vs. the combined WT comparison in the red box.

To combine replicates, we normalized the RNA/DNA ratios for inserts observed in each replicate by dividing by their median, and then averaged this normalized RNA/DNA ratio for each insert across replicates (red box in Fig. 2; Fig. 3A). Fig. 3A shows scatter plots of the resulting MT and WT RNA/DNA ratios colored by the type of insert and/or transcription factors considered in the design (Fig. S11 shows RNA/DNA ratio ranges by type of insert). As noted above, we observe a broader dynamic range in the WT experiment. Furthermore, the Spearman correlation between MT and WT is 0.792, which is considerably lower than the correlation observed when correlating replicates of the same experimental type (Spearman correlation of 0.908 (MT) and 0.944 (WT)). This is also the case in pairwise comparisons of MT versus WT replicates (i.e. prior to combining replicates) (yellow boxes in Fig. 2). Overall, these results show that there are substantial differences in regulatory activity between identical sequences assayed in an integrated vs. episomal context.

**Figure 3.**
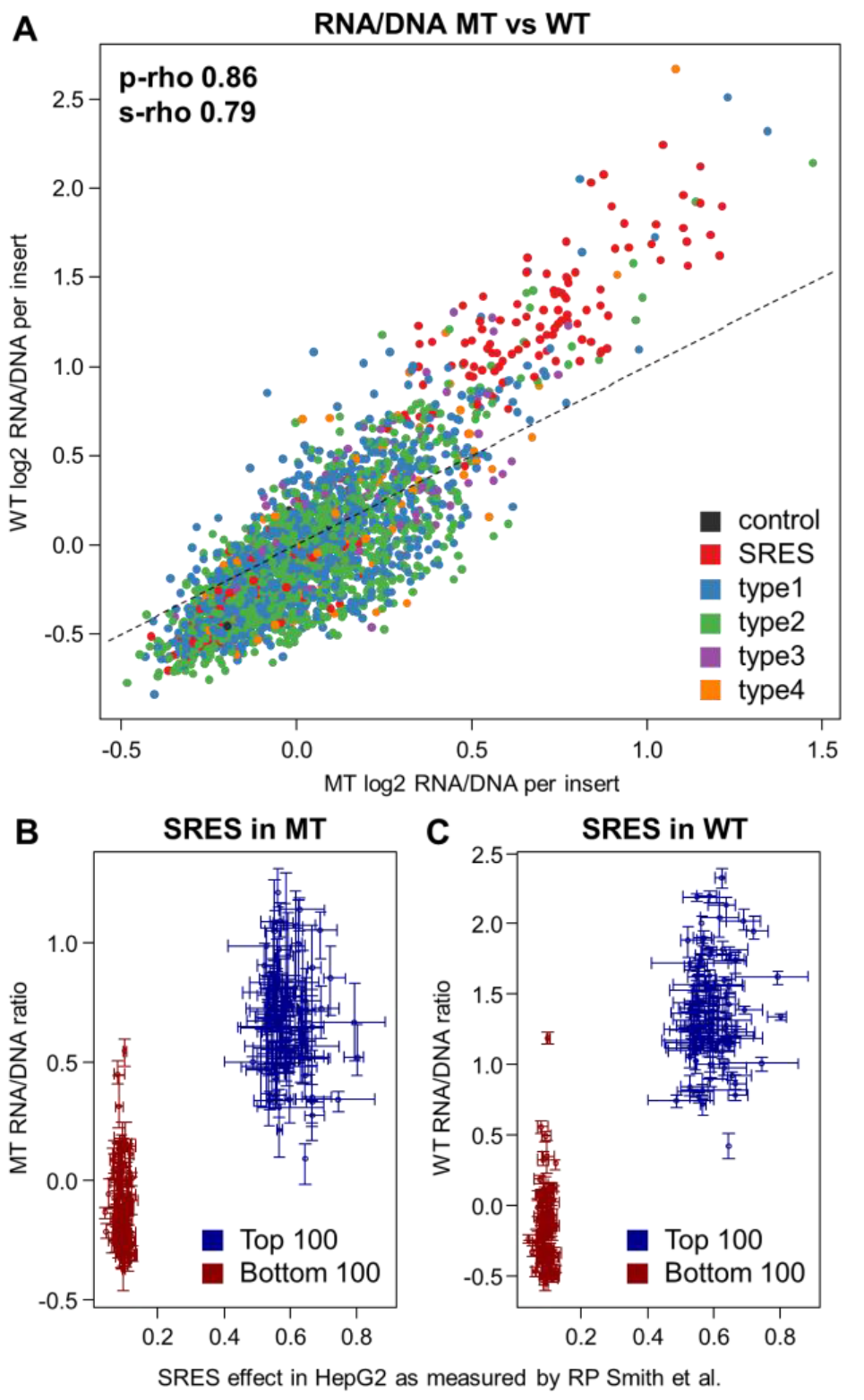
Comparisons between the non-integrating (MT) and integrating (WT) libraries. (*A*) Scatter plot of combined MT vs. WT RNA/DNA ratios. MT ratios show a smaller dynamic range and thus seem compressed compared to WT results. Data points are colored by the type of insert sequence, including two types of controls: a total of four positive and negative controls (black) as well as the highest 100 and lowest 100 synthetic regulatory element sequences (SRES, red) identified by Smith *etal.* (Smith et al. 2013). The four classes of putative enhancer elements are: Regions of FOXA1, FOXA2 or HNF4A binding that overlap H3K27ac and EP300 calls as well as at least one of three chromatin remodeling factors RAD21, CHD2 or SMC3 (type 1); Regions like in 1 but with no remodeling factor overlapping (type 2); EP300 peak regions overlapping H3K27ac as well as at least one of chromatin remodeling factor, but without peaks in FOXA1, FOXA2 or HNF4A (type 3); Regions like in 3 but with no remodeling factor overlapping (type 4). As shown here and in Fig. S11, we do not observe major differences between the four design types, either with respect to activity or MT vs. WT. (*B*+*C*) Enhancer activity of 200 synthetic regulatory element sequences (SRES) in the MT (*B*) and WT experiments (*C*). Scatter plot of RNA/DNA ratios for the top 100 positive and top 100 negative synthetic regulatory element (SRE) sequences in HepG2 experiments by Smith *et al.* (Smith et al. 2013). Plots show the combined RNA/DNA ratios on the y-axis and measurements by Smith *et al.* on the x-axis. Intervals indicate the mean, minimum and maximum values observed for three replicates performed with each experiment.

Importantly, we can see clear separation of positive and negative controls. Fig. 3B and 3C display RNA/DNA ratios obtained for the highest and lowest SRESs in the MT and WT experiments compared to their previously measured effects in HepG2. While the highest and lowest SRESs are well separated in both experiments (Kolmogorov-Smirnov and Wilcoxon Rank Sum p-values below 2.2E-16), the WT experiment separates the highest and lowest SRE controls slightly better than the MT experiment (Kolmogorov-Smirnov test D 0.97 vs 0.95, Wilcoxon Rank Sum test W 9951 vs 9937). Further, relative to the 90^th^ percentile of SRES negative controls in each experiment, a greater proportion of candidate enhancer sequences are active with integration (36% in WT vs. 28% in MT; Table S1).

We next sought to assess whether any of our design categories (i.e. types 1-4 defined above, reflecting subsets of candidate enhancers with coincident liver TF and/or chromatin remodeler ChIP-seq peaks) might underlie the observed differences. However, none of these design categories were meaningfully explanatory of enhancer activity or were predictive of differences between MT vs. WT (Figs. 3A and Fig S11).

### ENCODE and other genomic annotations that predict enhancer activity

Considering that our design categories were predictive of enhancer activity in neither episomal nor chromosomally based MPRA, nor of the differences between them, we explored whether other genomic annotations, some numerical and other categorical (Supplementary File 3), were predictive of our results in HepG2 cells. The performance of individual numerical annotations for predicting the observed activity of candidate enhancer sequences are shown in Fig. 4. We use Kendall’s tau, a non-parametric rank correlation that is more conservative than Spearman’s rho, because of the large number of zero-values in our annotations which can result in artifacts from ties with Spearman’s rho. In contrast with our design bins, many genomic annotations are observed to predict enhancer activity in both the WT and MT experiments. Across the board, annotations correlate better with the WT than the MT results, suggesting that integrated activity read-outs (WT) correlate better with endogenous functional genomic signals (e.g. ChIP-seq data) than do episomal activity read-outs (MT).

**Figure 4.**
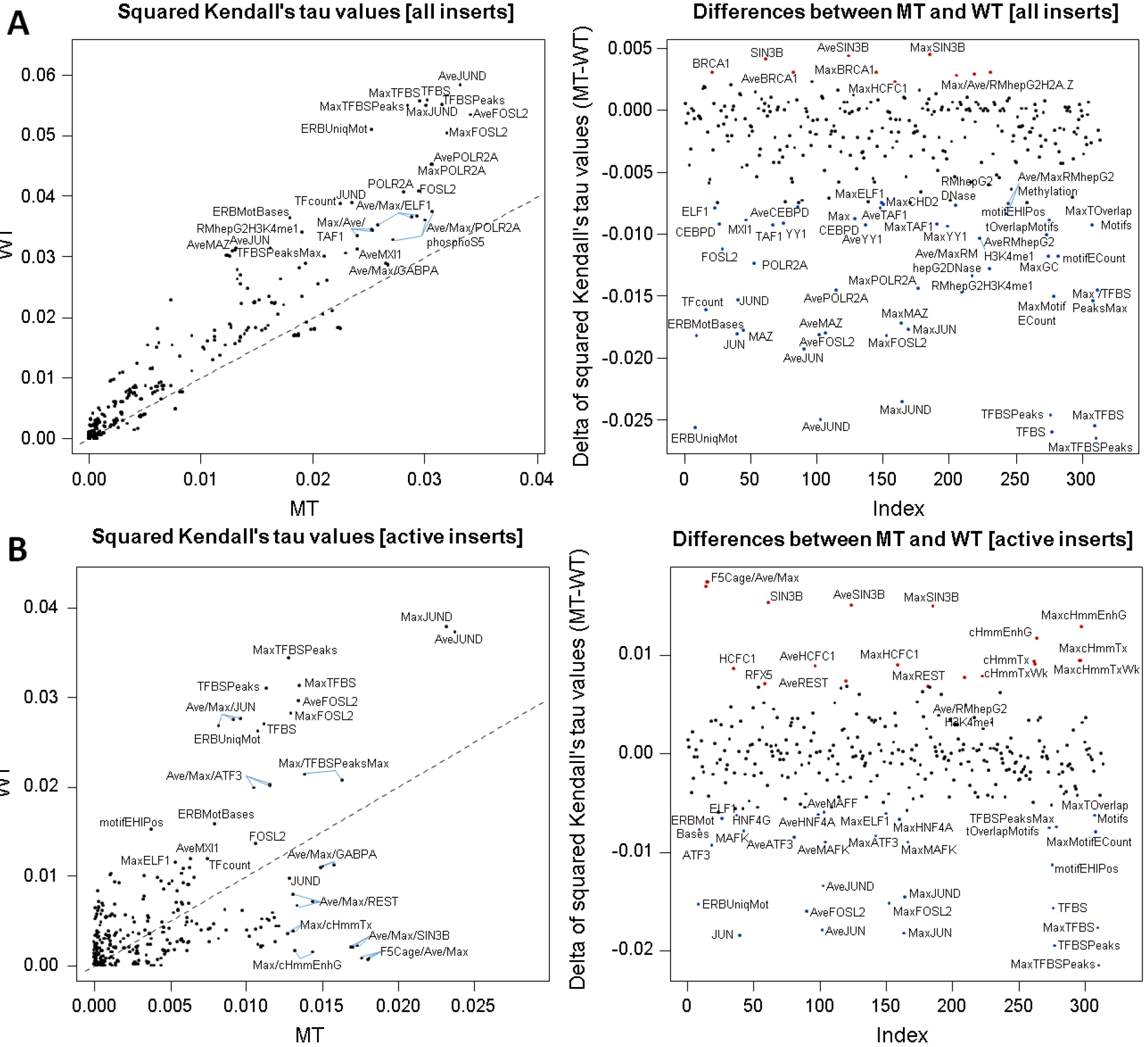
Squared Kendall’s tau (T^2^) values for available genome annotations for predicting the activity of candidate enhancer sequences in the non-integrating (MT) and integrating (WT) experiments. (*A*) WT RNA/DNA ratios correlate better with annotations than the respective MT values. The left panel highlights the top correlated annotations for WT and MT ratios. The right panel highlights annotations with the largest difference in T^2^ values between the MT and WT experiments. (*B*) Same analysis for the subset of active elements (Suppl. Table 1).

The most highly predictive numerical annotations, in both types of experiments, are HepG2 ChIP-seq datasets of JUND (Transcription Factor Jun-D) and FOSL2 (FOS-Like Antigen 2), consistent with a previous MPRA study which also highlighted the role of these transcription factors in HepG2 cells (Savic et al. 2015). For chromosomally based MPRA (WT), the number of overlapping ENCODE ChIP-seq peaks (TFBS) and the average ENCODE ChIP-seq signal (TFBSPeaks) as measured across different cell-lines also rank amongst the more highly predictive annotations. However, these same features are the most discrepant with MT; that is, substantially less predictive of episomal MPRA. Of note, the highest observed T^2^ for an individual annotation is only 0.034 (MT) and 0.058 (WT), leaving a large proportion of the variation in rank order unexplained and highlighting the need for a model combining annotations and other available information (see below).

We also analyzed how categorical annotations might predict the results of episomal and chromosomal enhancer reporter assays (Fig. S12-S15). Most of these annotations were derived for HepG2 cells by the ENCODE project. However, none of the cell-type specific categorical annotations (ChromHMM (Ernst and Kellis 2012), SegWay (Hoffman et al. 2012) and Open Chromatin annotation) were predictive of the measured RNA/DNA ratios. The multi-cell-type and higher resolution (25 vs 5 level) SegWay chromatin segmentation was most predictive of the measured RNA/DNA ratios. Here, sequences annotated as TSS (transcription start sites) exhibited the highest expression while sequences annotated as D (genomic death zones) exhibited the lowest expression. We note that potential promoters (defined as sites within 1kb of a TSS) comprise ~9% of all non-control sequences (208/2,236) and are enriched in type 3 (49/90) and type 4 (35/87) sequences. We also see the highest proportion of active sequences (where ‘active’ is defined relative to the 90^th^ percentile of SRES negative controls) in the type 3 and 4 categories, even when excluding promoters (Table S1; see also Fig. S11).

### Sequence-based predictors of functional activity

We next assessed the ability of sequence-based models to predict functional activity of our assayed sequences. Ghandi, Lee *et al.* (Ghandi et al. 2014) introduced a “gapped k-mer” approach for identifying active sequences in a specific cell-type from ENCODE ChIP-seq peaks and matched control sequences (gkm-SVM). The original publication trained models for individual binding factors from up to 5,000 ChIP-seq peaks and the same number of random control sequences. We collected all training data that Ghandi, Lee *et al.* used for HepG2, obtaining ~225,000 unique peak sequences as well as controls, and trained a combined, sequence-based model for predicting ChIP-seq peaks in HepG2 cells (see Methods). Based on a set-aside test dataset, the resulting model had a specificity of 71.8%, a sensitivity of 88.8% and a precision of 75.9% for separating ChIP-seq peak sequences from random control sequences.

We applied this model to our 171 bp candidate enhancer sequences, and asked how well the resulting gkm-SVM scores correlated with the RNA/DNA ratios obtained for the MT and WT experiments (Fig. 5A-B). The combined gkm-SVM HepG2 model results in a Spearman’s R^2^ of 0.082 and 0.128, for MT and WT respectively. This is comparable to the best results obtained for individual genomic annotations described before (MT Kendall’s T^2^ of 0.038 for gkm-SVM score vs 0.034 for the best individual annotation described before and WT Kendall’s T^2^ of 0.060 vs 0.058, respectively). However, we note that the correlation with the gkm-SVM model is at least partially driven by the synthetic control sequences, which can be scored with the sequence-based model but not with the genomic annotations. When excluding all control sequences, Spearman’s R^2^ values drop from 0.082 to 0.041 and from 0.128 to 0.076 for MT and WT, respectively. As such, there are a few ENCODE-based annotations which outperform the sequence-based gkm-SVM model, namely summaries of JUND/FOSL2 HepG2 ChIP-seq peaks, the number of overlapping ChIP-seq peaks (TFBS) or the average ChIP-seq signal (TFBSPeaks) measured across multiple ENCODE cell-types.

**Figure 5.**
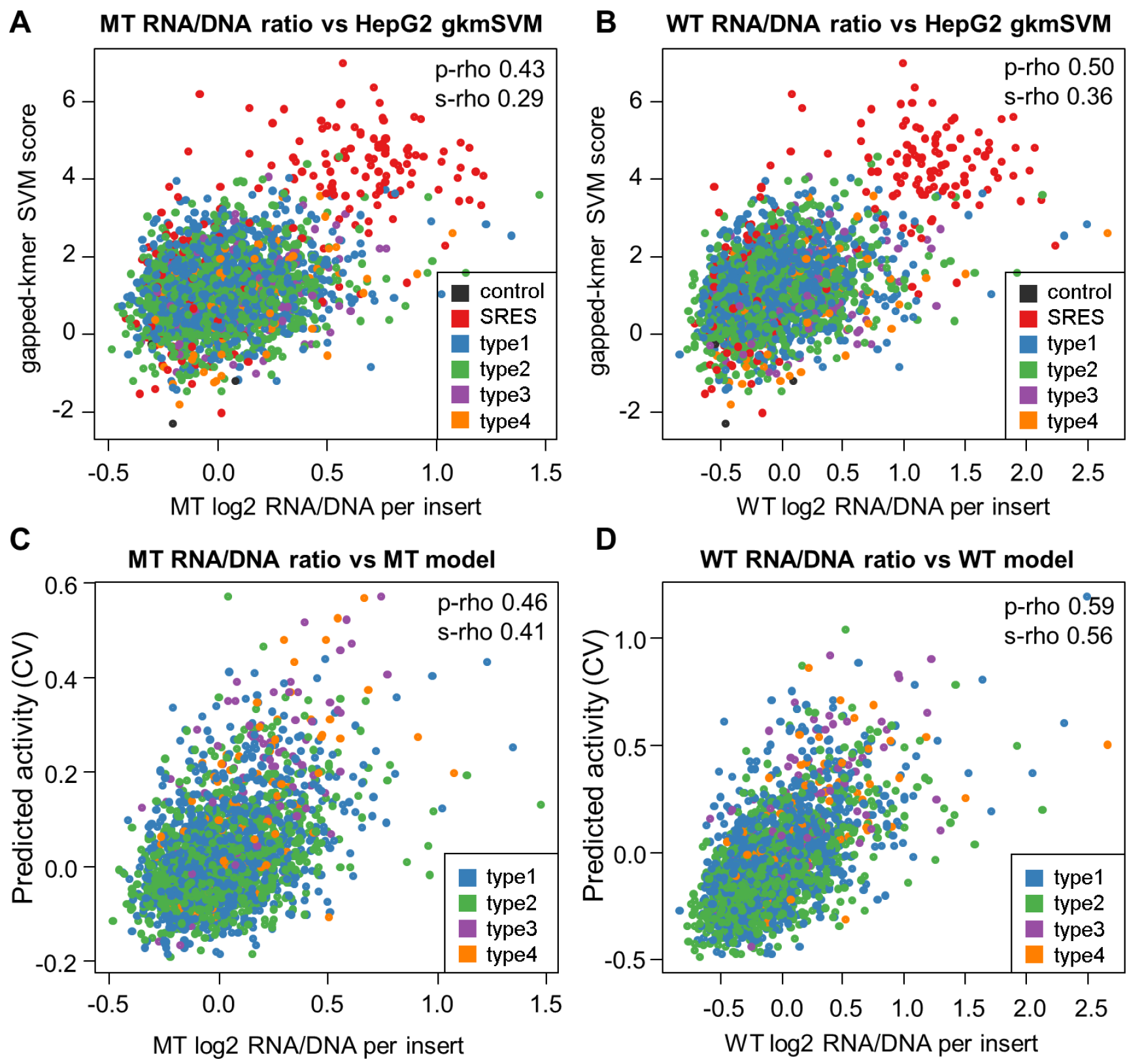
Prediction models. (*A*+*B*) Correlation of gkm-SVM scores obtained for a combined HepG2 model with RNA/DNA ratios obtained from the mutant (MT) and wild-type integrase (WT) experiments. Data points are colored by the type of insert sequence, including two types of controls, 200 synthetic regulatory element sequences (SRES, red) identified by Smith *et al.* (Smith et al. 2013) and four other control sequences (dark gray). The four classes of putative enhancer elements are: (type 1) regions of FOXA1, FOXA2 or HNF4A binding that overlap H3K27ac and EP300 calls as well as at least one of three chromatin remodeling factors RAD21, CHD2 or SMC3; (type 2) regions like in 1 but with no remodeling factor overlapping; (type 3) EP300 peak regions overlapping H3K27ac as well as at least one of chromatin remodeling factor, but without peaks in FOXA1, FOXA2 or HNF4A; (type 4) regions like in 3 but with no remodeling factor overlapping. Correlations are partially driven by the SRES; when excluding all controls, Spearman’s R^2^ values drop from 0.0817 to 0.0409 and from 0.1282 to 0.0756 for MT and WT, respectively. (*C*+*D*) Scatter plots of measured RNA/DNA ratios with predicted activity from linear Lasso models using annotations (numerical and categorical) as well as sequence-based (individual LS-GKM scores) information. Correlation coefficients are 0.46 Pearson / 0.41 Spearman for the non-integrated experiment (MT) and 0.59 Pearson / 0.56 Spearman for the integrated constructs (WT). The models selected 111 (MT) and 135 (WT) out of a total of 378 annotation features. Based on Pearson R^2^ values, these combined models explain 21.1% (MT) and 34.7% (WT) of the variance observed in these experiments.

### Combining annotations and sequence information to predict enhancer activity

We next sought to combine information across multiple annotations to better predict enhancer activity. We fit Lasso linear models and selected the Lasso tuning parameter value by crossvalidation (CV). Scatter plots as well as correlation coefficients were also obtained in a CV setup (see Methods). We built models with all the genomic annotations described above (including the categorical annotations as binary features) as well as with and without the sequence-based gkm-SVM score from scaled and centered annotation matrixes (Fig. S16-S18). SRESs and other controls were naturally excluded, as they are largely synthetic sequences and therefore missing genomic annotations. The resulting linear models were considerably more predictive of WT ratios than MT ratios (e.g. CV Spearman R^2^ of 0.272 WT vs. 0.146 MT; CV Pearson R^2^ of 0.307 WT vs. 0.193 MT). Including gapped-kmer SVM scores in the models improved performance further (CV Spearman R^2^ of 0.298 WT vs. 0.158 MT; CV Pearson R^2^ of 0.330 WT vs. 0.206 MT). We noticed that gapped-kmer SVM scores were assigned the largest model coefficients in both WT and MT models when they were included (Fig. S18). Thus, while reasonably performing models are obtained from genomic annotations, the sequence-based gkm-SVM scores appear to capture independently predictive information.

We therefore decided to further explore sequence-based models and turned to the faster and low-memory consumption LS-GKM implementation of gkm-SVM (Lee 2016). We trained models from each of the 64 narrow-peak ChIP-seq datasets for which we had included summary statistics for the annotation matrix above (see Methods). We then asked how well the LS-GKM scores generated by each of these 64 models predicted the results of the lentiMPRA experiments. Although the scores now correspond to sequence-based models of ChIP-seq peaks rather than the ChIP-seq peaks themselves, we once again observed the highest Spearman R^2^ values for the individual factors JUND (0.117 WT/0.055 MT) and FOSL2 (0.105 WT/0.053 MT), and these are also the factors that show the largest differences in predictive value for WT vs. MT (Fig. S19). As such, sequence-based models of binding by these two factors as well as other individual factors exceed the performance of the pooled gkm-SVM sequence model.

We fit Lasso linear models from the TF-specific LS-GKM SVM scores in order to predict the measured activities. The combined MT model (using 35 individual scores) achieves a CV Spearman R^2^ of 0.134 (CV Pearson R^2^ of 0.169), and the combined WT model (using 39 individual scores) of 0.231 (CV Pearson R^2^ of 0.263) (Fig. S20). This still falls short of models obtained purely from genomic annotations as described above. To test whether multiple ChIP-seq datasets should be combined in a sequence model rather than combining individual model scores in a linear model to improve prediction, we also trained LS-GKM models based on the peak sequences of the 35 (MT) and 39 (WT) scores selected by Lasso models as well as the top 5 and top 10 coefficients in the Lasso models for MT and WT. However, model performance only increased for combining small numbers of peak sets while combining all peaks in one sequence model reduces overall performance (Table S2).

Finally, when we used both genomic annotations and the individual LS-GKM scores in a single linear model to predict the measured activities, performance increased to a CV Spearman R^2^ of 0.171 (MT; Pearson R^2^ of 0.212) and 0.314 (WT; CV Pearson R^2^ of 0.347). These are our highest performing models predicting the activities of candidate enhancer sequences for both the episomally and chromosomally encoded MPRA experiments (Fig. 5C-D).

## Discussion

In this work, we report the first systematic comparison of episomal and chromosomally integrated reporter assays. Key aspects of our approach include: (1) Lentivirus-based MPRA or lentiMPRA, which can be used to in an episomal or integrated context by toggling whether a mutant vs. wild-type integrase is used, and can furthermore be used in a wide variety of cell types, including neurons; (2) The use of numerous barcodes per candidate enhancer sequence, which results in highly reproducible measurements of transcriptional activation; and (3) Extensive predictive modeling of our results, with the implicit assumption that a reasonable measure of a reporter assay’s biological relevance is the extent to which it is correlated with endogenous genomic annotations.

We find that the results of integrated reporter assays are more reproducible, robust and biologically relevant than episomal reporter assays. These conclusions are supported by the following observations: (1) We observed consistently greater reproducibility and dynamic range for the WT replicates as compared with the MT replicates. (2) The correlation of WT vs. MT replicates (Spearman correlation of 0.792) was substantially lower than for WT vs. WT (0.944) or MT vs. MT (0.908), with clear systematic differences between the integrated and episomal contexts that exceed technical noise (Fig. 2, Fig. S9). (3) The WT experiments were consistently more correlated with and more predictable by genomic annotations, which are based on biochemical marks measured in these sequences’ native genomic contexts. (4) Many genomic annotations significantly predict the results of the WT but not the MT experiments.

Of note, we observed generally higher levels of expression with integrated reporters (Fig. 3A and Table S1), consistent with previous findings that showed higher reporter gene levels for integrating relative to non-integrating HIV-1 (Gelderblom et al. 2008; Thierry et al. 2016). However, it is worth noting that we used a lentivirus (not HIV-1) with a self-inactivating (SIN) LTR, which lacks viral promoters or enhancers, potentially influencing these expression differences. Results from hydrodynamic tail vein assays (which delivers reporter constructs into the mouse liver) also show that when chromatinized plasmid DNA leads to higher expression levels than naked plasmid DNA (Kamiya et al. 2013). For HIV-1, both integrating and non-integrating HIV-1 viral DNA are associated with histones (Kantor et al. 2009), which is probably also the case for our lentiviral vector. However, even if the lentiviral episome is chromatinized, there remain myriad potential causes for the observed differences in expression, including differences in H1 stoichiometry, nucleosome positioning, cooperative TF binding (Hebbar and Archer 2007; Hebbar and Archer 2008), and/or nuclear location (Jeong and Stein 1994).

The number of overlapping ENCODE ChIP-seq peaks was one of the most strongly predictive annotations for our integrated sequences (Fig. 4, left). Interestingly, in experiments previously performed on the *MMTV* promoter it was observed that non-integrating constructs could not adequately assess cooperative TF binding due to differences in H1 stoichiometry and nucleosome positioning (Hebbar and Archer 2007), which may relate to the fact that this multiple TF binding was also one of the most differentiating annotations between the WT vs. MT experiments (Fig. 4, right). Specific TF ChIP-seq-based sequence models that are similarly differentiating (Fig. S19) include JUND, FOSL2, ATF3 and ELF1, which are known to interact and form complexes. Jun and Fos family members form the heterodimeric protein complex AP-1, which regulates gene expression in response to various stimuli including stress (Hess et al. 2004) and in the liver has known roles in hepatogenesis (Hilberg et al. 1993) and hepatocyte proliferation (Alcorn et al. 1990). AP-1 is known to form complexes with several additional protein partners (Hess et al. 2004), including ATF proteins such as ATF3 (Hai and Curran 1991) and ELF1, an ETS transcription factor (Bassuk and Leiden 1995). Of note, while lentivirus infection can induce stress potentially leading to increased expression of AP-1 and related factors, these same TFs were less predictive in MT-infected sequences, and furthermore the ChIP-seq datasets were generated on cells in normal physiological conditions. Combined, these findings suggest that differences in cooperative TF binding, possibly involving TFs including JUND, FOSL2, ELF1 and ATF3, might drive differences in the results of integrated vs. episomal reporter assays.

Using single-feature models, we also systematically evaluated more than 400 genomic annotations and sequence models to explore which are significantly predictive of expression in the integrated and/or episomal lentiMPRA experiments (Figure S21). Consistent with our other analyses, there are many more annotations that are significantly predictive of the integrated (WT) assay but not the episomal (MT) assay. These include several annotations related to histone acetylation (HDAC2, EP300, and ZBTB7A) as well as a factor with increased liver expression and which is associated with adipogenesis, gluconeogenic and hematopoiesis (CEBPB) (Tsukada et al. 2011). Interestingly, there are also a number of annotations which are only significantly predictive of the episomal assay, including BRCA1 ChIP-seq peaks and SIN3 transcriptional regulator family member B (SIN3B) binding motifs.

A contemporary challenge for our field is how to best identify, prioritize, and functionally validate *cis*-regulatory elements, especially enhancers. To address this, we envision a virtuous cycle, in which annotation and/or sequence-based models are used to nominate candidate enhancer sequences for validation, these candidates are tested in massively parallel reporter assays, and then the results are used to improve the models which in turn results in higher quality nominations. Eventually, this will lead to not only a catalog of validated enhancers but also a deeper mechanistic understanding of the relationship between primary sequence, transcription factor binding, and quantitative enhancer activity. In this study, our best performing model achieves a Pearson’s R^2^ of 0.347 in predicting the results of the integrated lentiMPRA, with both genomic annotations and sequence-based models providing independent information. Of note, these are quantitative predictions of activity, a more challenging task than simply categorizing enhancers vs. non-enhancers. Although far from perfect, we are able to garner insights into the determinants of enhancer function, and may be able to use this model to select a much larger number of candidate enhancer sequences for testing and further modeling.

As our field scales MPRAs to characterize very large numbers of candidate enhancers, it is obviously critical that the reporter assays are as reflective as possible of endogenous biology. Our results directly test a longstanding concern about episomal reporter assays, and suggest that there are substantial differences between the integrated and episomal contexts. Furthermore, based on the fact that their output is more correlated with genomic annotations, we infer that integrated reporter assays are more reflective of endogenous enhancer activity. This fits with our expectation, as both the integrated reporter and endogenous enhancers reside within chromosomes as opposed to episomes. We urge caution in the interpretation of the results of all reporter assays, and that integrated reporter assays such as lentiMPRA be used where possible.

## Methods

### Lentivirus enhancer construct generation

To generate the lentivirus vector (pLS-mP), a minimal promoter sequence, which originates from pGL4.23 (Promega), including an *SbfI* site was obtained by annealing of oligonucleotides (Sense: 5’ - CTAGA<u>CCTGCAGG</u>CACTAGAGGGTATATAATGGAAGCTCGACTTCCAGCTTGGCAATCCGGTACTGTA-3’, Antisense: 5’- CCGGTACAGTACCGGATTGCCAAGCTGGAAGTCGAGCTTCCATTATATACCCTCTAGTG<u>CCTGCAGG</u>T-3’ *SbfI* site is underlined), and subcloned into *XbaI* and *AgeI* sites in the pLB vector (Addgene 11619; (Kissler et al. 2006)) replacing the U6 promoter and CMV enhancer/promoter sequence in the vector. To generate pLS-mP-SV40, the SV40 enhancer sequence was amplified from pGL4.13 (Promega) using primers (Forward: 5’- CAGGGCCCGCTCTAGAGCGCAGCACCATGGCCTGAA-3’, Reverse: 5’- TGCCTGCAGGTCTAGACAGCCATGGGGCGGAGAATG-3’) and inserted into *XbaI* site in the vector using In-Fusion (Clontech). pos1, pos2, neg1, and neg2 sequences were amplified from human (pos1, neg2, pos2) or mouse (neg1) genome, and inserted into *EcoRV* and *HindIII* site in pGL4.23 (Promega). Primers used are shown in Table S3 and the annotated plasmid sequence file is available as Supplementary File 1.

### Library sequence design

We picked 171bp candidate enhancer sequences based on ChIP-seq peaks calls for HepG2. We used narrow peak calls for DNA binding proteins/transcription factors (FOXA1, FOXA2, HNF4A, RAD21, CHD2, SMC3 and EP300) and wide peak calls for histone marks (H3K27ac). We downloaded the call sets from the ENCODE portal (Sloan et al. 2016) (https://www.encodeproject.org/)with the following identifiers:ENCFF001SWK, ENCFF002CKI, ENCFF002CKJ, ENCFF002CKK, ENCFF002CKN, ENCFF002CKY, ENCFF002CUS, ENCFF002CTX, ENCFF002CUU, ENCFF002CKV, and ENCFF002CUN. We defined four classes of sites: 1) Regions centered over peak calls of ≤171bp for FOXA1, FOXA2 or HNF4A that overlap H3K27ac and EP300 calls as well as at least one of three chromatin remodeling factors RAD21, CHD2 or SMC3. 2) Regions like in 1, but with no remodeling factor overlap. 3) Regions of 171 bp centered in an EP300 peak overlapping H3K27ac as well as at least one of three chromatin remodeling factors RAD21, CHD2 or SMC3, but without peaks in FOXA1, FOXA2 or HNF4A. 4) Regions like in 3 but with no remodeling factor overlap. Sites of type 1 and 2 involving HNF4 were the most abundant sites and we used those to fill-up our design after exhausting other target sequences.

Potential 171bp target sequences were inserted into a 230bp-oligo backbone with a 5’-flanking sequence (15bp, AGGACCGGATCAACT), 14bp-spacer sequence (CCTGCAGGGAATTC), 15bp designed tag sequences (see below) and a 3’-flanking sequence (15bp, CATTGCGTGAACCGA) (Supplementary Fig. Y). Sequences were checked for *SbfI* and *EcoRI* restriction sites after joining the target sequence with the 5’-flanking sequence and the spacer sequence. Such potential target sequences were discarded.

In our final array design, we included 2,440 different target sequences each with 100 different barcodes (i.e. a total of 244,000 oligos). These included the highest 100 and lowest 100 synthetic regulatory element (SRE) sequences identified by Smith RP et al. (Smith et al. 2013), 4 control sequences (neg1 MGSCv37 chr19 35,531,983-35,532,154, neg2 GRCh37 chr5 172,177,151-172,177,323, pos1 GRCh37 chr3 197,439,136-197,439,306, pos2 GRCh37 chr19 35,531,984-35,532,154) which we tested using Luciferase assays in the HepG2 cell line (Fig. S2), 1,029 type 1 inserts (202 FOXA1, 180 FOXA2, 464 HNF4A, 120 FOXA1&FOXA2, 33 FOXA1&HNF4A, 17 FOXA2&HNF4A, 13 F OXA1&F OXA2&HNF 4A), 1,030 type 2 inserts (195 FOXA1, 174 FOXA2, 470 HNF4A, 126 FOXA1&FOXA2, 31 FOXA1&HNF4A, 20 FOXA2&HNF4A, 14 FOXA1&FOXA2&HNF4A), 90 type 3 inserts and 87 type 4 inserts.

Tag sequences of 15bp length were designed to have at least two substitutions and one 1bp-insertion distance to each other. Homopolymers of length 3 bp and longer were excluded in the design of these sequences, and so were ACA/CAC and GTG/TGT trinucleotides (bases excited with the same laser during Illumina sequencing). More than 556,000 such barcodes were designed using a greedy approach. The barcodes were then checked for the creation of *SbfI* and *EcoRI* restriction sites when adding the spacer and 3’-flanking sequences. From the remaining sequences, a random subset of 244,000 barcodes was chosen for the design. The final designed oligo sequences are available in Supplementary File 2.

### Generation of MPRA libraries

The lentiviral vector pLS-mP was cut with *SbfI* and *EcoRI* to temporarily liberate the minimal promoter and EGFP reporter gene. Array-synthesized 230bp oligos (Agilent Technologies) containing an enhancer, spacer, and barcode (Fig. S3) were amplified with adaptor primers (pLSmP-AG-f and pLSmP-AG-r) that have overhangs complementary to the cut vector backbone (Table S3), and the products were cloned using NEBuilder HiFi DNA Assembly mix (E2621). The adaptors were chosen to disrupt the original *SbfI* and *EcoRI* sites in the vector. The cloning reaction was transformed into electrocompetent cells (NEB C3020). Multiple transformations were pooled and midi-prepped (Chargeswitch Pro Filter Plasmid Midi Kit, Invitrogen CS31104). This library of cloned enhancers and barcodes was then cut using *SbfI* and *EcoRI* sites contained within the spacer, and the minimal promoter and *EGFP* that were removed earlier were reintroduced via a sticky end ligation (T4 DNA Ligase, NEB M0202) between the enhancer and its barcode. These finished vectors were transformed and midi-prepped as previously mentioned.

### Quality control of designed array oligos

Before inserting the minimal promoter and EGFP reporter gene, the plasmid library was sampled by high-throughput sequencing on an Illumina MiSeq (206/200+6 cycles) to check for the quality of the designed oligos and the representation of individual barcodes (sequencing primers are pLSmP-AG-seqR1, pLSmP-AG-seqIndx, and pLSmP-AG-seqR2; Table S3). We sequenced the target, spacer and tag sequences from both read ends and called a consensus sequence from the two reads. We obtained 19.2 million paired-end consensus sequences from this sequencing experiment. 52.6% of those sequences had the expected length, 26.1% of sequences were 1bp short and 8.9% were 2bp short (summing up to 87.6%). Only 1.6% of sequences showed an insertion of 1bp. These results are in line with expected dominance of small deletion errors in oligo synthesis. We aligned all consensus sequences back to all designed sequences using BWA MEM (Li and Durbin 2009) with parameters penalizing soft-clipping of alignment ends (-L 80). We consensus called reads aligning with the same outer alignment coordinates and SAM-format CIGAR string to reduce the effects of sequencing errors. We analyzed all those consensus sequences based on at least three sequences with a mapping quality above 0. We note that substitutions are removed in the consensus calling process if the correct sequence is the most abundant sequence. Among these 992,513 consensus sequences, we observe instances of 91% designed oligos and 78% of oligos with one instance matching the designed oligo perfectly. Across all consensus sequences the proportion of perfect oligos is only 19%, however the proportion vastly increases with the number of observations (69% at 20 counts, 99% at 40 counts; Table S4). These observations are in agreement with most molecular copies of an oligo being correct, in combination with high representation differences in the library. Supplementary Fig. S4A shows the distribution of alignment differences (as a proxy for synthesis errors) along the designed oligo sequences. Errors are distributed evenly along the designed insert sequence, with deletions dominating the observed differences. We observe that at some positions the deletion rate is reduced while the insertion rate is increased. We speculate that this might be due to certain sequence contexts.

### Limited coverage of designed oligos in MPRA libraries

From the analysis of oligo quality and oligo abundance above, we saw a first indication of the existence of a wide range of oligo abundance and that more frequent sequences tend to match the designed sequences perfectly (see Table S4). We characterized the abundance of oligos further and looked at the consequences that this has for generating libraries of lentivirus constructs with limited complexity (due to the transformation of a limited number of bacteria). Rather than looking at full length oligos, we focused only on the tag sequences. Tag sequences were identified from the respective alignment positions of the alignments created above. To match the RNA/DNA count data analysis (see below), we only considered barcodes of 15bp length (10.96M/57.0%, similar to the proportion of correct length sequences above). Of those 10.96M barcodes, 345,247 different sequences are observed. We clustered (dnaclust (Ghodsi et al. 2011)) the remaining sequences allowing for one substitution and selecting the designed or most abundant sequence (reducing to 238,206 different sequences). The clustered sequences were matched against the designed barcodes (217,176 sequences, 99.2% of counts). The distribution of the abundance of these barcodes is available in Supplementary Fig. S4B. We used those counts to simulate sampling from this over dispersed pool of sequences, as done when taking a sample of plasmids infusing the reporter gene and minimal promoter and again transforming the resulting plasmids. We sampled 10 times and averaged the number of unique designed barcodes: 150k clones – 87,944 unique barcodes, 250k clones – 116,297 unique barcodes, 350k clones – 135,222 unique barcodes, 500k clones – 154,090 unique barcodes, 600k clones – 163,831 unique barcodes, 750k clones – 172,770 unique barcodes and 1M clones – 183,685 unique barcodes. Thus, even for high sampling depth only a subset of barcodes will be captured in the final library. We observe on average 145,876 different barcodes which is concordant with more than 430k clones going into the lenti construction.

### Cell culture and GFP / luciferase assays

HepG2 cells were cultured as previously described (Smith et al. 2013). K562, H1-ESC, HeLa-S3, T-47D and Sk-n-sh cells were culture as previously described (Dunham I et al. 2012). Sk-n-sh cells were treated with 24 uM *all trans*-retinoic acid (Sigma) to induce neuronal differentiation. K562, H1-ESC, HeLa-S3, T-47D and Sk-n-sh were treated with retinoic acid and infected with pLS-mP or pLS-SV40-mP lentivirus along with 8 μg/ml polybrene, and incubated for 2 days, when they have an estimated 30, 60, 90, 90, and 90 viral particles/cell, respectively. The number of viral particles/cell was measured as described below. For the four control sequences (two negatives and two positives) luciferase assay, we amplified the controls from the designed oligo pool (primer sequences available in Table S5) and inserted those into the pGL4.23 (Promega) reporter plasmid. 2×10^4^ HepG2 cells/well were seeded in a 96-well plate. 24 hour later, the cells were transfected with 90 ng of reporter plasmids (pGL4.23-neg1, pGL4.23-neg2, pGL4.23-pos1, and pGL4.23- pos2) and 10 ng of pGL4.74 (Promega), which constitutively expresses Renilla luciferase, using X-tremeGENE HP (Roche) according to the manufacturer’s protocol. The X-tremeGENE:DNA ratio was 2:1. Three independent replicate cultures were transfected. Firefly and Renilla luciferase activities were measured as previously described (Smith et al. 2013).

### Lentivirus packaging, titration and infection

Twelve million HEK293T cells were plated in 15 cm dish and cultured for 24 hours. The cells were co-transfected with 8 μg of the liver enhancer library and 4 μg of packaging vectors using jetPRIME (Polyplus-transfections). psPAX2 that encodes wild-type *pol* was used to generate integrating lentivirus, while pLV-HELP (InvivoGen) that encodes a mutant *pol* was used to generate non-integrating lentivirus. pMD2.G was used for both. The transfected cells were cultured for 3 days and lentivirus were harvested and concentrated as previously described (Wang and McManus 2009).

To measure DNA titer for the integrating and non-integrating lentivirus library, HepG2 cells were plated at 2×10^5^ cells/well in 12-well plates and incubated for 24 hours. Serial volume (0, 1, 5, 25 uL) of the lentivirus was added with 8 μg/ml polybrene, to increase infection efficiency. The infected cells were cultured for 2-5 days and then washed with PBS three times. Genomic DNA was extracted using the Wizard SV genomic DNA purification kit (Promega). Copy number of viral particle per cell was measured as relative amount of viral DNA (WPRE region) over that of genomic DNA (intronic region of *LIPC* gene) by qPCR using SsoFast EvaGreen Supermix (BioRad), according to manufacturer’s protocol. PCR primer sequences are shown in Table S3. For the lentiMPRA, 2.4 million HepG2 cells were plated on a 10 cm dish and cultured for 24 hours. The cells were infected with integrating or non-integrating lentivirus libraries along with 8 μg/ml polybrene, and incubated for 4 and 3 days, when they have an estimated 50 and 100 viral particles/cell, respectively. Three independent replicate cultures were infected. The cells were washed with PBS three times, and genomic DNA and total RNA was extracted using AllPrep DNA/RNA mini kit (Qiagen). Copy number of viral particle per cell was confirmed by qPCR and shown in Supplementary Fig. S5B. Messenger RNA (mRNA) was purified from the total RNA using Oligotex mRNA mini kit (Qiagen) and treated with Turbo DNase to remove contaminating DNA.

### RT-PCR, amplification and sequencing of RNA/DNA

For each replicate, 3x500ng was reverse transcribed with SuperscriptII (Invitrogen 18064-014) using a primer downstream of the barcode (pLSmP-ass-R-i#, Table S3), which contained a sample index and a P7 Illumina adaptor sequence. The resulting cDNA was pooled and split into 24 reactions, amplified with Kapa Robust polymerase for 3 cycles using this same reverse primer paired with a forward primer complementary to the 3’ end of EGFP with a P5 adaptor sequence (BARCODE_lentiF_v4.1, Table S3). The implemented two-round PCR set-up is supposed to reduce PCR jack-potting effects and allows for incorporating unique molecular identifiers (UMIs), which could be used to correct for other PCR biases in future experiments. PCR products are then cleaned up with AMPure XP beads (Beckman Coulter) to remove the primers and concentrate the products. These products underwent a second round of amplification in 8 reactions per replicate for 15 cycles, with a reverse primer containing only P7. All reactions were pooled at this point, run on an agarose gel for size-selection, and submitted for sequencing. For the DNA, 16x500ng of each replicate was amplified for 3 cycles just as the RNA. First round products were cleaned up with AMPure XP beads, and amplified for another 16 reactions, each for 20 cycles. Reactions were pooled, gel-purified, and sequenced. Sequencing primers are BARCODE-SEQ-R1-V4, pLSmP-AG-seqIndx, and BARCODE-SEQ-R2-V4 for both RNA and DNA barcodes (Table S3).

RNA and DNA for each of three replicates was sequenced on an Illumina NextSeq instrument (2×26 + 10bp index). The forward and reverse reads on this run each sequenced the designed 15bp barcodes as well adjacent sequence to correctly trim and consensus call barcodes. We obtained a minimum of 2.9M and a maximum of 5.9M raw counts for DNA (average 4.1M) and a minimum of 20.0M and a maximum of 32.3M raw counts for RNA (average 25.6M). Across replicates and sample type, 97% of barcodes were of the correct length of 15bp.

The number of unique sequences was on average 446k for DNA and 1.2M for RNA. When clustering sequences with one substitution (dnaclust; (Ghodsi et al. 2011)), the average number of unique sequences reduced to 280k for DNA and 697k for RNA. When overlapping the observed with the designed sequences, clustering keeps more counts but reduces the total number of observed barcodes (93.1% vs. 90.3%, 145k vs 151k). We believe this is due to too many errors in barcodes which are sufficiently similar to cause clusters to merge across different designed tag sequences. We therefore dismissed the clustered data and only matched against the designed barcodes. This is further supported by counts being more highly correlated between replicates when using the non-clustered data (Spearman’s rho without clustering: DNA replicates 88.6%, RNA replicates 98.0%; with clustering: DNA replicates 85.0%, RNA replicates 94.3%).

### Replicates, normalization and RNA/DNA ratios

To normalize RNA and DNA for different sequencing depths in each sample, we divided reads by the sum of observed counts and reported them as counts per million. Only barcodes observed in RNA and DNA of the same sample were considered. Subsequently, RNA/DNA ratios were calculated. We observe that the dynamic range observed in the WT experiments is larger and that the average Spearman’s rho is also higher for the WT experiments (44.3% vs. 39.0%). To determine the RNA/DNA ratios per insert, we summed up the counts of all barcodes contributing and determined the ratio of the average normalized counts. We explored how stable the correlation of RNA/DNA ratios is between replicates when limiting the number of barcodes per insert (Fig S10). We limited the maximum number of barcodes considered by (1) randomly down sampling and (2) requiring an exact number of barcodes per insert (i.e. down sampling those with more and excluding those inserts with less barcodes).

Even though normalized individually, the three replicates of each experiment do not seem to be on the exact same scale (Figure 2 & S9). We therefore chose to divide the RNA/DNA ratios by the median across the technical replicate value before averaging them.

### Predictors of sequence effects

To correlate available annotations with the observed sequence activity in HepG2 cells, we downloaded additional narrow peak calls for DNA binding proteins/transcription factors in HepG2 from ENCODE data. We obtained call-sets for the following 64 factors: ARID3A, ATF3, BHLHE40, BRCA1, CBX1, CEBPB, CEBPD, CHD2, CTCF, ELF1, EP300, EZH2, FOSL2, FOXA1, FOXA2, FOXK2, GABPA, GATA4, HCFC1, HDAC2, HNF4A, HNF4G, IRF3, JUN, JUND, MAFF, MAFK, MAX, MAZ, MBD4, MXI1, MYBL2, MYC, NFIC, NR2C2, NRF1, POLR2A, POLR2AphosphoS2, POLR2AphosphoS5, RAD21, RCOR1, REST, RFX5, RXRA, SIN3A, SIN3B, SMC3, SP1, SP2, SRF, TAF1, TBP, TCF12, TCF7L2, TEAD4, TFAP4, USF1, USF2, YY1, ZBTB33, ZBTB7A, ZHX2, ZKSCAN1, and ZNF274. Additionally, we downloaded ChromHMM segmentations for HepG2, Open Chromatin State, SegWay, and DHS call-sets from the ENCODE portal (Sloan et al. 2016). From the NIH Roadmap Epigenomics Consortium we obtained RNAseq, DNA methylation, DNase, CAGE, H2A.Z, H3K4me1, H3K4me2, H3K4me3, H3K9ac, H3K9me3, H3K27ac, H3K27me3, H3K36me3, H3K79me2, and H4K20me1 tracks (Kundaje et al. 2015). We also downloaded the Fantom5 Robust Enhancer annotations (Andersson et al. 2014), Fantom5 CAGE data for HepG2 (Forrest et al. 2014), GenoSTAN Enhancer and promoter predictions (http://i12g-gagneurweb.in.tum.de/public/paper/GenoSTAN/), enhancerFinder predictions (Erwin et al. 2014), as well as motif scan results and annotated regulatory features from the Ensembl Regulatory Build (Zerbino et al. 2015). For further genome-wide and organismal metrics, we turned to the CADD v1.3 annotation file (Kircher et al. 2014) and extracted local GC and CpG content, SegWay, chromHMM state across the NIH RoadMap cell types, priPhCons, mamPhCons, verPhCons, priPhyloP, mamPhyloP, verPhyloP, GerpN, GerpS, GerpRS, bStatistic, tOverlapMotifs, motifECount, motifEHIPos, TFBS, TFBSPeaks, TFBSPeaksMax, distance to TSS, and the actual CADD score column. We included the number of bases covered by peak calls as well as the average and maximum values across the designed sequences for those metrics. Supplementary File 3 outlines all annotations used.

### Gapped-kmer SVM (gkm-SVM) model of HepG2 activity

We collected training data of individual ChIP-seq binding factors described by M Ghandi & D Lee et al. (Ghandi et al. 2014) for HepG2 (5k specific ChIP-seq peak regions and the same number of random controls, http://www.beerlab.org/gkmsvm/) and removed duplicate sequences, obtaining 225k peak sequences as well as matched random controls. Attempting to train a classification model with the gkm-SVM software based on all peak sequences exceeded reasonable memory requirements (>1TB). Therefore, we iteratively reduced the number of training examples and ended up sampling each 50k peak and 50k control sequences for a combined HepG2 sequence model. Based on a test data set (2k sampled from the unused training data set), the obtained model has a specificity of 71.8%, a sensitivity of 88.8% and precision of 75.9% for separating ChIP-seq peak from the random control sequences.

### Linear models integrating individual annotations

We used the R glmnet package to fit Lasso-penalized linear models to predict RNA/DNA ratios. We used 10-fold cross-validation (cv.glmnet) to determine the Lasso tuning parameter lambda resulting in the minimum squared error. The Lasso forces small coefficients to zero, and thereby performs regression and feature selection simultaneously. Categorical features with K levels were included as K-1 binary columns. We excluded ZNF274 and EZH2 annotations from the model as none of the inserts overlapped with these ChIP-seq tracks. Otherwise missing annotation values were mostly in count features (70.1%) or absence of the conserved block annotation “GerpRS” (27.5%) and thus all these values were imputed to zero. All annotation features were scaled and centered. To report unbiased correlation values and scatter plots between the true and predicted RNA/DNA ratios, we randomly split up our data into 10 folds, trained models using 9 folds and the above identified tuning parameter and then extracted the fitted values after applying the model to the remaining fold.

### Sequence-based LS-GKM models

LS-GKM (Lee 2016) is a faster and lower memory profile version of gkm-SVM. Its default settings are different from gkm-SVM (e.g. using 11 bases with 7 informative positions rather than 10 bases with 6 informative positions). We applied LS-GKM using parameters corresponding to gkm-SVM (-l 10 -k6 -d3 -t2 -T4 -e0.01) as well as default parameters (-T4 -e 0.01) on the HepG2 training data described for gkm-SVM above (225,327 positive/negative sequences each, 10,000 kept set aside for validation). We also compared performance for using the negative sequences as described for gkm-SVM (Ghandi et al. 2014) versus obtaining negative sequences by permutation of the real sequences maintaining dinucleotide content (Jiang et al. 2008). We found that best results were obtained for LS-GKM defaults in combination with selected negative sequences rather than permuted sequences (Table S6). However, permuted sequences as negative set produced a higher true positive rate and substantially simplify computation. We therefore used permuted sequence sets and ran LS-GKM with default parameters for all models. We extracted genomic sequences (GRCh37) below the 64 ChIPseq peak sets by concatenating multiple call-sets for the same factor and merging overlapping peak regions using bedtools (Quinlan and Hall 2010). We extracted up to 1kb of sequence for each peak, or centered 1kb fragments on the peak for larger peak calls. We chose the model convergence parameter *e* based on the number of positive training sequences (mean 16600, min. 186, max. 63948) multiplied with 1E-07; investing more training iterations for smaller training data sets.

We then used Lasso regression (as described above) to create combined models and we also trained LS-GKM models from pooled peak data sets (Table S2). For this purpose, we pooled sequences using the peak data sets underlying the top 5, top 10 and all sequence models selected using Lasso regression.

### Individual feature models

To explore whether certain annotations are more strongly predictive for either the non-integrated (MT) or integrated (WT) expression measurements (despite the correlations among the annotations), we used the R glm (Generalized Linear Models) implementation to fit 430 linear single coefficient plus intercept models for predicting log2 RNA/DNA ratios for MT and WT experiments. We report the two-sided p-value for the t-statistic corresponding to the coefficient in the linear model, and used a significance criterion of 0.05 after Bonferroni correction (Figure S21, Table S7).

## Data access

Raw sequencing data, designed oligo sequences and processed count and RNA/DNA ratio data including annotations was submitted to the NCBI Gene Expression Omnibus (GEO) and was assigned accession <u>GSE83894</u>.

## Acknowledgements

We thank members of the Ahituv, McManus and Shendure laboratories for helpful discussions and suggestions. Our work was supported by the National Human Genome Research Institute (NHGRI) grant number 1R01HG006768 (NA & JS), NHGRI and National Cancer Institute grant number 1R01CA197139 (GC, DW, NA, JS) and National Institute of Mental Health grant number 1R01MH109907 (NA). JS is an investigator of the Howard Hughes Medical Institute.

### Author’s contributions

FI, MK, BM, MTM, NA and JS designed experiments; FIand BM performed all wet lab experiments; MK, JS, NA, DMW and GMC outlined data analysis; MK performed data analysis; FI, MK, NA, JS interpreted the experimental results; FI, MK, BM, NA and JS wrote the manuscript.

## Supplemental Information

### Supplementary figures

**Supplemental Figure S1.**
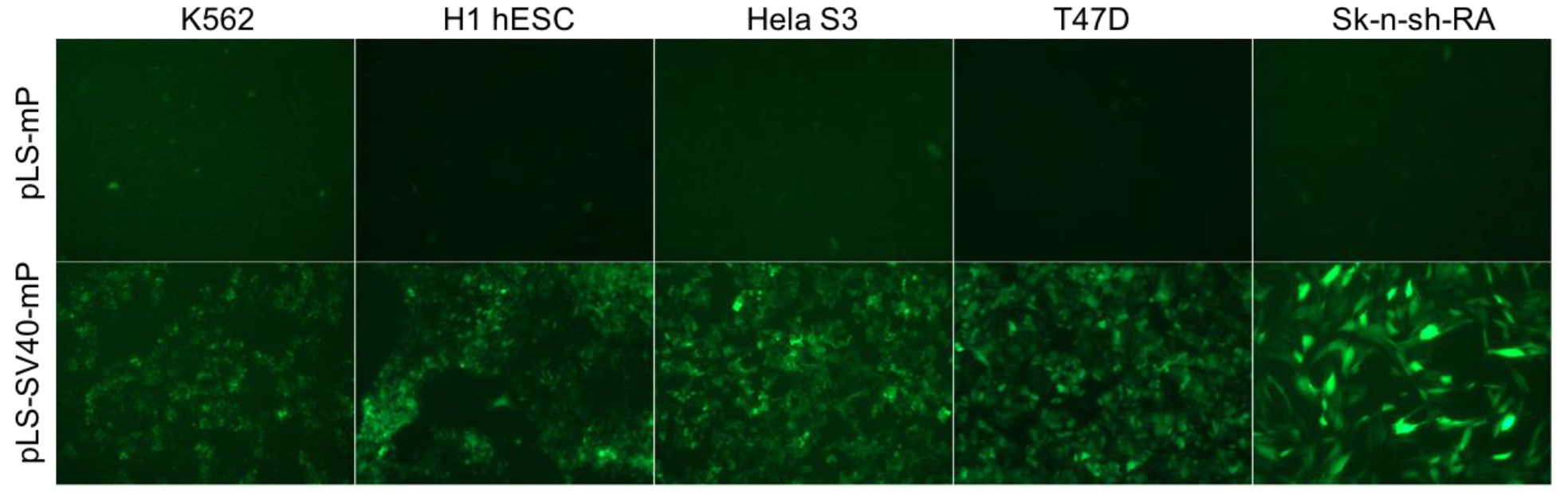
Validating lentiviral infection in several different cell lines. pLS-mP or pLS-SV40-mP infection results following 48 hours for K562, H1 human embryonic stem cells (H1 hESC), HeLa S3, T47D, and Sk-n-sh cells treated with retinoic acid (Sk-n-sh-RA).

**Supplemental Figure S2.**
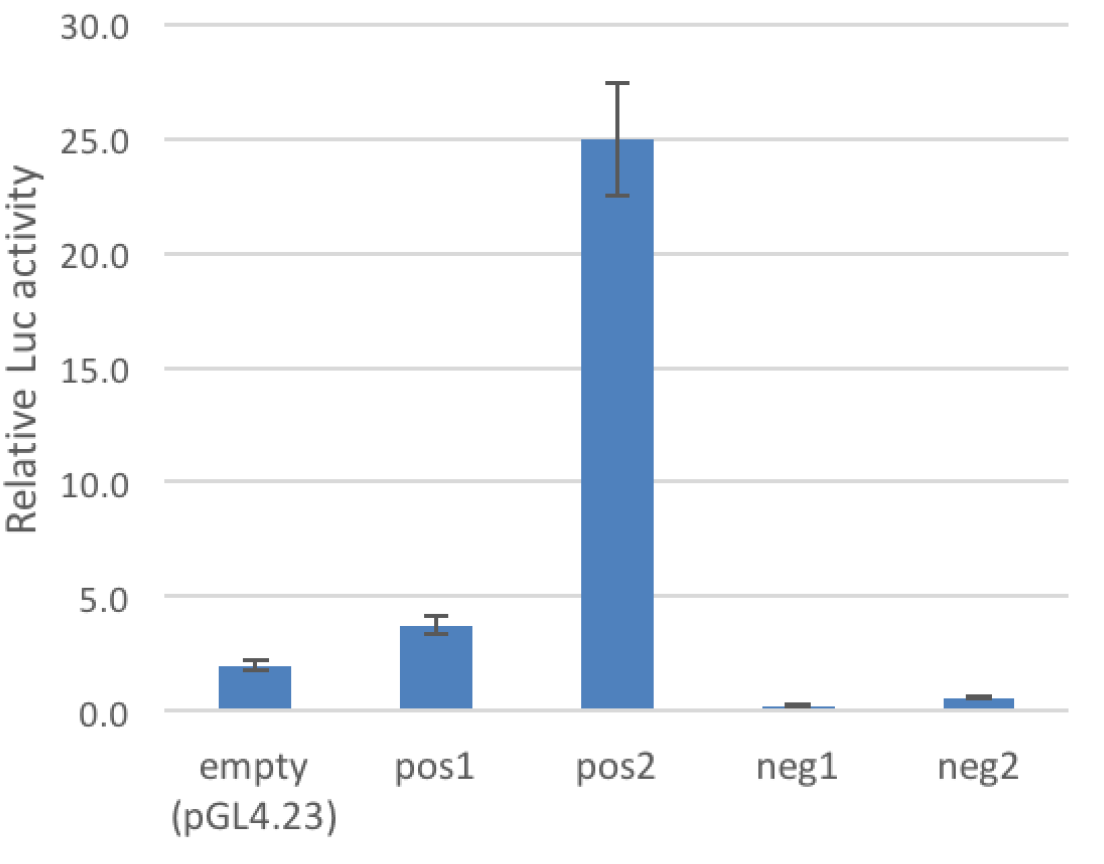
Luciferase assay for two positive (pos1 and pos2) and two negative (neg1 and neg2) control sequences. The reporter vectors and the empty vector (pGL4.23) were transfected into HepG2 cells, and the enhancer activity was measured as relative luciferase activity compared to Renilla luciferase expression following 24 hours.

**Supplemental Figure S3.**
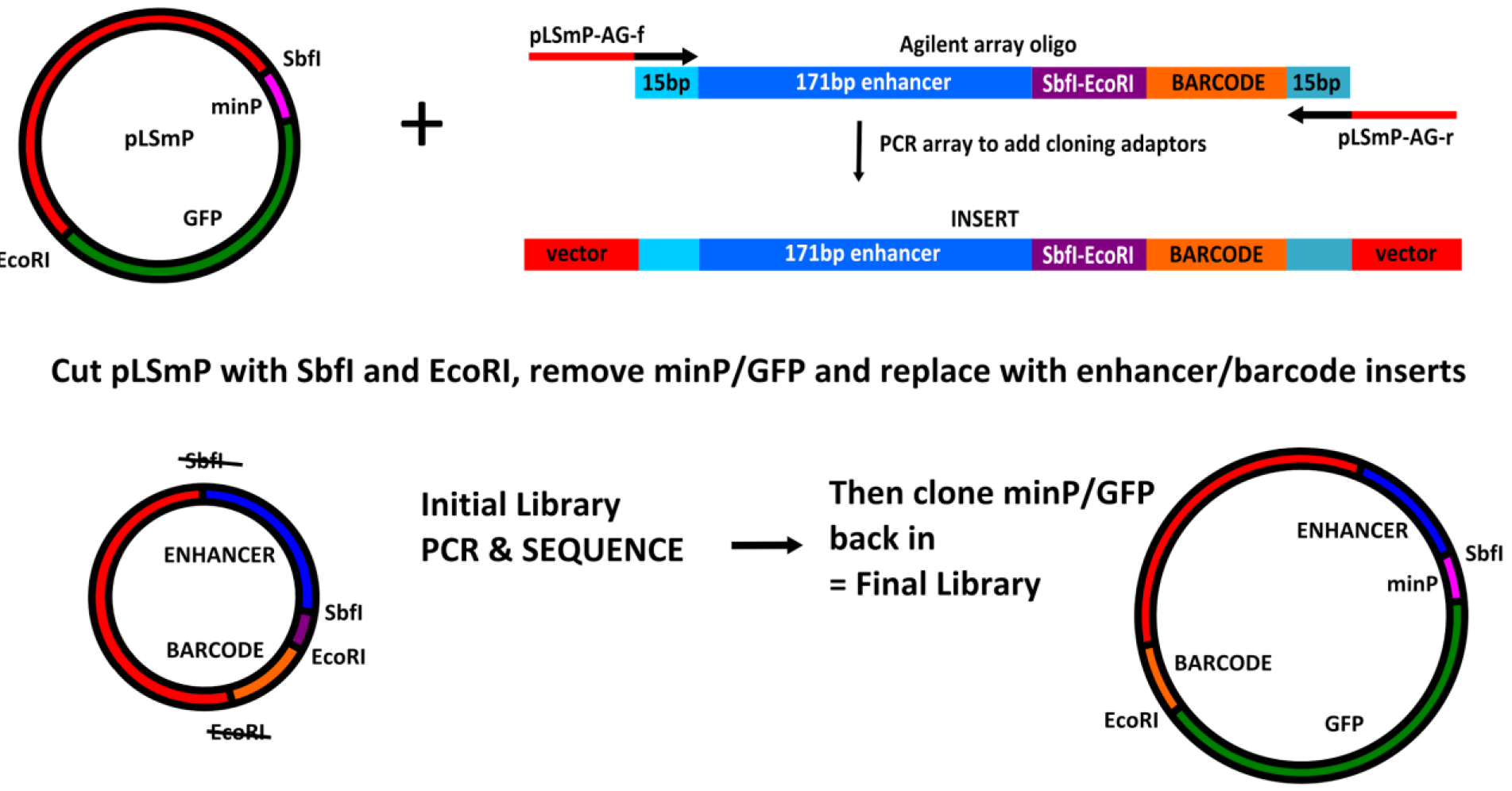
MPRA library cloning strategy. GFP, green fluorescent protein, minP, minimal promoter; *SbfI*, *EcoR1* restriction sites. The lenti vector pLS-mP is cut with *SbfI* and *EcoRI* to remove the minimal promoter and GFP. The enhancer/barcode agilent array is amplified with adaptor primers, and these PCR products are cloned into the pLS-mP backbone using NEBuilder HiFi Assembly mix. Cloning disrupts the original *SbfI* and *EcoRI* sites. This initial library can be sequenced to validate its accuracy and complexity. The library is then digested with *SbfI* and *EcoRI* to re-insert the minimal promoter and GFP between the enhancer and the barcode with a sticky-end ligation.

**Supplemental Figure S4.**
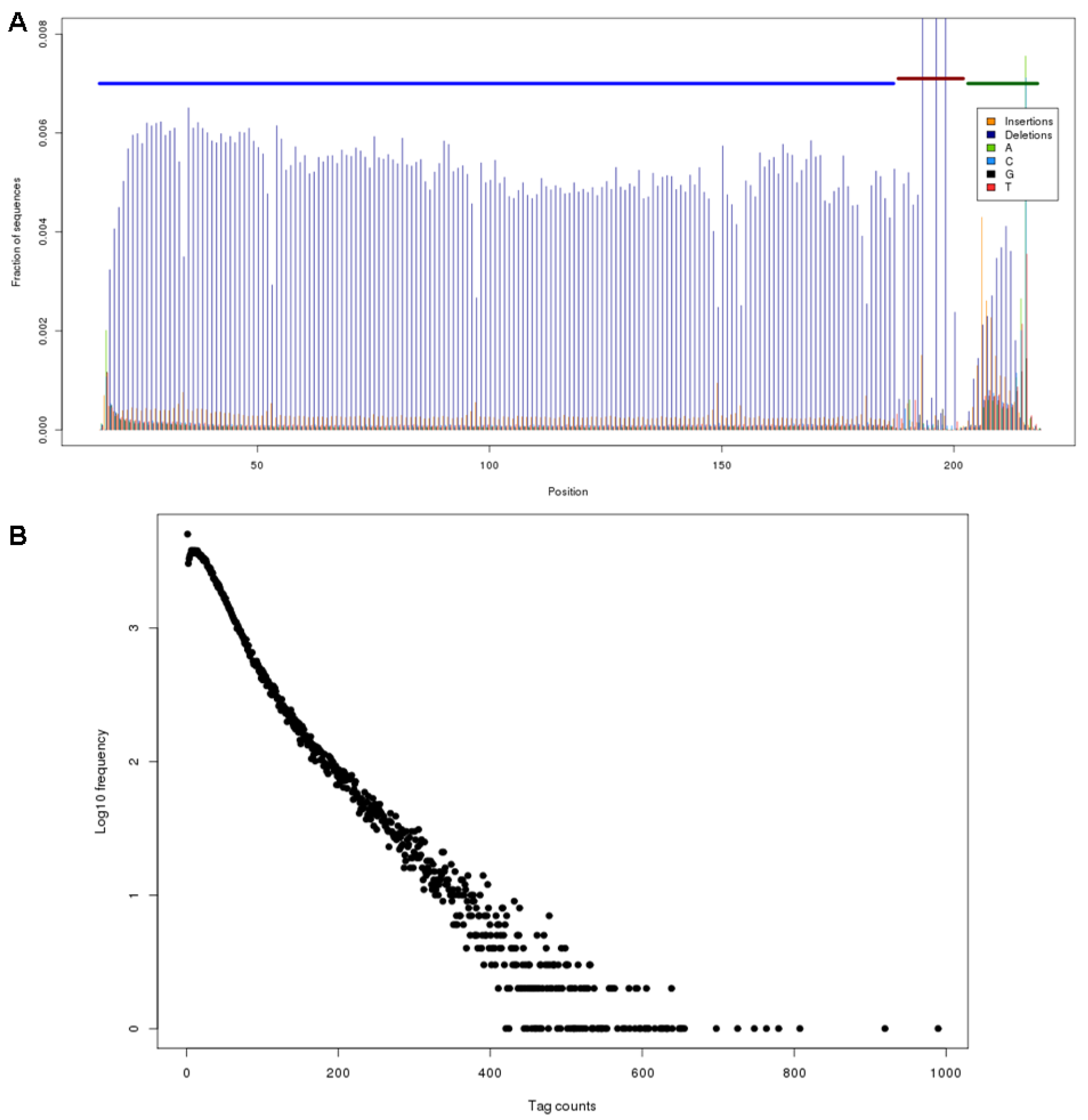
Sequence quality and barcode representation in the array-synthesized lentiMPRA library. *(A)* Positioning of errors/differences to the designed oligo sequences as observed from consensus-called BWA alignments. Thick horizontal lines near the top indicate the different oligo parts (insert - blue, spacer - red, barcode - green). Deletions towards read ends are misrepresented in this plot as those that result in altered outer alignment coordinates rather than the identification of a deletion. Filtering for sequence mapability results in artifacts around the barcode sequence which is required to disambiguate oligos. (*B*) Number of times designed reporter barcodes are observed in the oligo validation sequencing experiment. The frequency axis (y) has been log-transformed to show an over dispersion effect in the library, where a minority of barcodes contribute many of the observations. This effect can be observed from a change in gradient between barcodes observed below 150 times and barcodes observed more than 150 times. The 11,889 barcodes (5.5%) observed with more than 150 observations account for 24.5% of all observations.

**Supplemental Figure S5.**
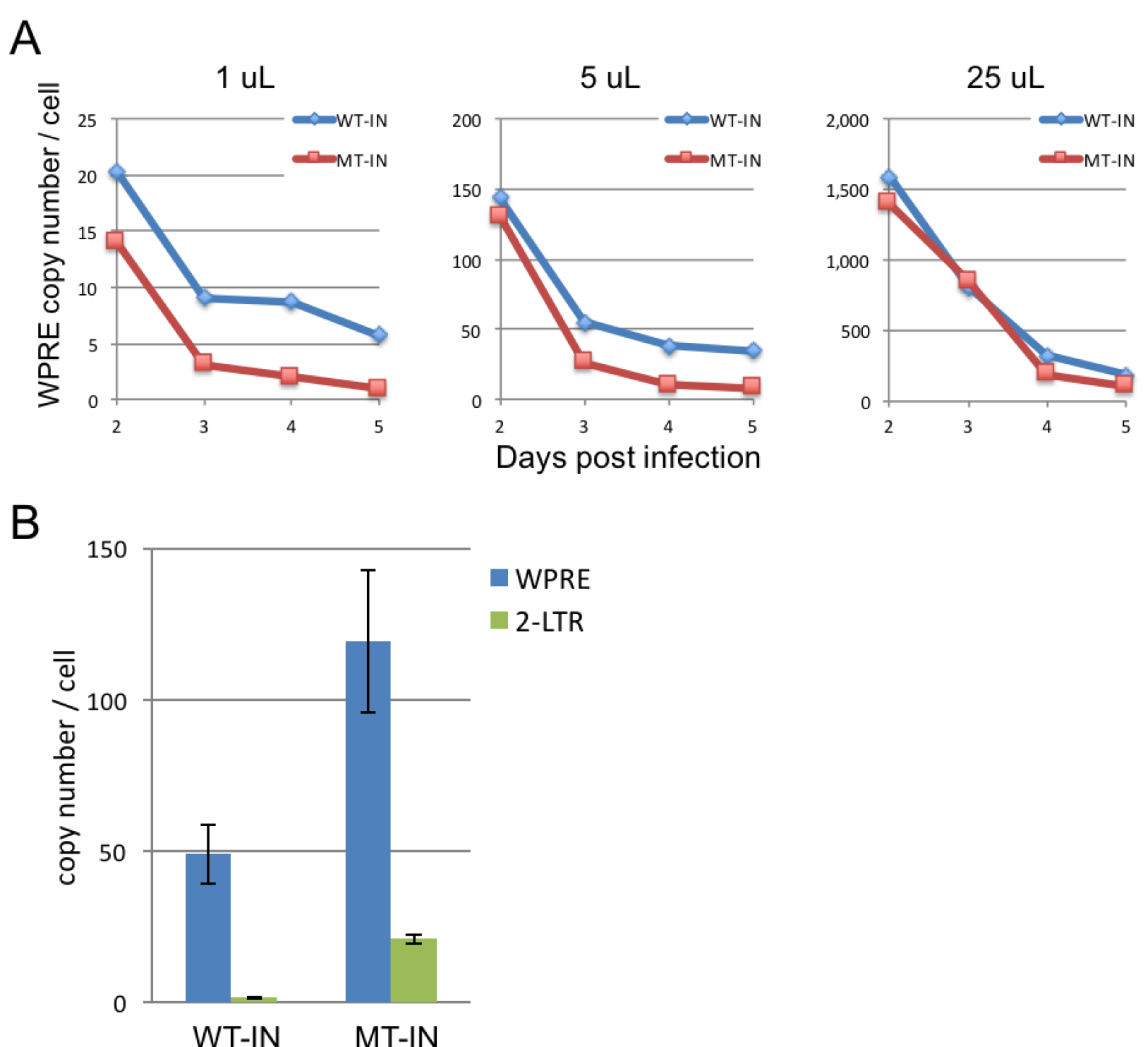
Analysis of wild-type integrase (WT-IN) and mutant integrase (MT-IN) virus infection conditions. (*A*) DNA titer for pLS-SV40-mP using either a WT-IN and MT-IN virus as determined by qPCR with primers against WPRE compared to genomic primers for the intronic region of the *LIPC* gene. Three different volumes (1, 5, and 25 μl per well of a 24-well plate) of lentivirus were analyzed at days 2-5 days. (*B*) Analysis of total viral DNA (WPRE) and unintegrated viral DNA (2-LTR circular DNA) for WT-IN library 4 days after infection and MT-IN library 3 days after infection using qPCR. The relative amount of viral DNA was measured using primers that target an intronic region of the *LIPC* gene.

**Supplemental Figure S6.**
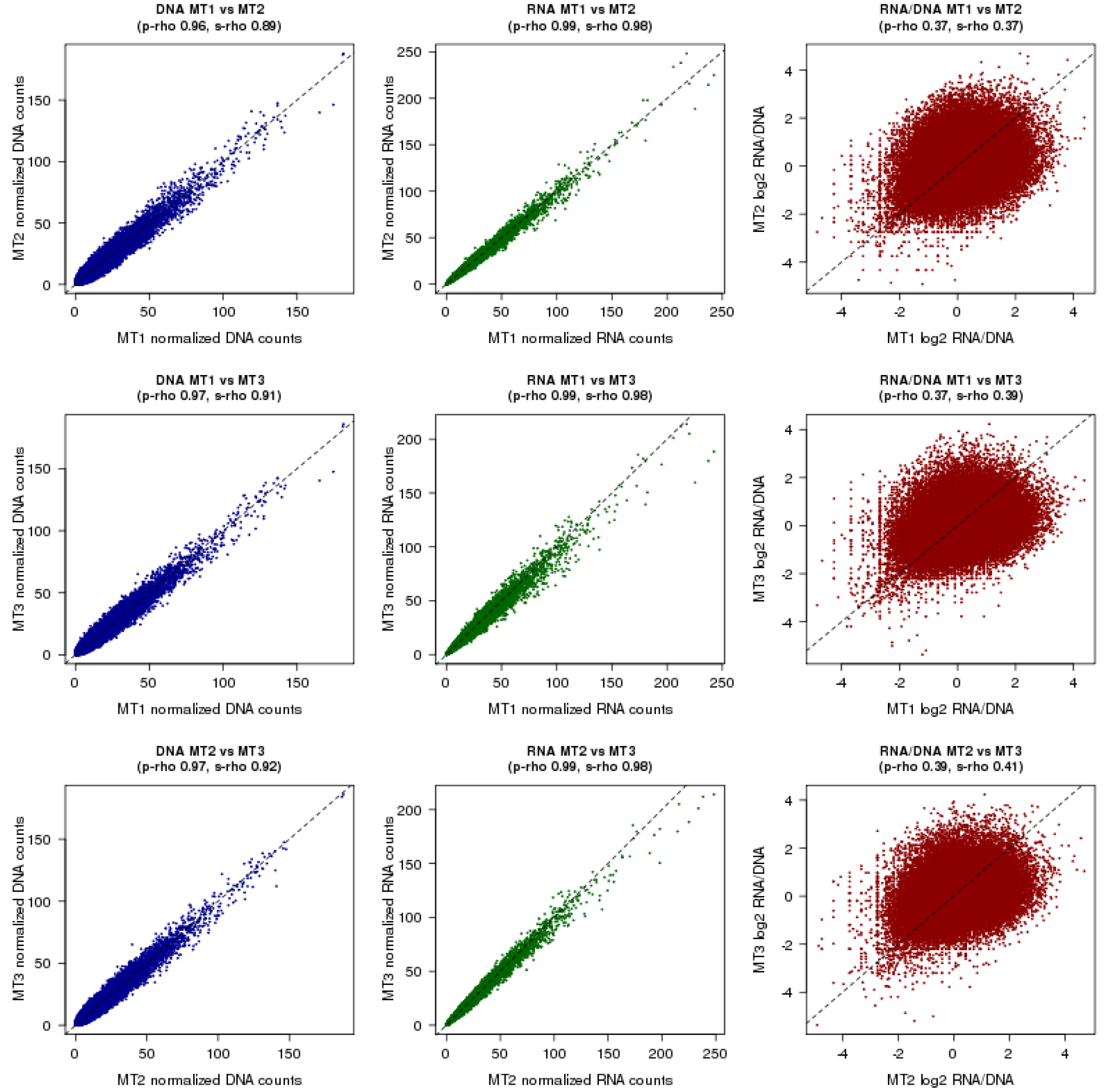
Correlation (Pearson/p-rho and Spearman/s-rho) of DNA tag counts (left column), RNA tag counts (middle column) as well as RNA/DNA ratios (right column) for pairwise comparisons of three mutant-integrase (MT) library replicates (rows). These are for individual barcodes, *i.e*. before summing across barcodes representing the same candidate enhancer sequence.

**Supplemental Figure S7.**
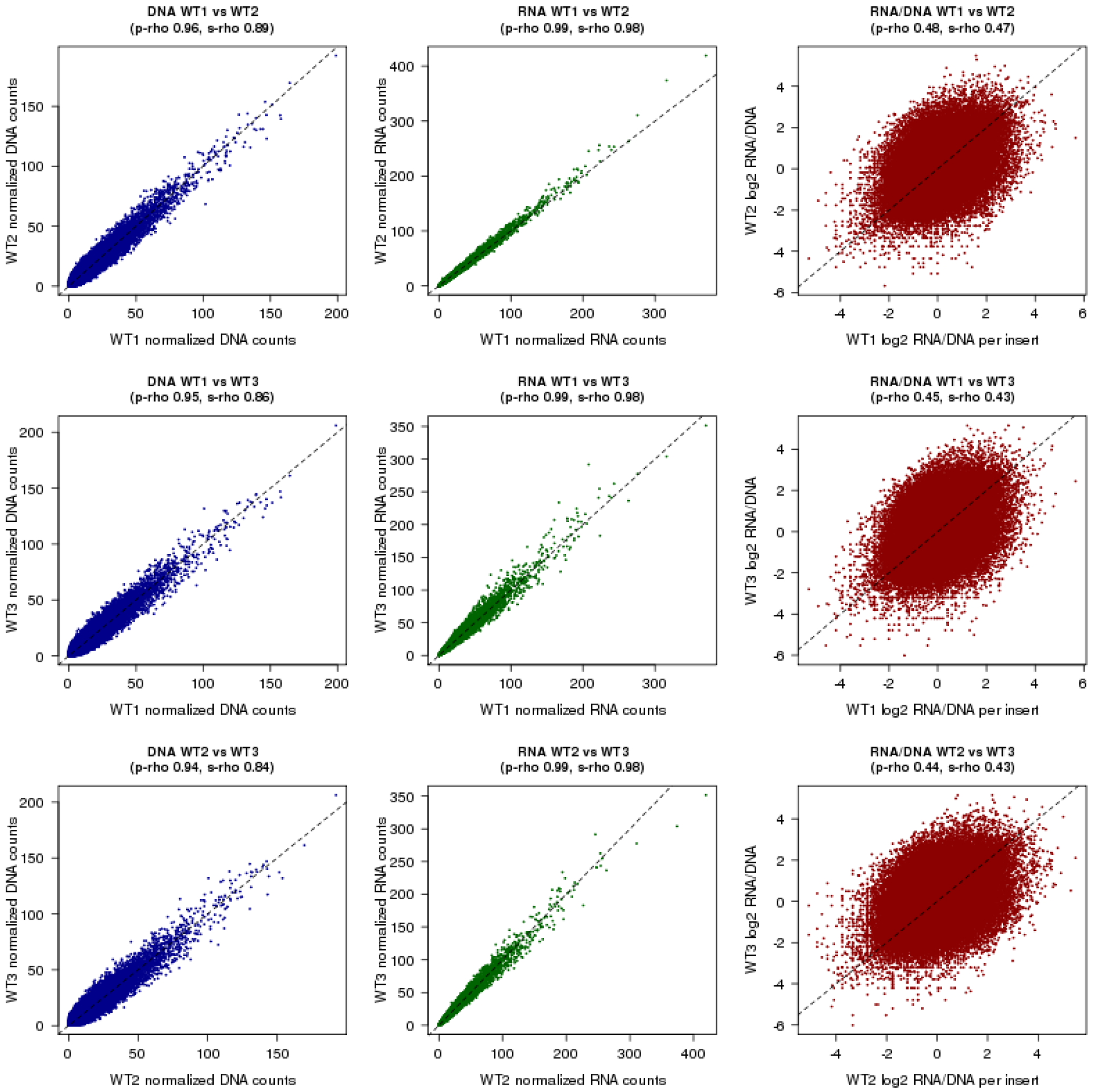
Correlation (Pearson/p-rho and Spearman/s-rho) of DNA tag counts (left column), RNA tag counts (middle column) as well as RNA/DNA ratios (right column) for pairwise comparisons of three wild-type-integrase (WT) library replicates (rows). These are for individual barcodes, *i.e*. before summing across barcodes representing the same candidate enhancer sequence.

**Supplemental Figure S8.**
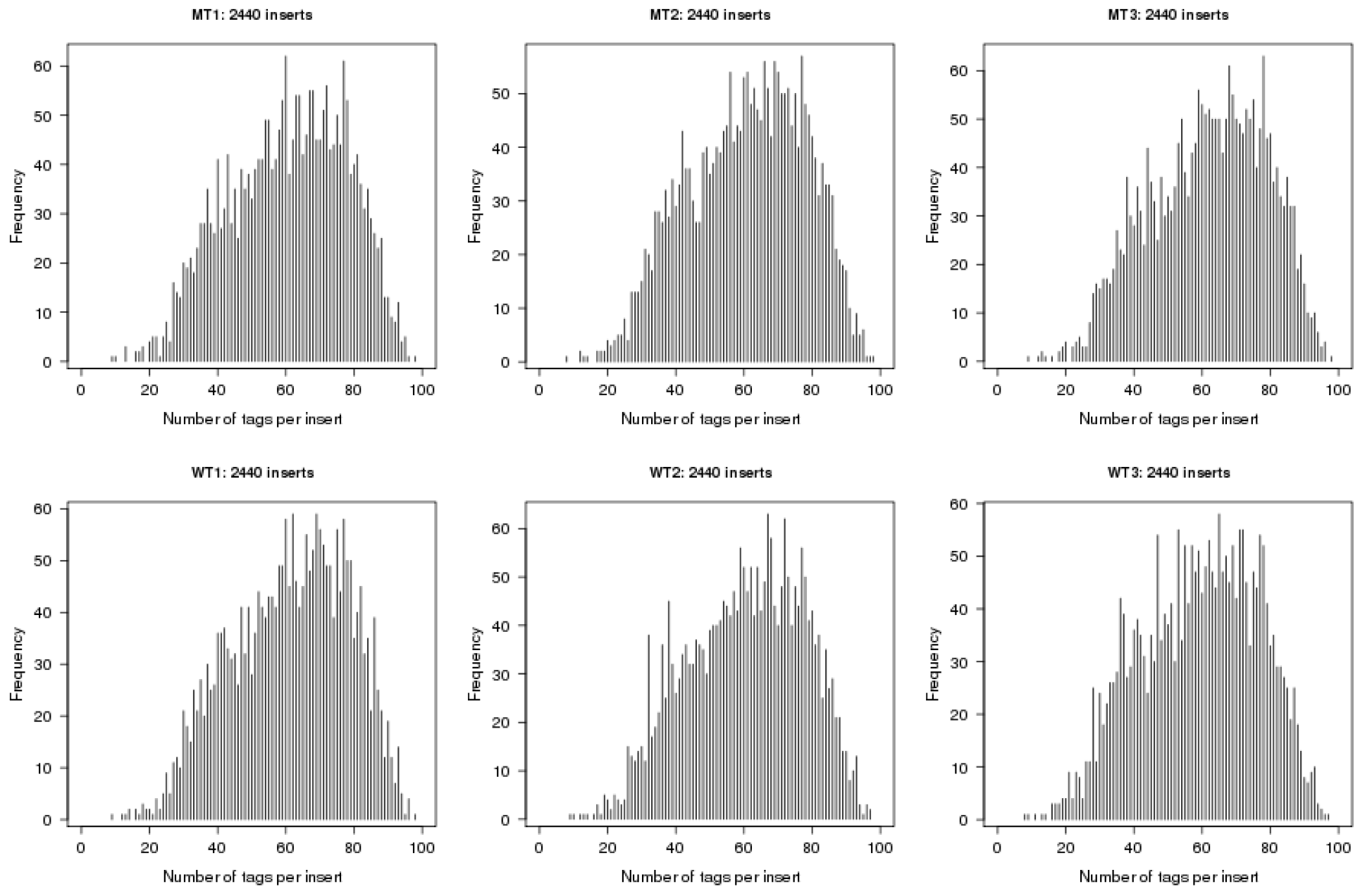
Histogram of the number of barcodes per insert across all 2,440 designed candidate enhancer sequences. We observe barcodes for all 2,440 inserts in each of the six experiments. On average, we observe 59-62 barcodes per insert (minimum 8-9, maximum 97-98).

**Supplemental Figure S9.**
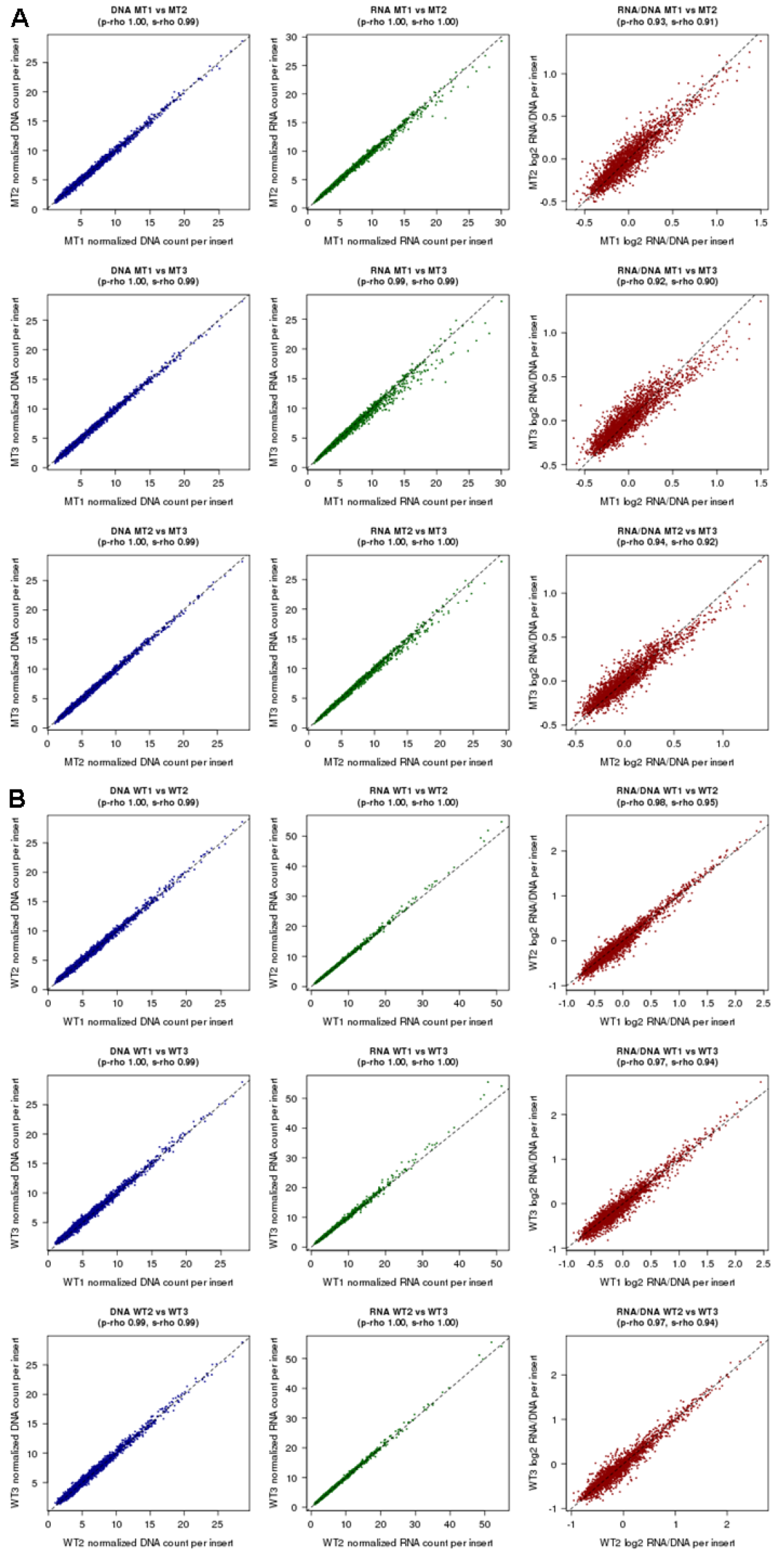
Reproducibility of counts and ratios across technical replicates. Correlation of DNA tag counts (left/blue), RNA tag counts (middle/green) as well as RNA/DNA ratios (right/red) for pairwise comparisons of the three mutant-integrase (MT) library replicates (*A*) and the three wild-type-integrase (WT) library replicates (*B*) along the rows. Both the counts and the ratios are calculated from RNA and DNA counts that are summed across ~60 barcodes representing a given candidate enhancer sequence and represented in a given experiment.

**Supplemental Figure S10.**
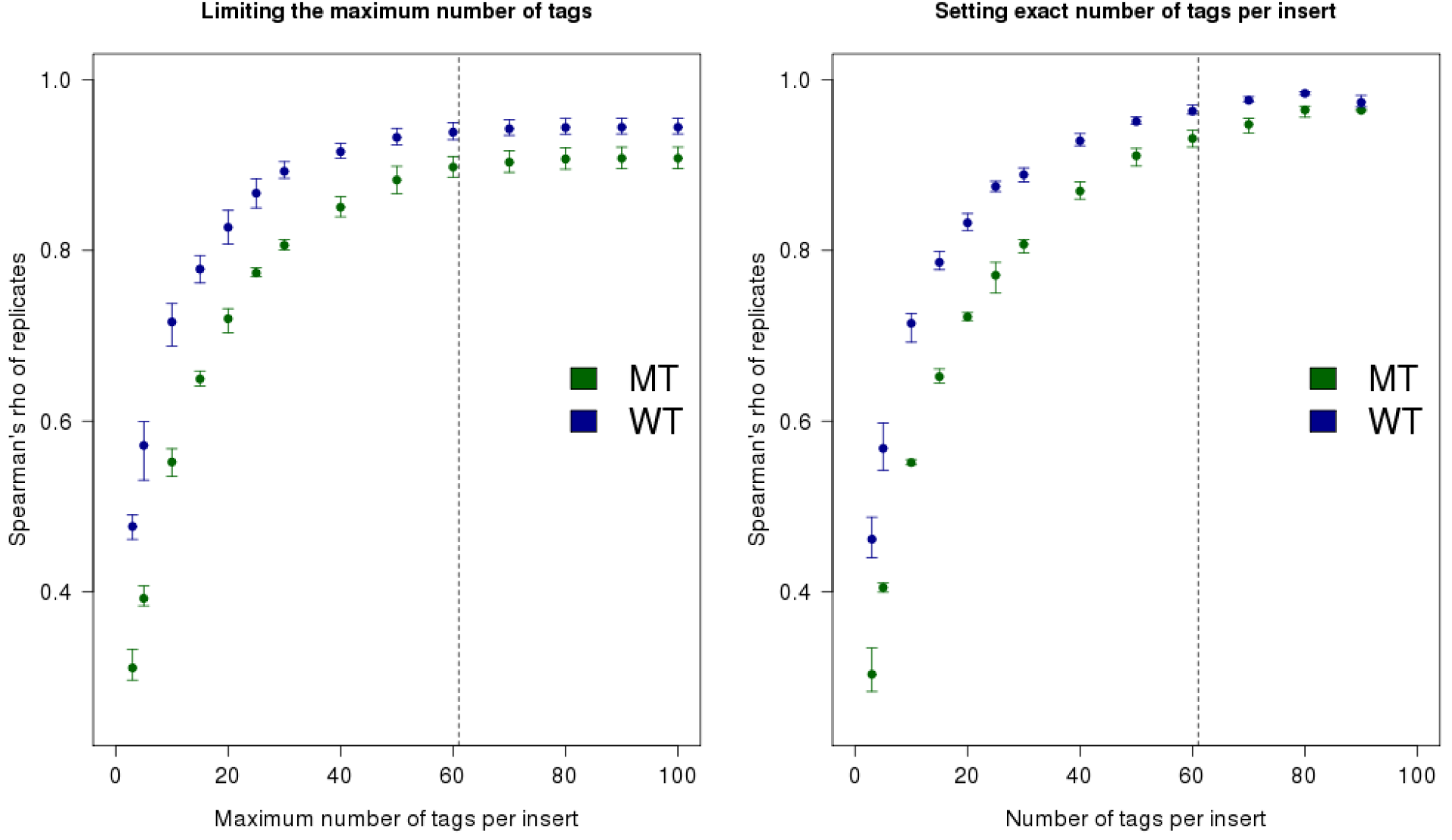
Effect of the number of barcodes averaged per insert on the correlation of experimental replicates for mutant (MT) and wild-type integrase (WT) experiments. Intervals indicate the mean, minimum and maximum values observed for pairwise Spearman correlations of the three replicates. Dashed lines indicate the average number of barcodes per insert as observed across all six experiments. The left plot is with setting a maximum number of tags per insert. The right plot is with fixing an exact number of tags per insert.

**Supplemental Figure S11.**
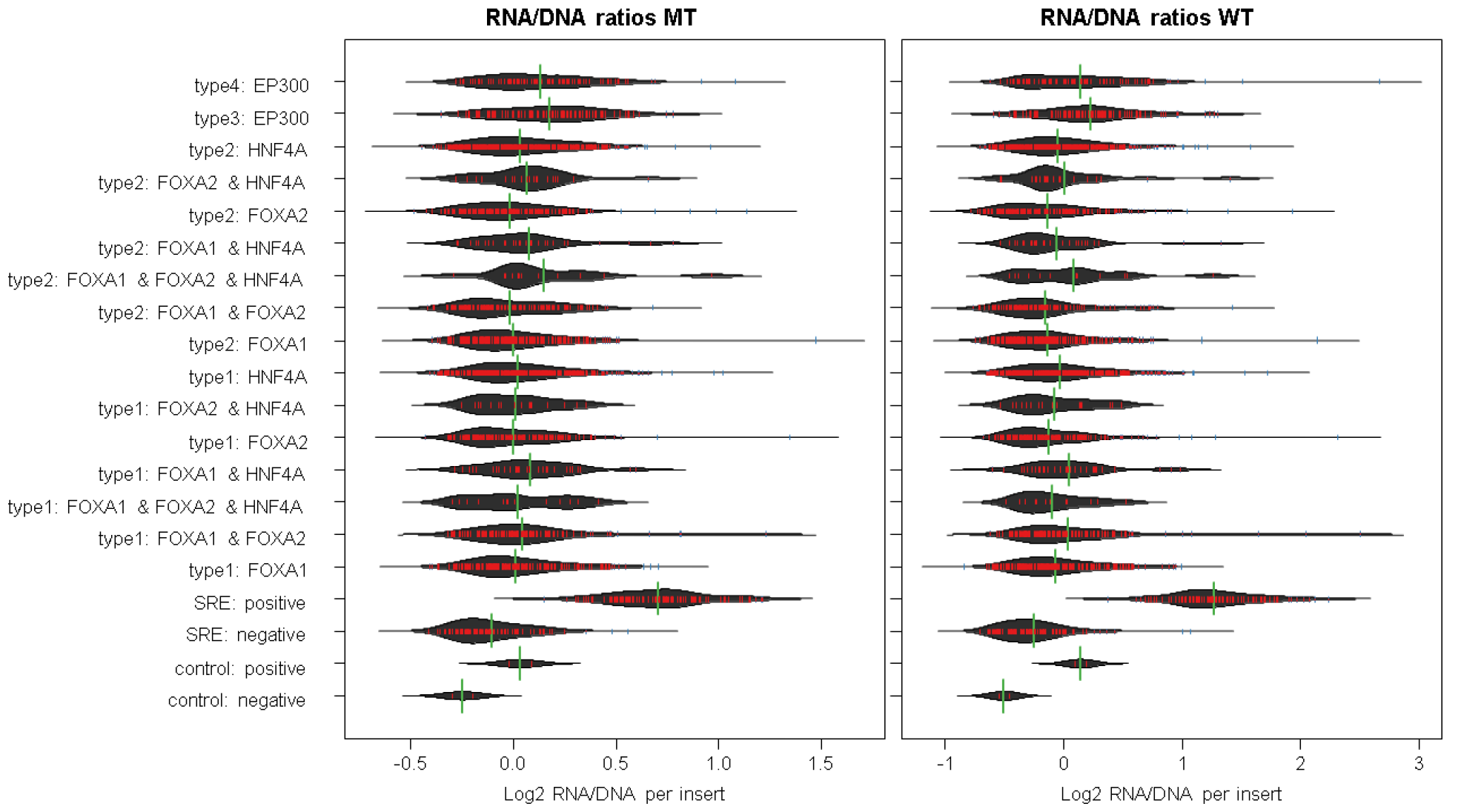
Bean plot of MT (left) and WT (right) RNA/DNA ratios split by design category. The four classes of putative enhancer elements are: Regions of FOXA1, FOXA2 or HNF4A binding that overlap H3K27ac and EP300 calls as well as at least one of three chromatin remodeling factors RAD21, CHD2 or SMC3 (type 1); Regions like in 1 but with no remodeling factor overlapping (type 2); EP300 peak regions overlapping H3K27ac as well as at least one of chromatin remodeling factor, but without peaks in FOXA1, FOXA2 or HNF4A (type 3); Regions like in 3 but with no remodeling factor overlapping (type 4). As shown here and in Fig. 3A, we do not observe major differences between design types, either with respect to activity or MT vs. WT.

**Supplemental Figure S12.**
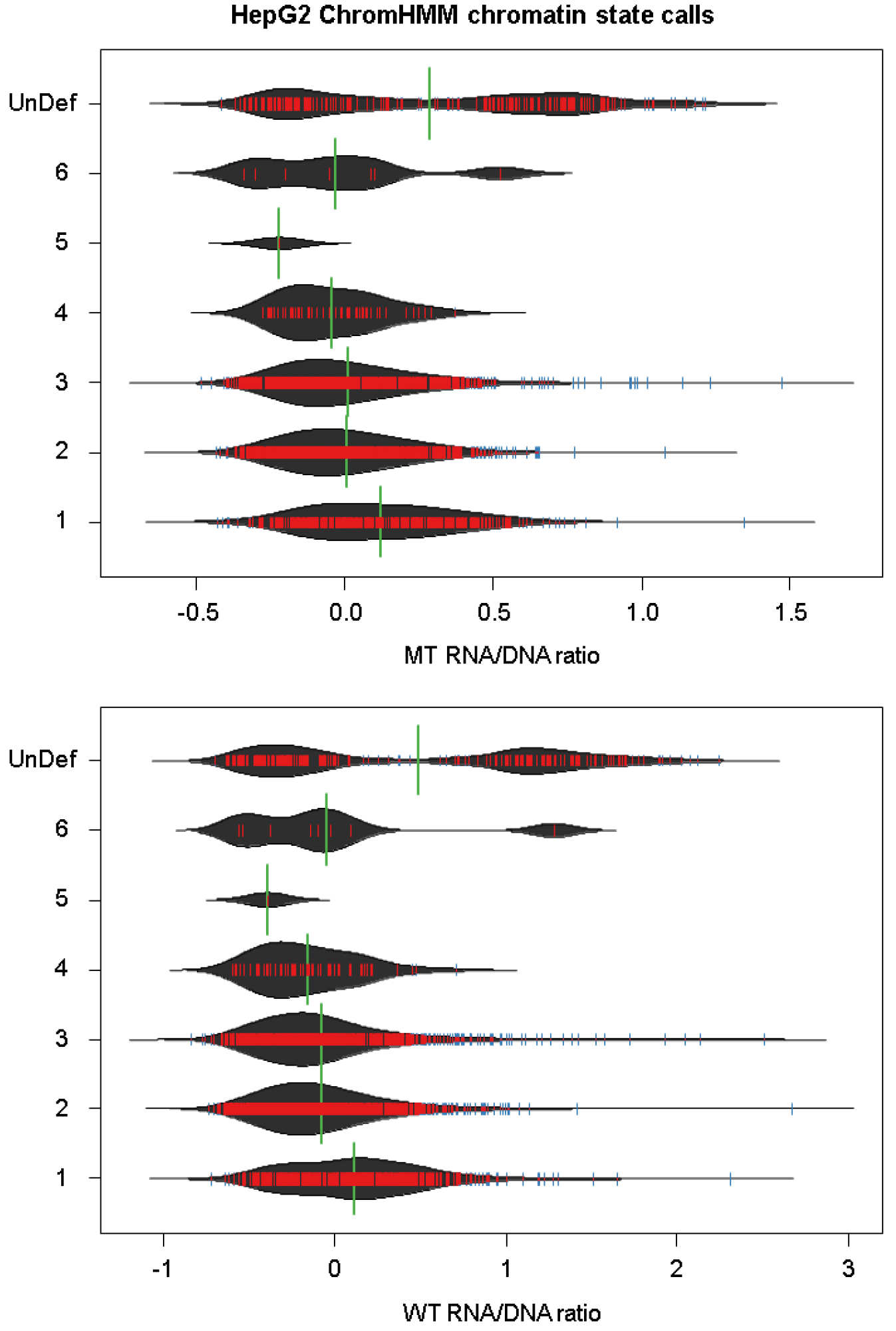
Distribution of RNA/DNA ratios for MT (top) and WT (bottom) split by HepG2 ChromHMM states. ChromHMM states were downloaded from the ENCODE project and had not been annotated with further labels. Inserts not represented in the available annotations were assigned to UnDef.

**Supplemental Figure S13.**
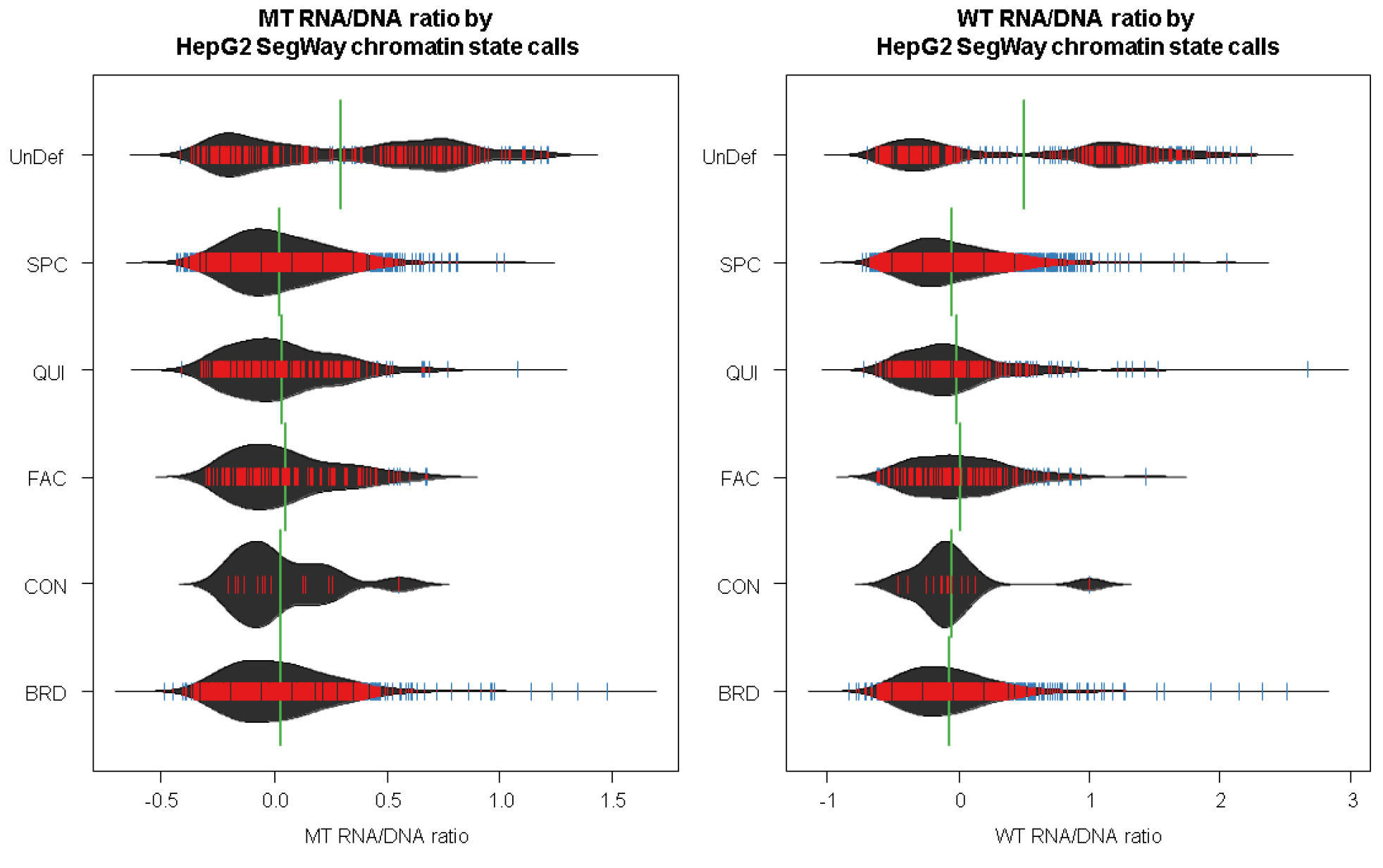
Distribution of RNA/DNA ratios for MT (left) and WT (right) split by HepG2 SegWay chromatin states. Chromatin labels were downloaded from the ENCODE project. Inserts not represented in the available annotations were assigned to UnDef.

**Supplemental Figure S14.**
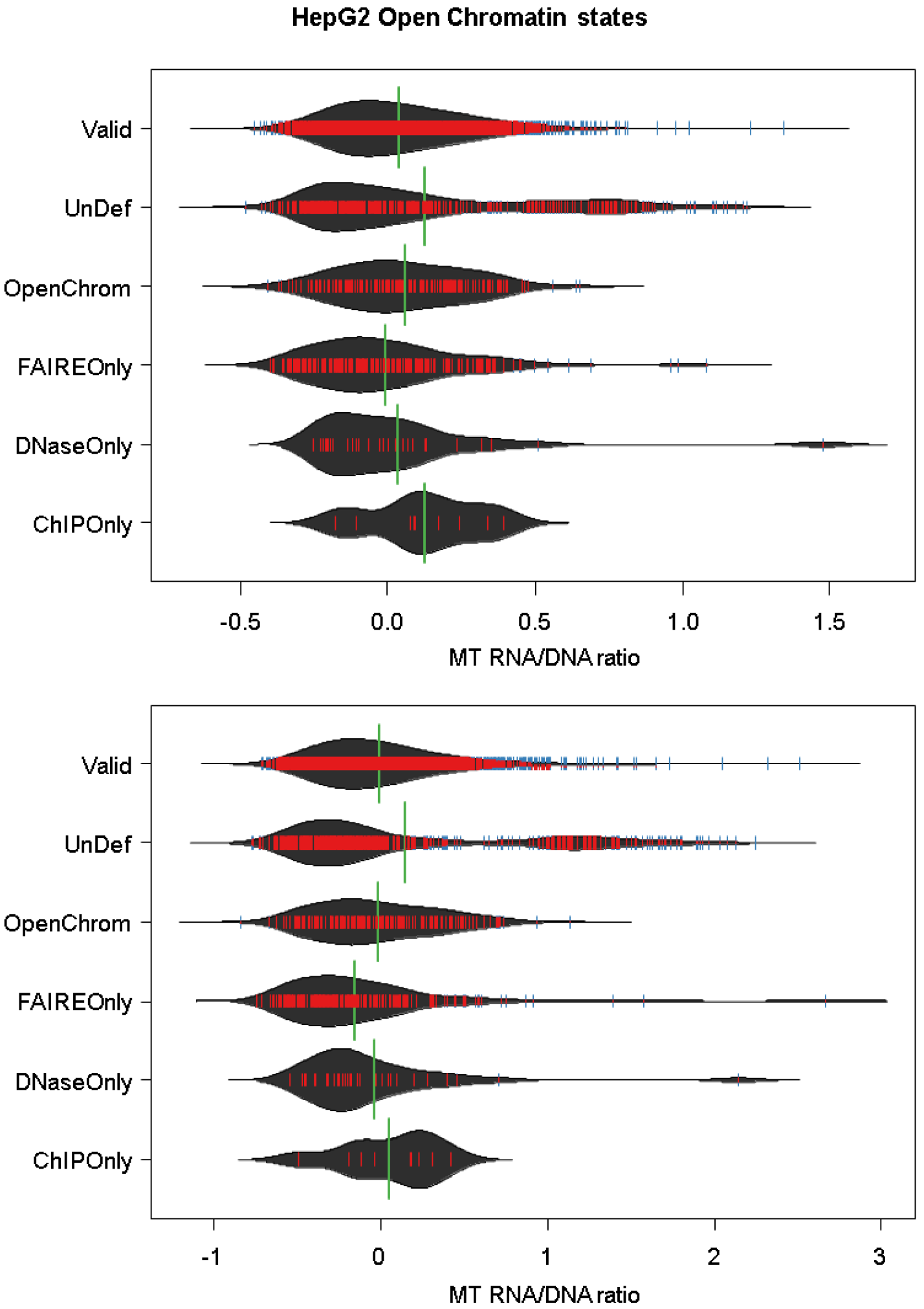
Distribution of RNA/DNA ratios for MT (top) and WT (bottom) split by HepG2 Open Chromatin states. Chromatin labels were downloaded from the ENCODE project. Inserts not represented in the available annotations were assigned to UnDef.

**Supplemental Figure S15.**
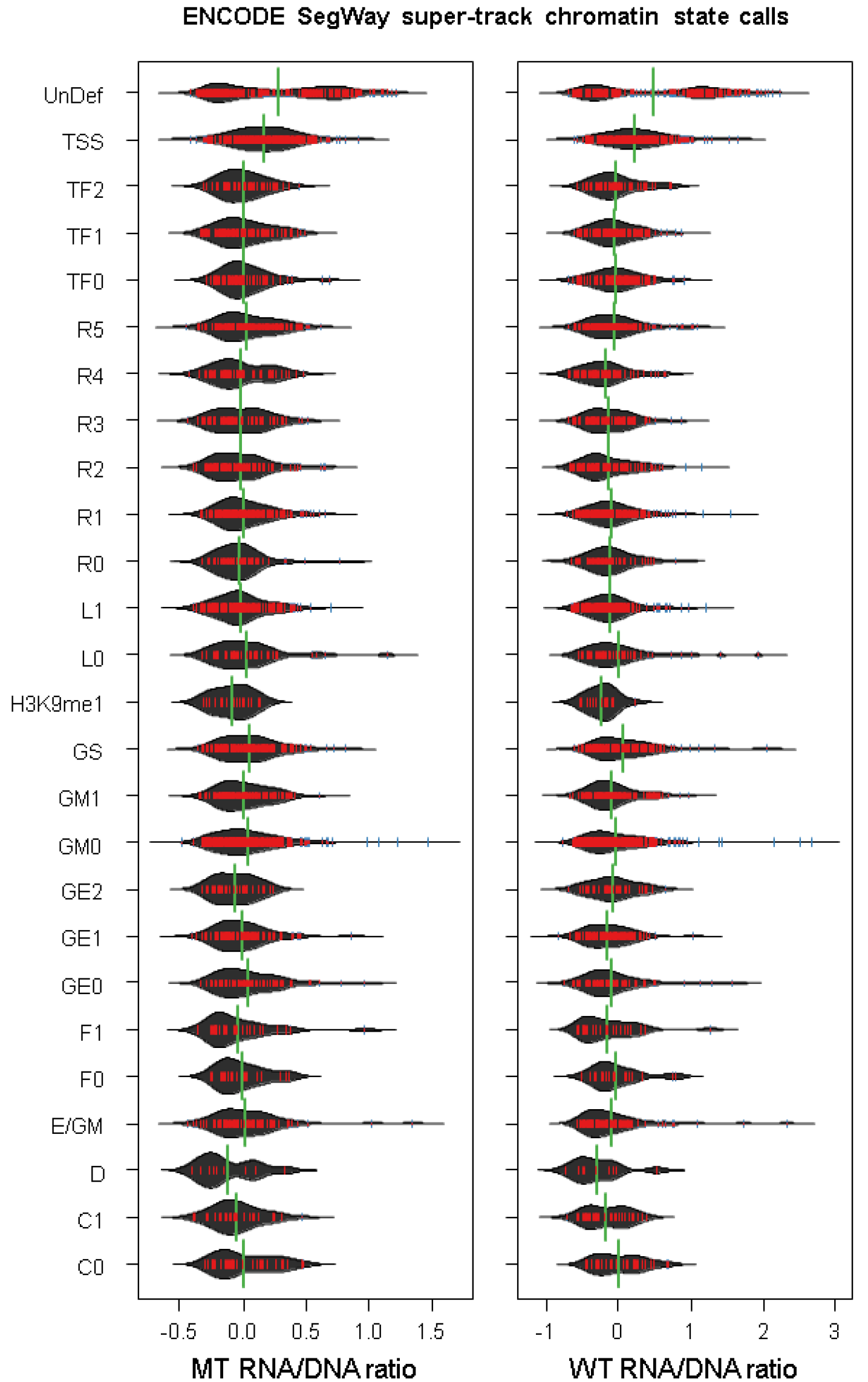
Distribution of RNA/DNA ratios for the MT (left) and WT (right) experiments split by SegWay chromatin states of the UCSC supertrack. Inserts not represented in the available annotations were assigned to UnDef.

**Supplemental Figure S16.**
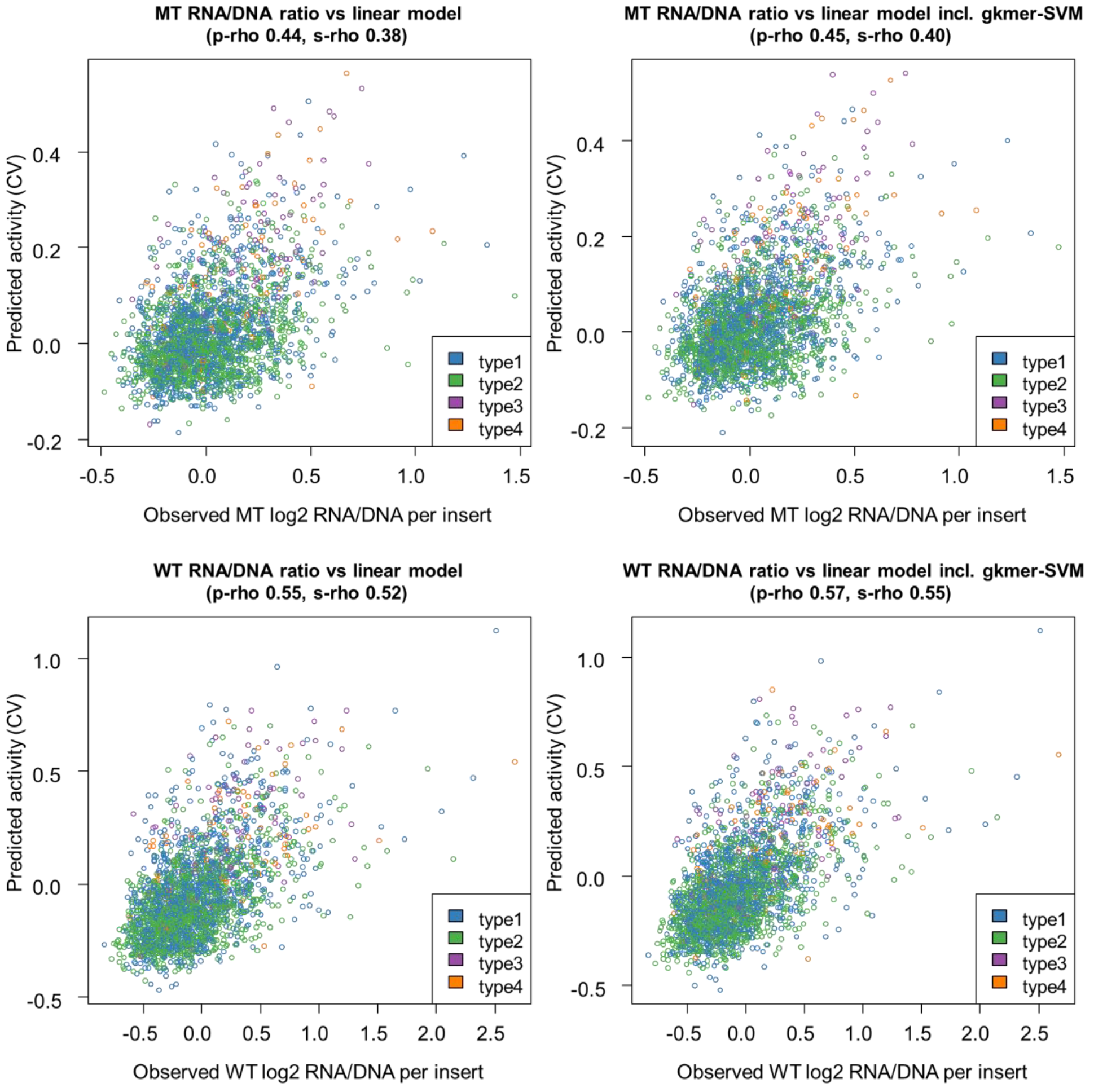
Correlation (Pearson/p-rho and Spearman/s-rho) of MT and WT RNA/DNA ratios fit in linear models derived from all genomic annotations with and without HepG2 gkm-SVM scores. SRES and other control sequences were excluded from this analysis as genomic annotations are mostly missing for those. WT linear models correlate better with the WT ratios than do MT linear models for the MT ratios (e.g. Spearman R^2^ of 0.2717 WT vs. 0.1459 for MT / Pearson R^2^ of 0.3069 for WT vs. 0.1931 for MT). Gapped-kmer SVM scores further improve R^2^ values especially for WT ratios (Spearman R^2^ 0.2975 for WT vs. 0.1581 for MT / Pearson R^2^ 0.3295 for WT vs. 0.2062 for MT).

**Supplemental Figure S17.**
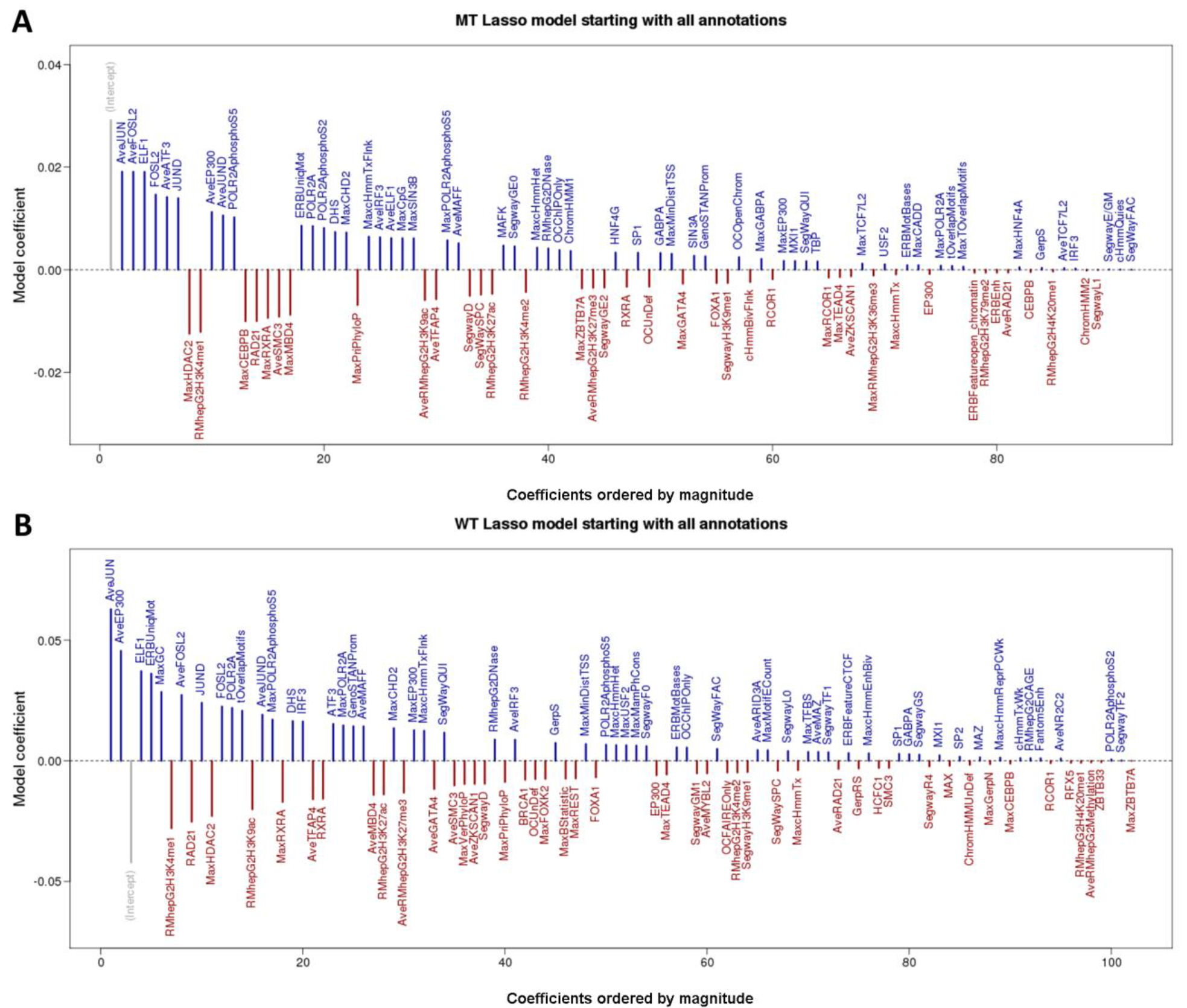
Coefficients for lasso models to predict RNA/DNA for MT (panel A) and WT (panel B) using genomic annotations, without including gkm-SVM score. The R glmnet package was used to fit the models, and the tuning parameter for each model was selected via 10-fold cross-validation. Categorical features were coded as K-1 binary columns, where K is the number of levels of the categorical feature. We excluded ZNF274 and EZH2 annotations from the model, because none of the inserts overlapped with these ChIP-seq tracks. All annotation features were scaled and centered before fitting the lasso model.

**Supplemental Figure S18.**
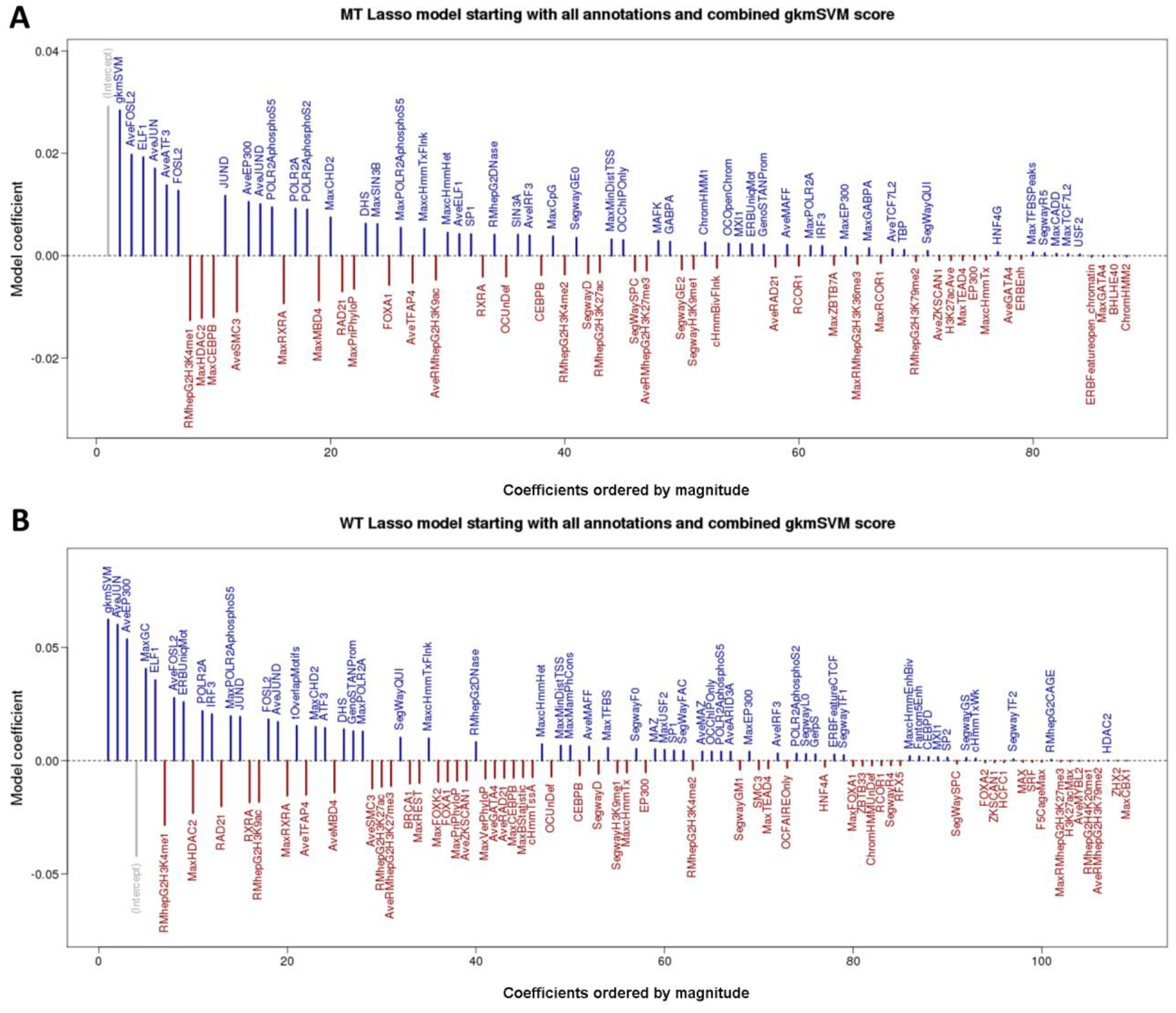
Coefficients for lasso models to predict RNA/DNA for MT (panel A) and WT (panel B) using genomic annotations and the combined HepG2 gkm-SVM score as predictors. Additional details are as in Figure S17. In the resulting models, the sequence based gkm-SVM score is assigned the highest model coefficient.

**Supplemental Figure S19.**
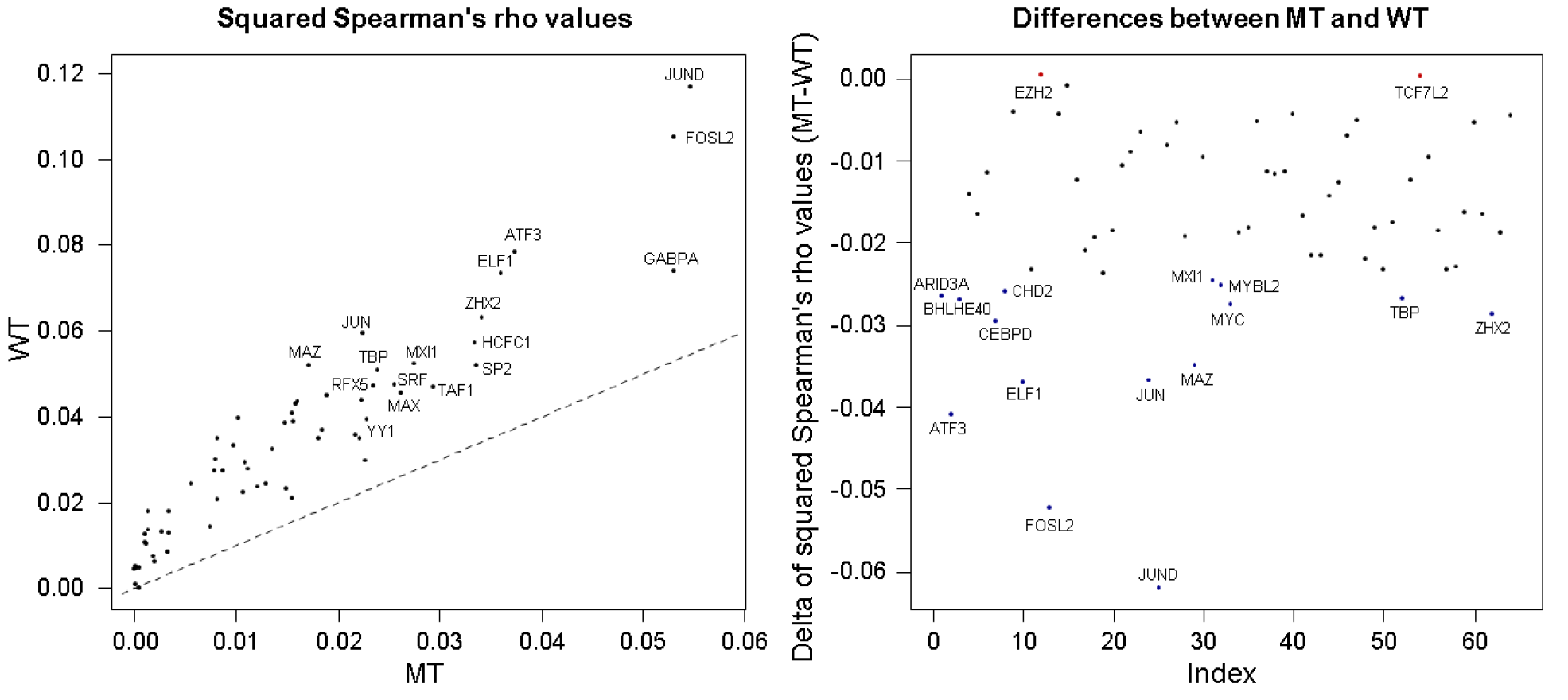
Spearman correlation coefficients (R^2^) of measured non-control insert activity with 64 LS-GKM models trained from HepG2 ChIP-Seq data. WT RNA/DNA ratios correlate better with annotations than the respective MT values. The left panel highlights the top correlated annotations for WT and MT ratios. The right panel highlights annotations with the largest difference in R^2^ values between MT and WT.

**Supplemental Figure S20.**
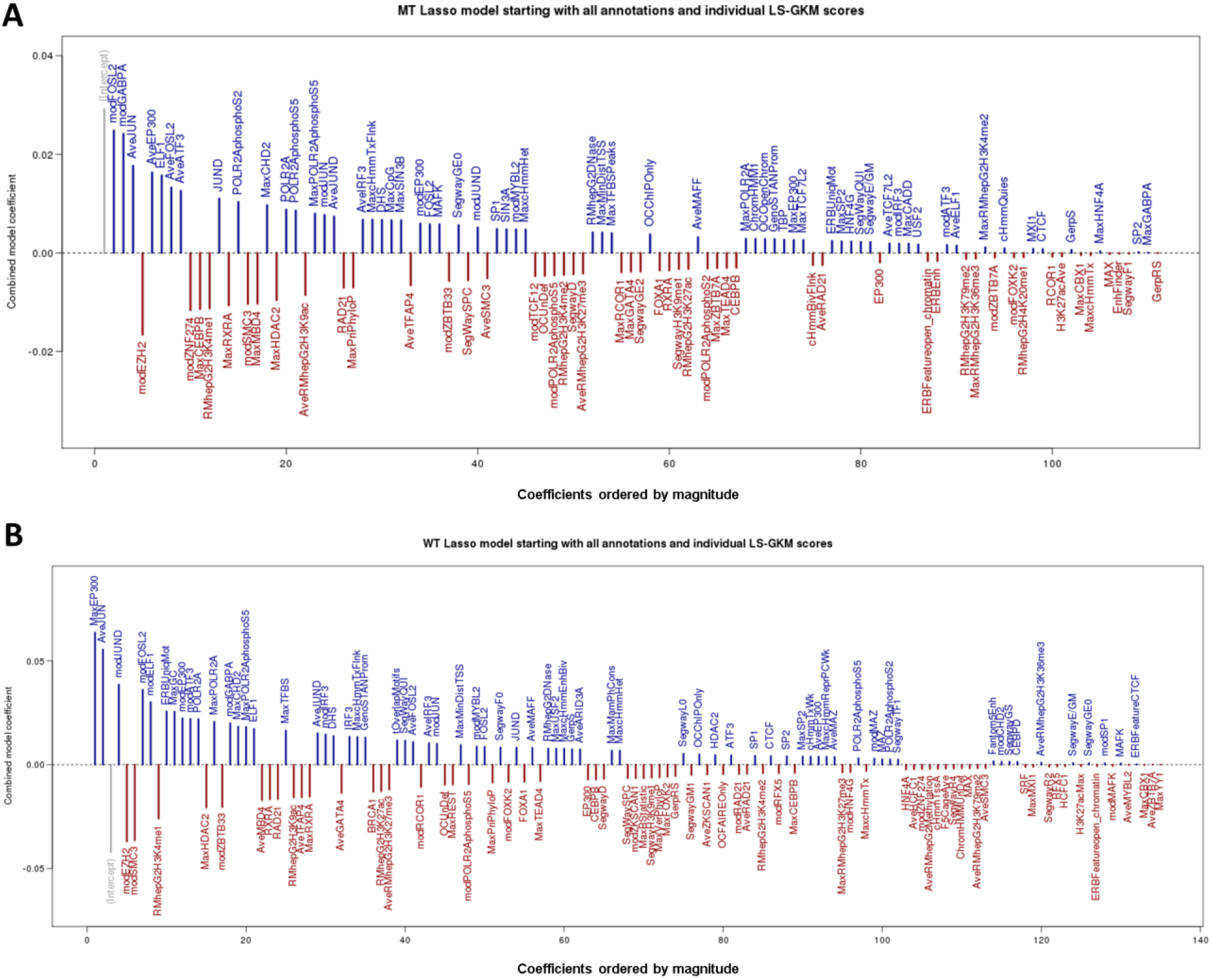
Coefficients for a lasso model that uses genomic annotations as well as LS-GKM scores to predict RNA/DNA ratios. Models were fit as described in the methods and summarized in Fig. S17. In addition, the 64 HepG2 ChIP-seq derived LS-GKM model scores were included as predictors (Mod-prefix).

**Supplemental Figure S21.**
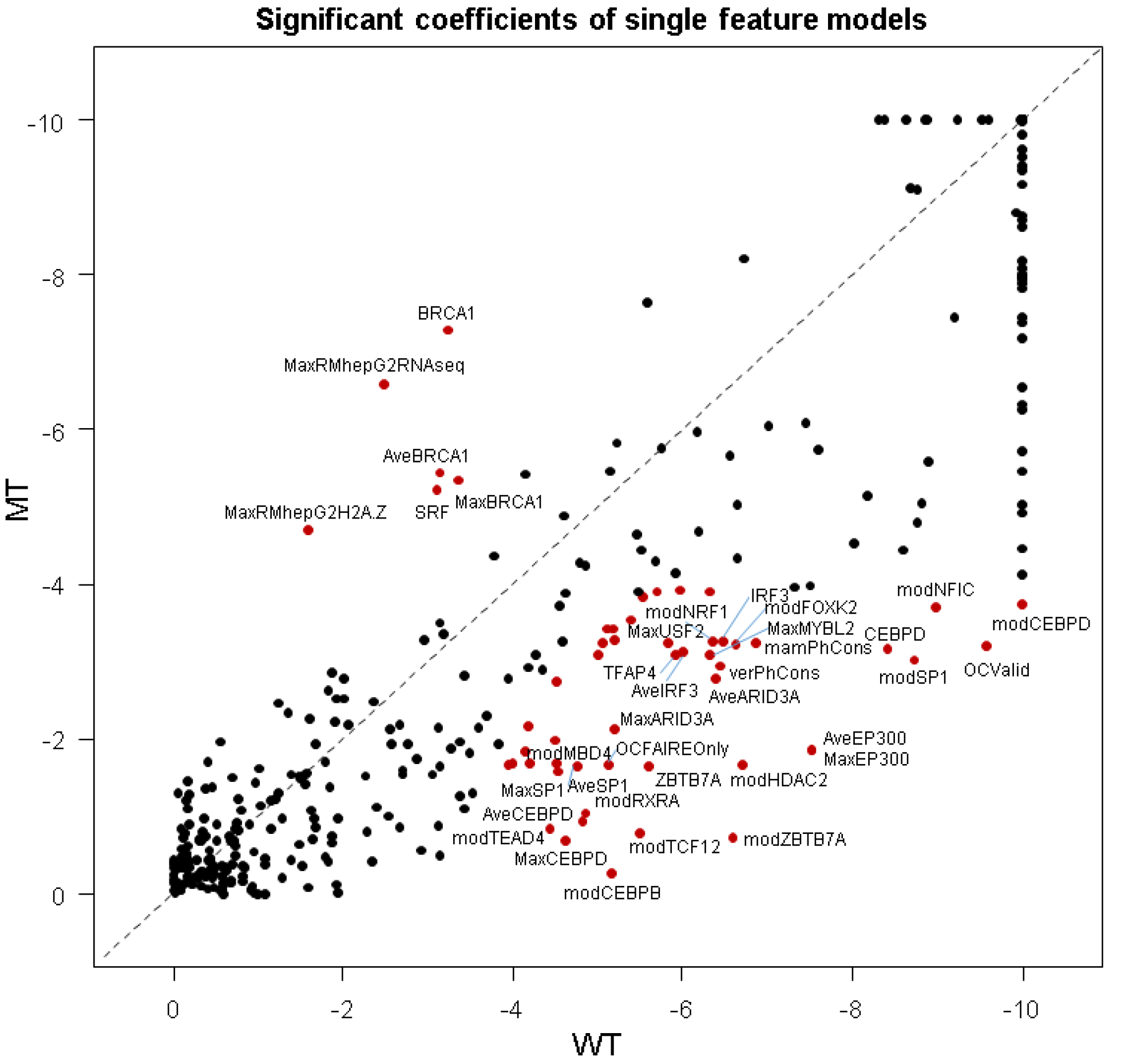
The results of 440 single-feature linear models predicting log2 RNA/DNA ratios using a single genomic annotation. The two-tailed p-value corresponding to the t-ratio based on a Student-t distribution (Table S7) is plotted for the inclusion of each coefficient in a single coefficient plus intercept linear model for predicting log2 RNA/DNA ratios for MT and WT experiments. For plotting purposes, p-values smaller than 10^−10^ were set to this threshold and p-values were log10 transformed. Highlighted in red are coefficients passing a p-value threshold of 0.05 (0.00012 after Bonferroni correction) in MT or WT while failing it in the other experiments. We also require a minimum p-value difference (0.0025) between MT and WT to account for stochasticity in the p-value estimates.

### Supplementary tables

**Supplemental Table S1.**
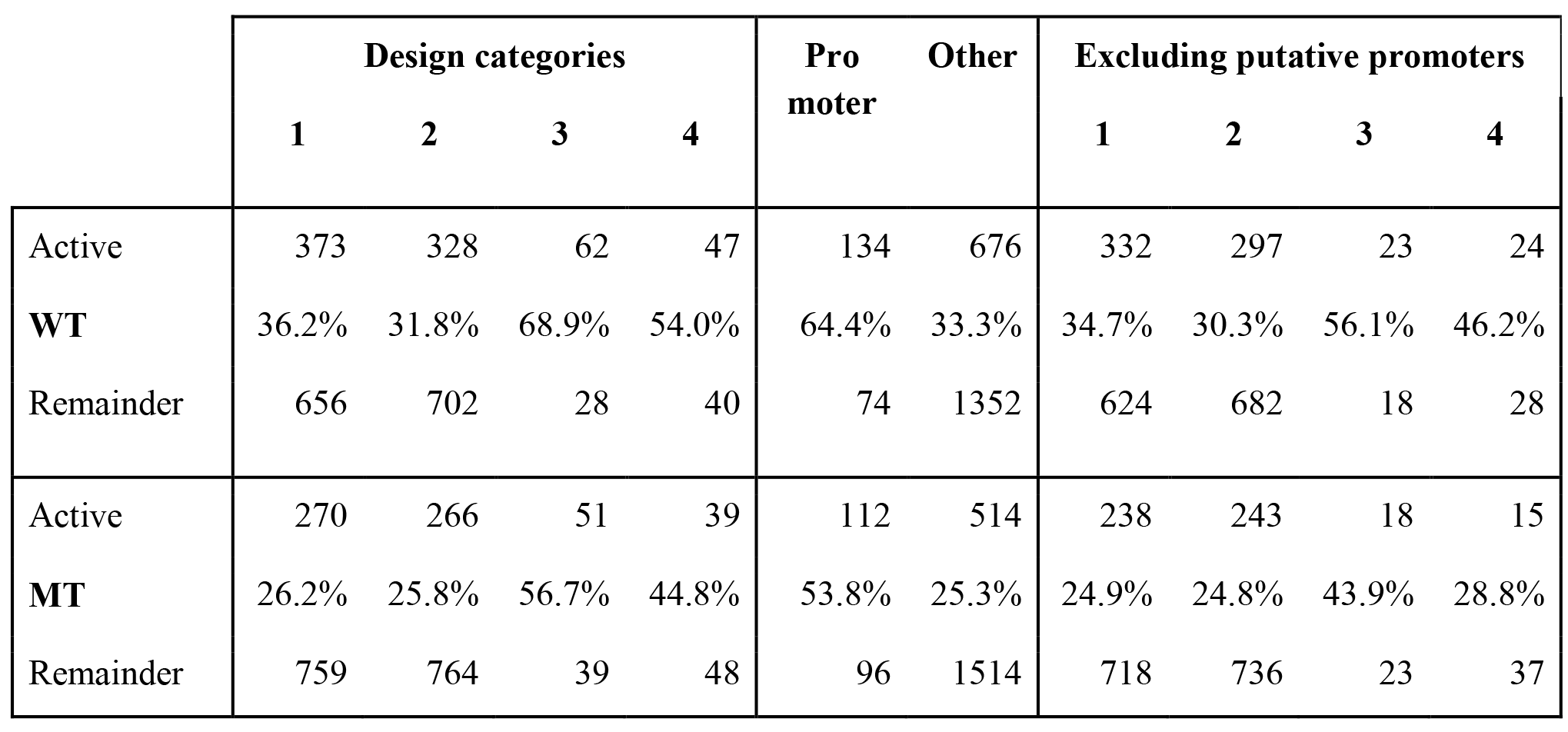
Active sequences by design category. The four classes of putative enhancer elements are: Regions of FOXA1, FOXA2 or HNF4A binding that overlap H3K27ac and EP300 calls as well as at least one of three chromatin remodeling factors RAD21, CHD2 or SMC3 (type 1); Regions like in 1 but with no remodeling factor overlapping (type 2); EP300 peak regions overlapping H3K27ac as well as at least one of chromatin remodeling factor, but without peaks in FOXA1, FOXA2 or HNF4A (type 3); Regions like in 3 but with no remodeling factor overlapping (type 4). Active sequences are defined as being above the 90^th^ percentile of RNA/DNA ratios observed for the negative synthetic controls (SRES). Promoters are defined based on a value less or equal to 1000 in the minDistTSS annotation field.

|  | Design categories |  |  |  | Pro<br>moter | Other | Excluding putative promoters |  |  |  |
| --- | --- | --- | --- | --- | --- | --- | --- | --- | --- | --- |
|  | 1 | 2 | 3 | 4 |  |  | 1 | 2 | 3 | 4 |
| Active | 373 | 328 | 62 | 47 | 134 | 676 | 332 | 297 | 23 | 24 |
| **WT** | 36.2% | 31.8% | 68.9% | 54.0% | 64.4% | 33.3% | 34.7% | 30.3% | 56.1% | 46.2% |
| Remainder | 656 | 702 | 28 | 40 | 74 | 1352 | 624 | 682 | 18 | 28 |
| Active | 270 | 266 | 51 | 39 | 112 | 514 | 238 | 243 | 18 | 15 |
| **MT** | 26.2% | 25.8% | 56.7% | 44.8% | 53.8% | 25.3% | 24.9% | 24.8% | 43.9% | 28.8% |
| Remainder | 759 | 764 | 39 | 48 | 96 | 1514 | 718 | 736 | 23 | 37 |

**Supplemental Table S2.** Performance of pooled sequence models measured as Spearman R^2^ with RNA/DNA ratios of MT and WT experiments. The column “gkmSVM” refers to the sequence-based model trained from combining individual ChIP-seq binding factors described by M Ghandi & D Lee et al. (Ghandi et al. 2014). JUND is the LS-GKM model trained only on JUND ChIP-seq peaks. This is the model showing the highest correlation with MT and WT RNA/DNA ratios. "Top 5", "Top 10" and "All Lasso" LS-GKM models were trained on the pooled ChIP-seq data underlying the individual TF LS-GKM models selected by linear Lasso regression. The "Top 5" factors are FOSL2, JUND, GABPA, EZH2, SMC3 (MT) as well as JUND, FOSL2, EZH2, JUN, and SMC3 (WT). The "Top 10" additionally includes MYBL2, FOXA2, CEBPB, JUN, ATF3 (MT) as well as ELF1, MYBL2, ATF3 GABPA, ZBTB33 (WT). The full set of factors used for "All Lasso" (in alphabetical order) is ATF3, BRCA1, CEBPB, ELF1, EZH2, FOSL2, FOXA1, FOXA2, FOXK2, GABPA, HCFC1, HNF4G, IRF3, JUN, JUND, MAFF, MYBL2, NR2C2, POLR2A, POLR2AphosphoS2, POLR2AphosphoS5, RAD21, RCOR1, RFX5, SIN3B, SMC3, SP2, SRF, TCF12, TEAD4, TFAP4, ZBTB33, ZBTB7A, ZNF274 (MT) and ARID3A, ATF3, CEBPB, CEBPD, CHD2, ELF1, EZH2, FOSL2, FOXA1, FOXA2, FOXK2, GABPA, HNF4G, IRF3, JUN, JUND, MAFK, MAZ, MYBL2, MYC, NR2C2, POLR2A, POLR2AphosphoS2, POLR2AphosphoS5, RAD21, RCOR1, RFX5, SIN3A, SMC3, SP2, SRF, TAF1, TBP, TEAD4, TFAP4, ZBTB33, ZBTB7A, ZKSCAN1 (WT).

| Experiment | gkmSVM | JUND | Top 5 | Top 10 | All Lasso |
| --- | --- | --- | --- | --- | --- |
| WT | 7.56% | 11.68% | 13.76% | 15.18% | 9.39% |
| MT | 4.09% | 5.47% | 8.16% | 5.04% | 5.05% |

**Supplemental Table S3.**
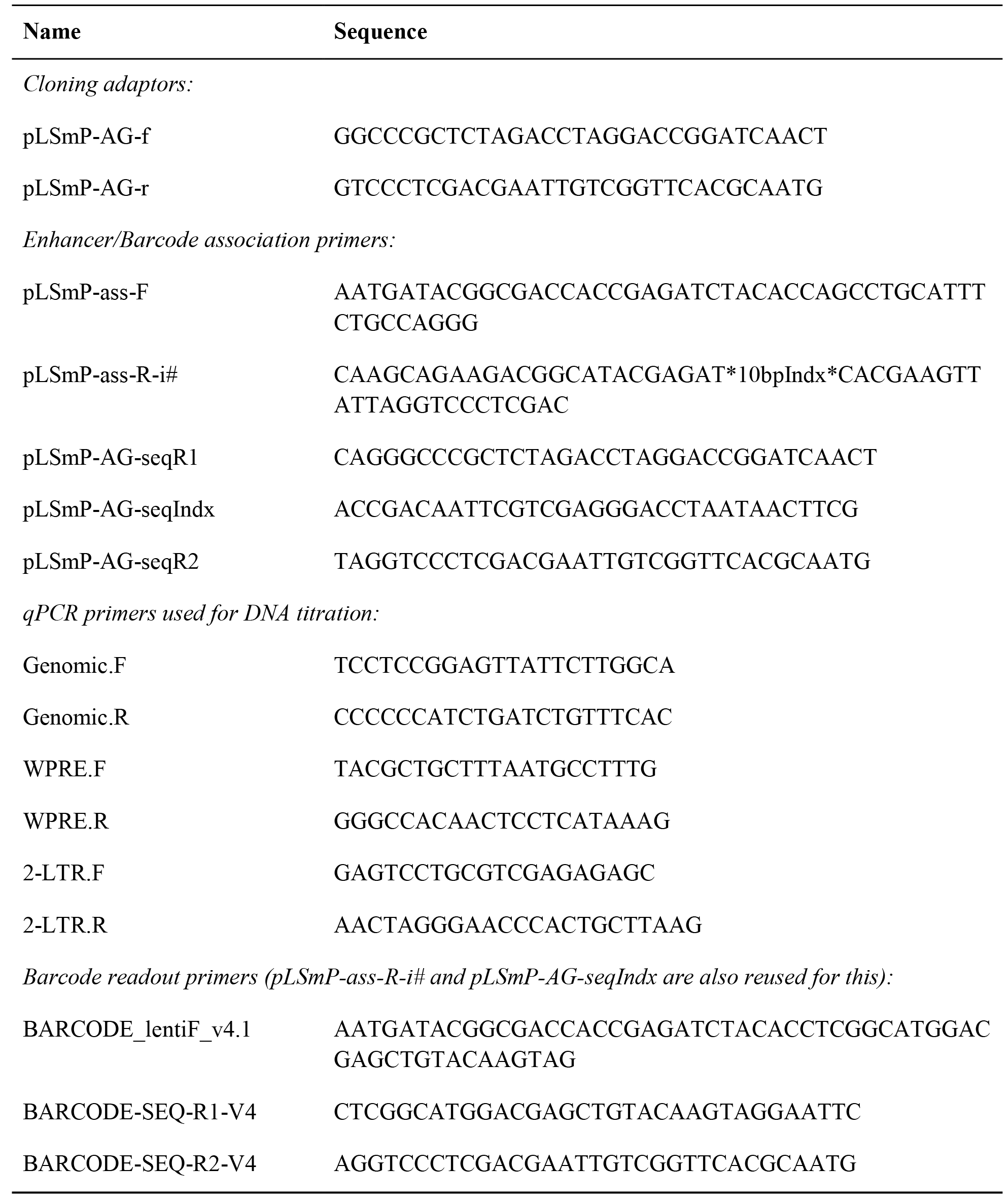
Primer sequences used for cloning, barcode amplification and sequencing.

**Supplemental Table S4.** Proportion of correct oligos for increasing number of observations / number of sequences going into consensus calling. The major effect is due to insertion/deletion (InDels) events in the oligos, substitutions have a minor effect and are removed in the consensus calling process as long as the correct sequence is the most abundant sequence.

| Min.<br>sequence<br>count | Oligos<br>without<br>InDels | Correct oligos | All oligos | Percent<br>InDels | Percent<br>correct |
| --- | --- | --- | --- | --- | --- |
| 0 | 197752 | 190093 | 992513 | 19.92% | 19.15% |
| 5 | 191537 | 184487 | 870677 | 22.00% | 21.19% |
| 10 | 171625 | 166177 | 547199 | 31.36% | 30.37% |
| 15 | 151109 | 147218 | 312297 | 48.39% | 47.14% |
| 20 | 132546 | 129953 | 188893 | 70.17% | 68.80% |
| 25 | 115851 | 114238 | 132995 | 87.11% | 85.90% |
| 30 | 101675 | 100728 | 106507 | 95.46% | 94.57% |
| 35 | 89282 | 88754 | 90699 | 98.44% | 97.86% |
| 40 | 78521 | 78224 | 78976 | 99.42% | 99.05% |
| 45 | 69247 | 69081 | 69434 | 99.73% | 99.49% |
| 50 | 61229 | 61144 | 61317 | 99.86% | 99.72% |
| 55 | 54256 | 54212 | 54300 | 99.92% | 99.84% |
| 60 | 48228 | 48205 | 48253 | 99.95% | 99.90% |
| 65 | 43094 | 43076 | 43110 | 99.96% | 99.92% |
| 70 | 38746 | 38730 | 38755 | 99.98% | 99.94% |
| 75 | 34866 | 34853 | 34873 | 99.98% | 99.94% |
| 80 | 31434 | 31426 | 31437 | 99.99% | 99.97% |
| 85 | 28429 | 28423 | 28431 | 99.99% | 99.97% |
| 90 | 25801 | 25795 | 25803 | 99.99% | 99.97% |
| 95 | 23502 | 23497 | 23503 | 100.00% | 99.97% |
| 100 | 21422 | 21418 | 21423 | 100.00% | 99.98% |

**Supplemental Table S5.**
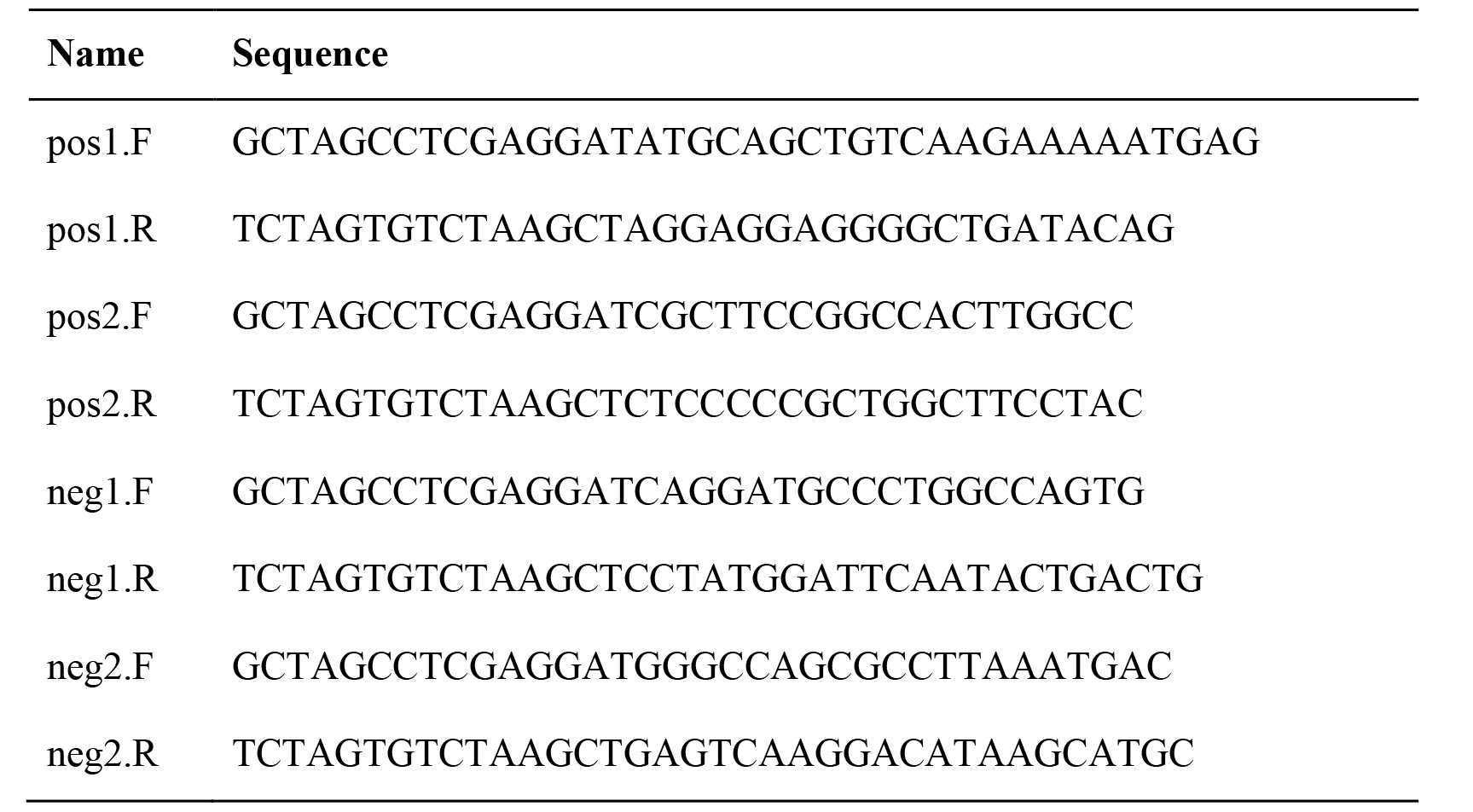
Primers used for cloning 2 negative and 2 positive control sequences in Luciferase Assay experiments.

**Supplemental Table S6.**
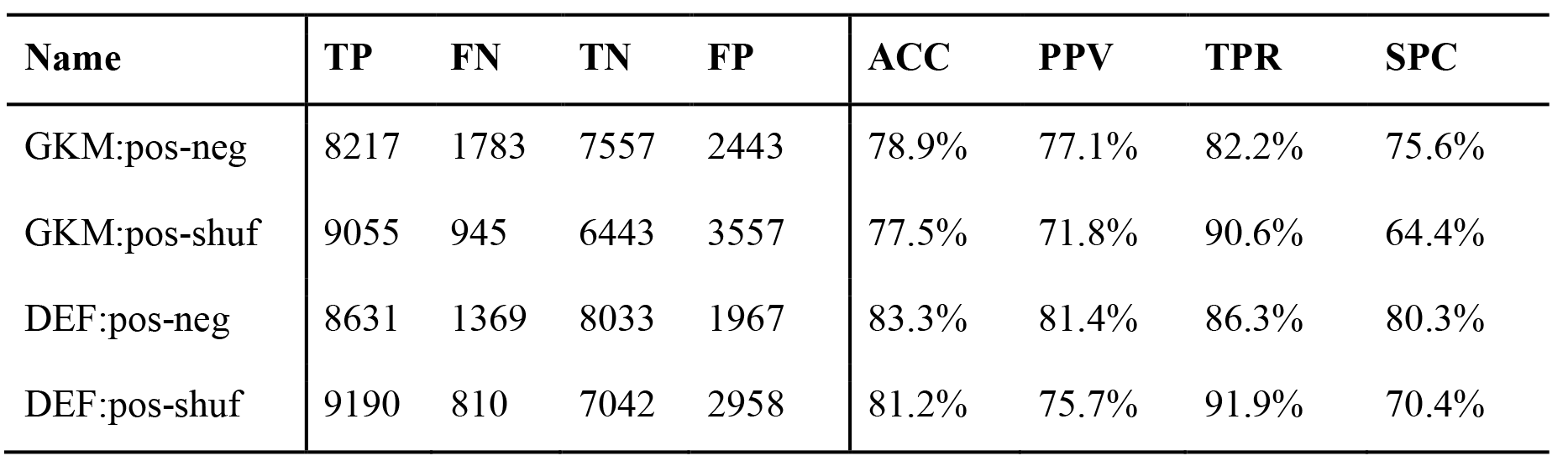
LS-GKM training on 225,327 HepG2 combined ChIP-seq peaks (positive set) and a positive validation set of 10,000 left-out sequences. Training was performed using LS-GKM defaults (DEF, -T4 -e0.01) and parameters matching gkm-SVM (GKM, -l 10 -k6 -d3 -t2 -T4 -e0.01). Negative sequences were either created by permuting the positive sequence while maintaining dimer-content or used the negative sequence set provided with gkm-SVM.

| Name | TP | FN | TN | FP | ACC | PPV | TPR | SPC |
| --- | --- | --- | --- | --- | --- | --- | --- | --- |
| GKM:pos-neg | 8217 | 1783 | 7557 | 2443 | 78.9% | 77.1% | 82.2% | 75.6% |
| GKM:pos-shuf | 9055 | 945 | 6443 | 3557 | 77.5% | 71.8% | 90.6% | 64.4% |
| DEF:pos-neg | 8631 | 1369 | 8033 | 1967 | 83.3% | 81.4% | 86.3% | 80.3% |
| DEF:pos-shuf | 9190 | 810 | 7042 | 2958 | 81.2% | 75.7% | 91.9% | 70.4% |

**Supplemental Table S7.**
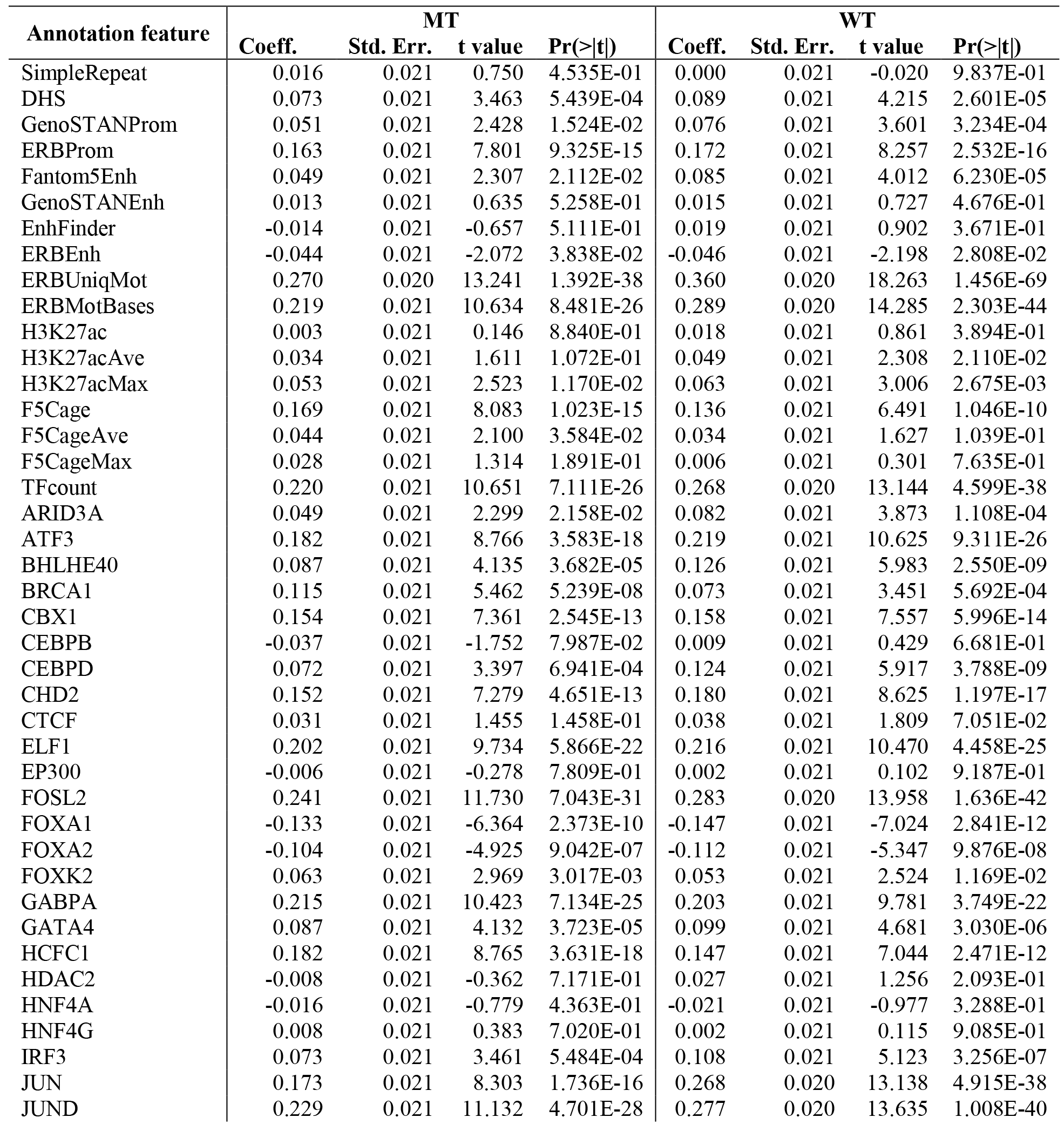
Model coefficients determined for 440 one feature models (single coefficient plus offset) for predicting log2 RNA/DNA ratios for MT and WT experiments. Categorical features were included as n-1 binary columns, where n is the number of levels of the categorical feature. ZNF274 and EZH2 annotations (but not the sequence based models modZNF274 and modEZH2) as none of the inserts overlapped with these ChIP-seq tracks. All annotation features and the output variable (MT/WT log2 RNA/DNA ratios) were scaled and centered before fitting the models to allow interpretation of coefficient values. The table provides the coefficient estimate (Coeff.), standard error (Std. Err.), t-values, and Pr(>|t|), which is the two-tailed p-value corresponding to the t-ratio based on a Student-t distribution. These values are returned by the R glm.summary routine for each model.

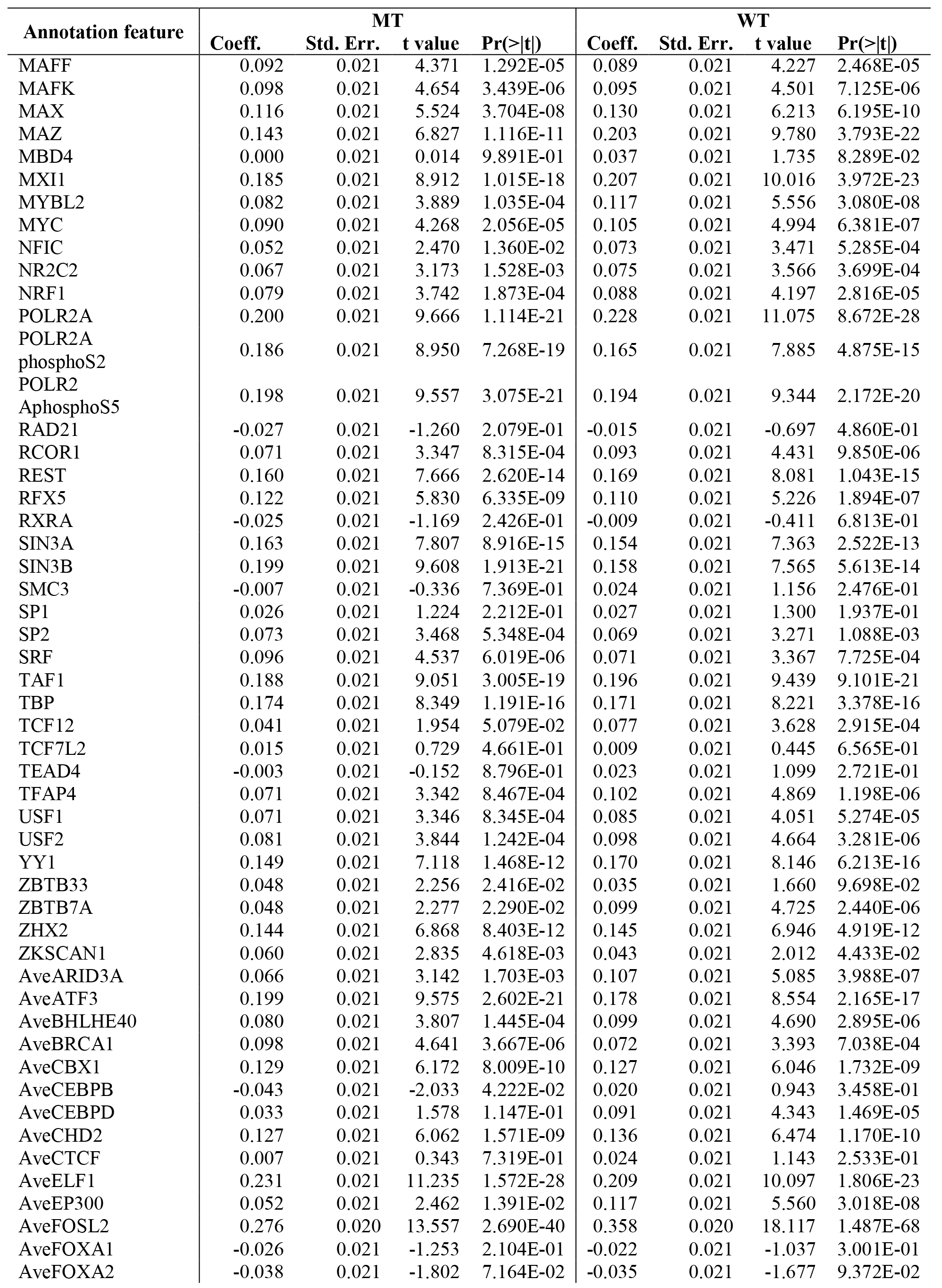

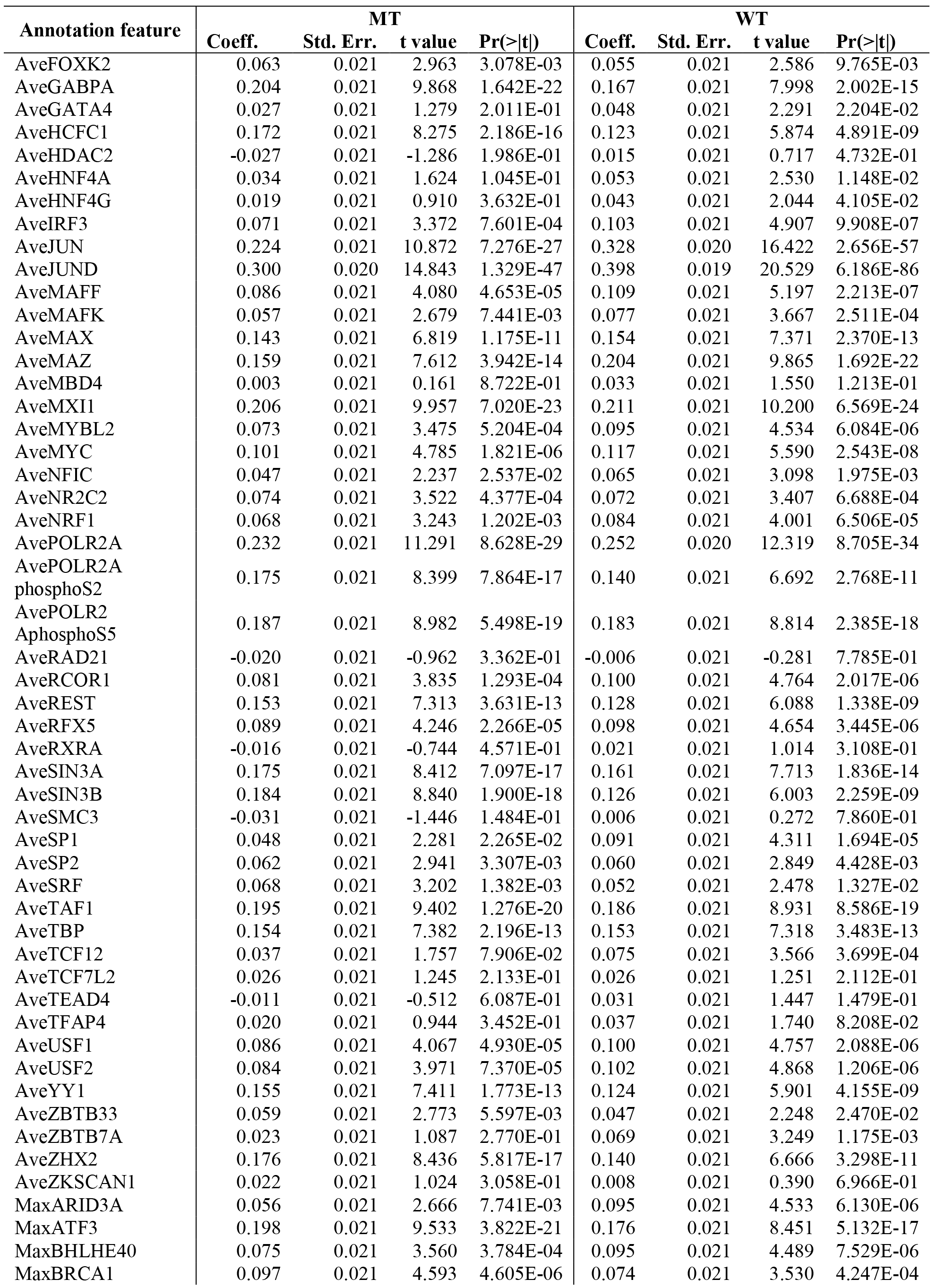

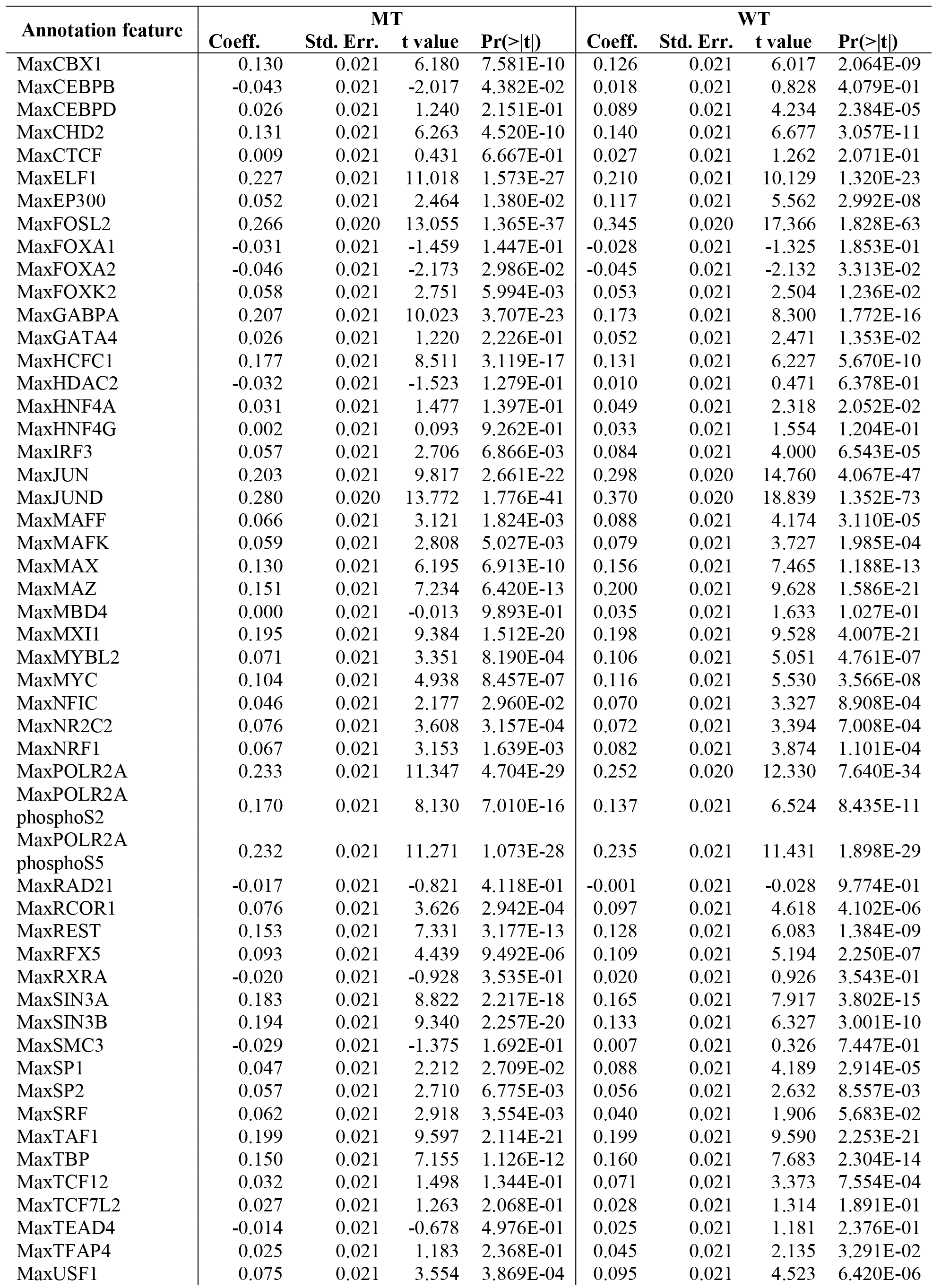

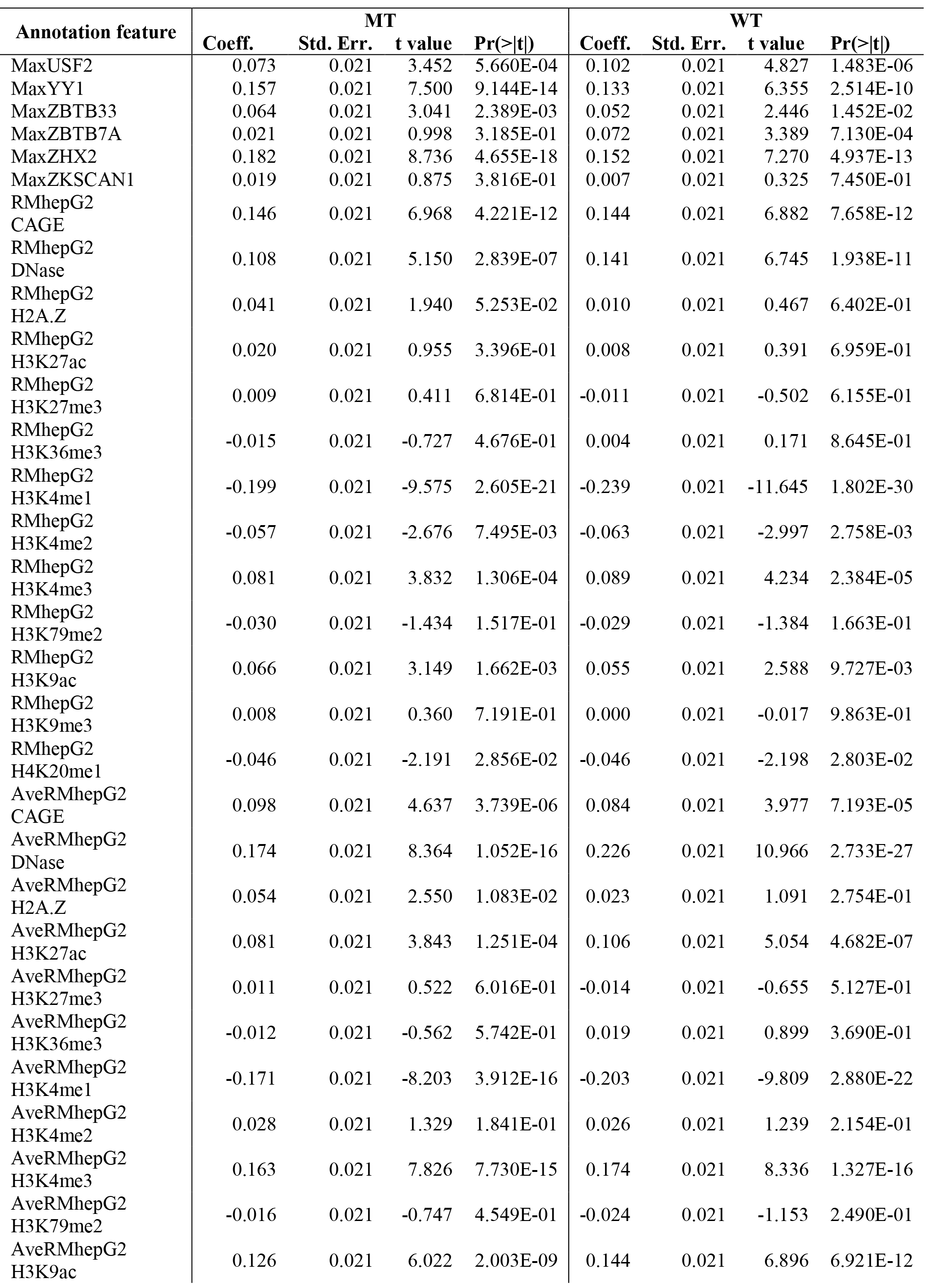

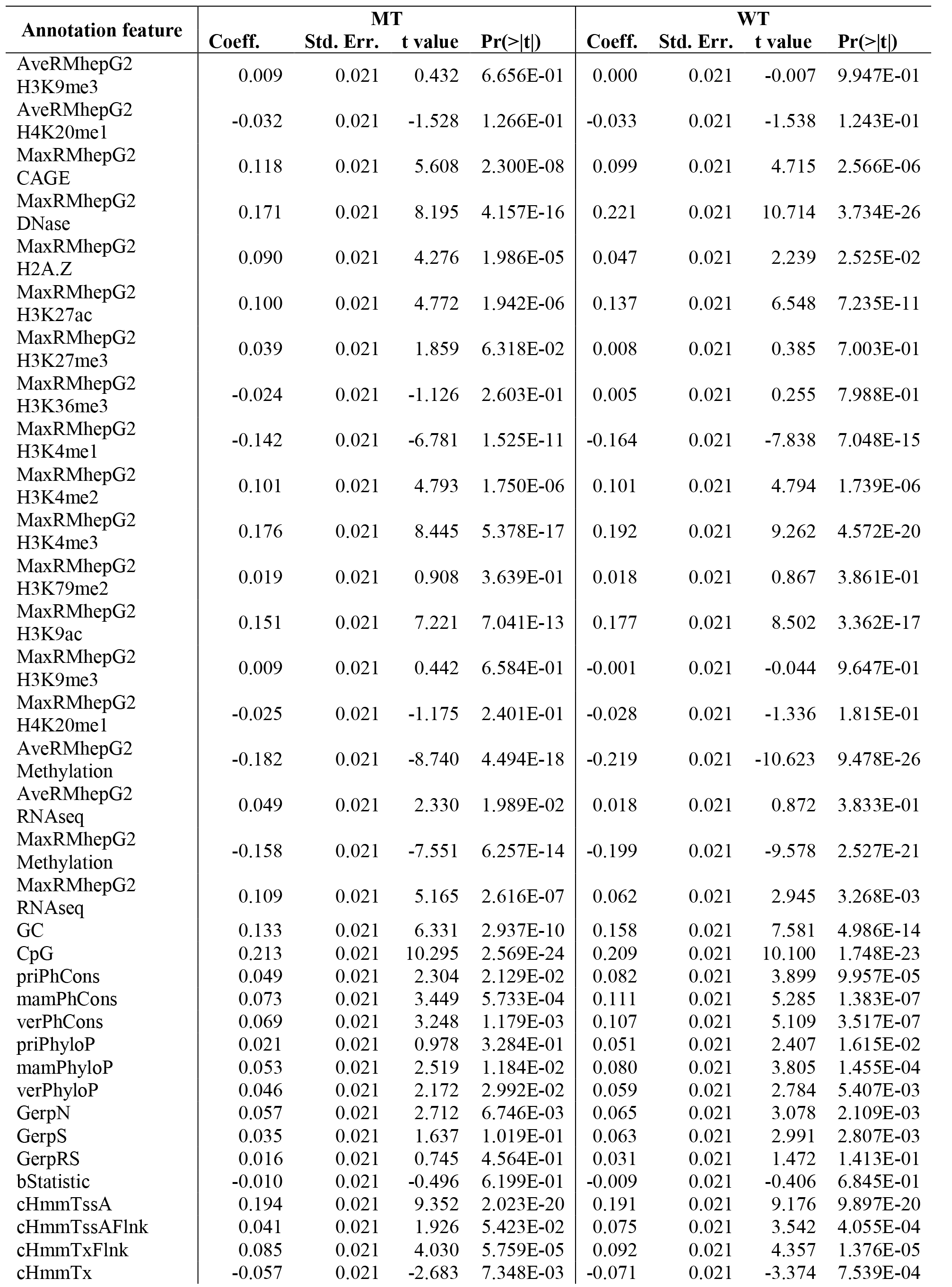

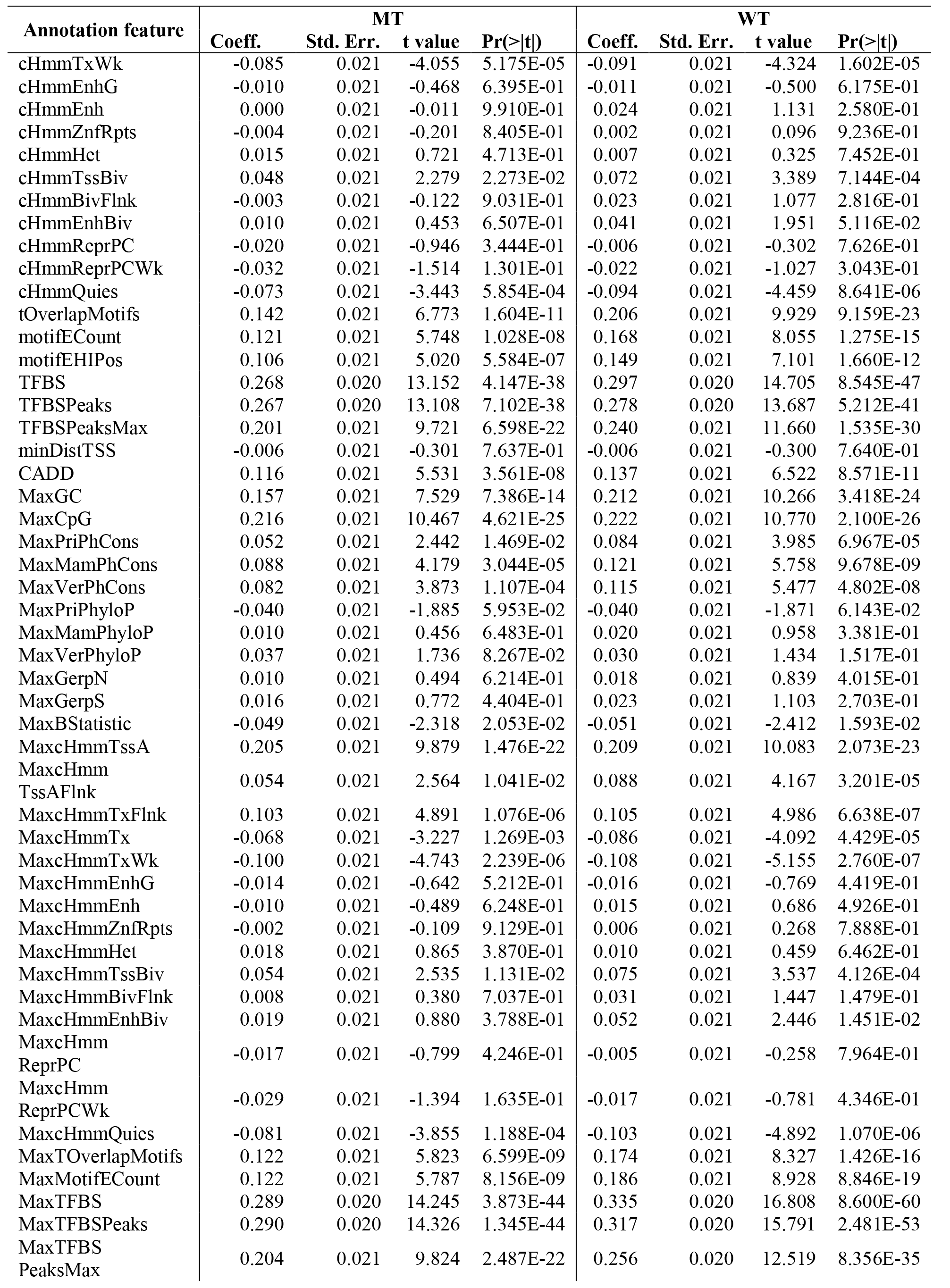

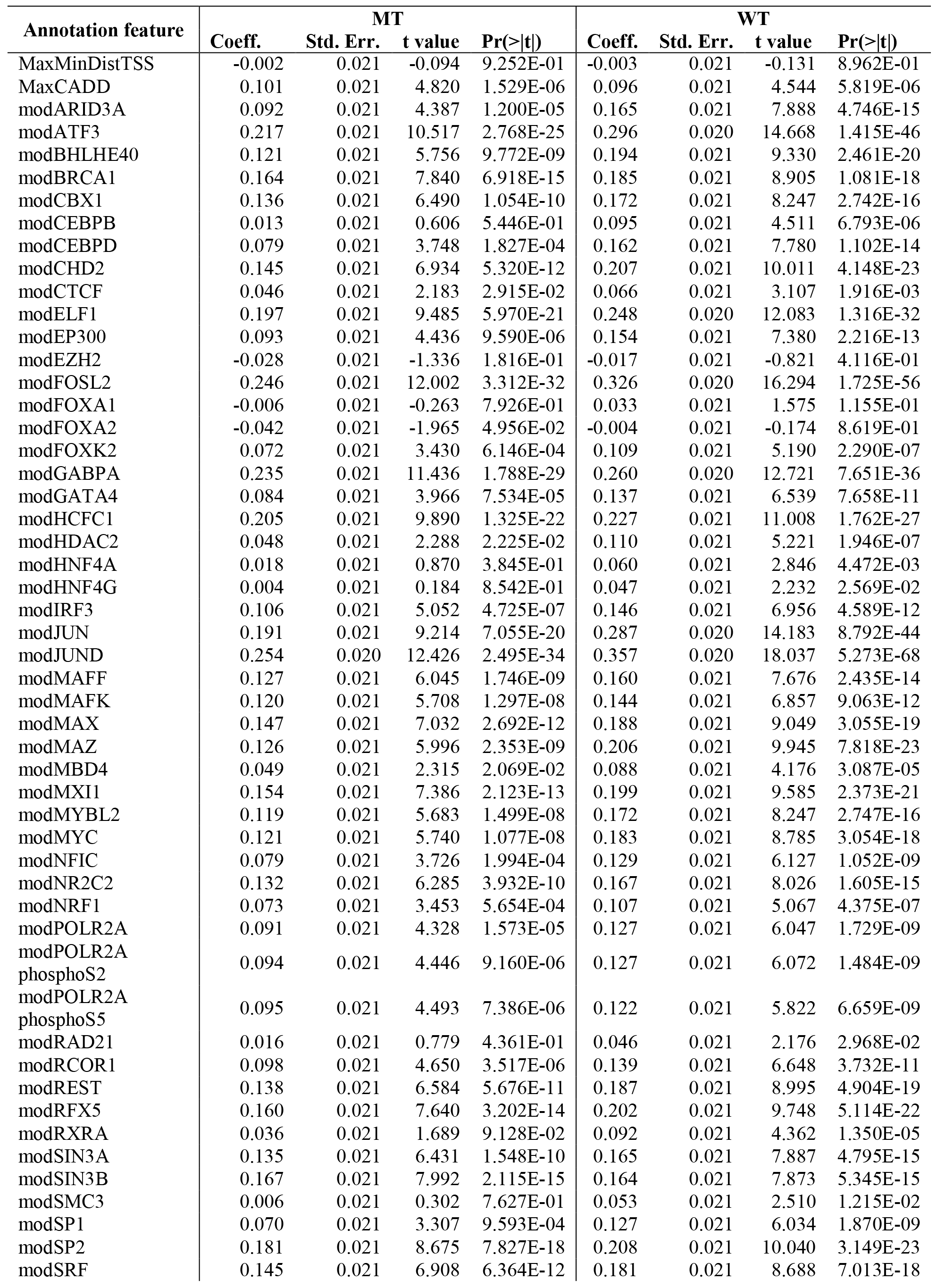

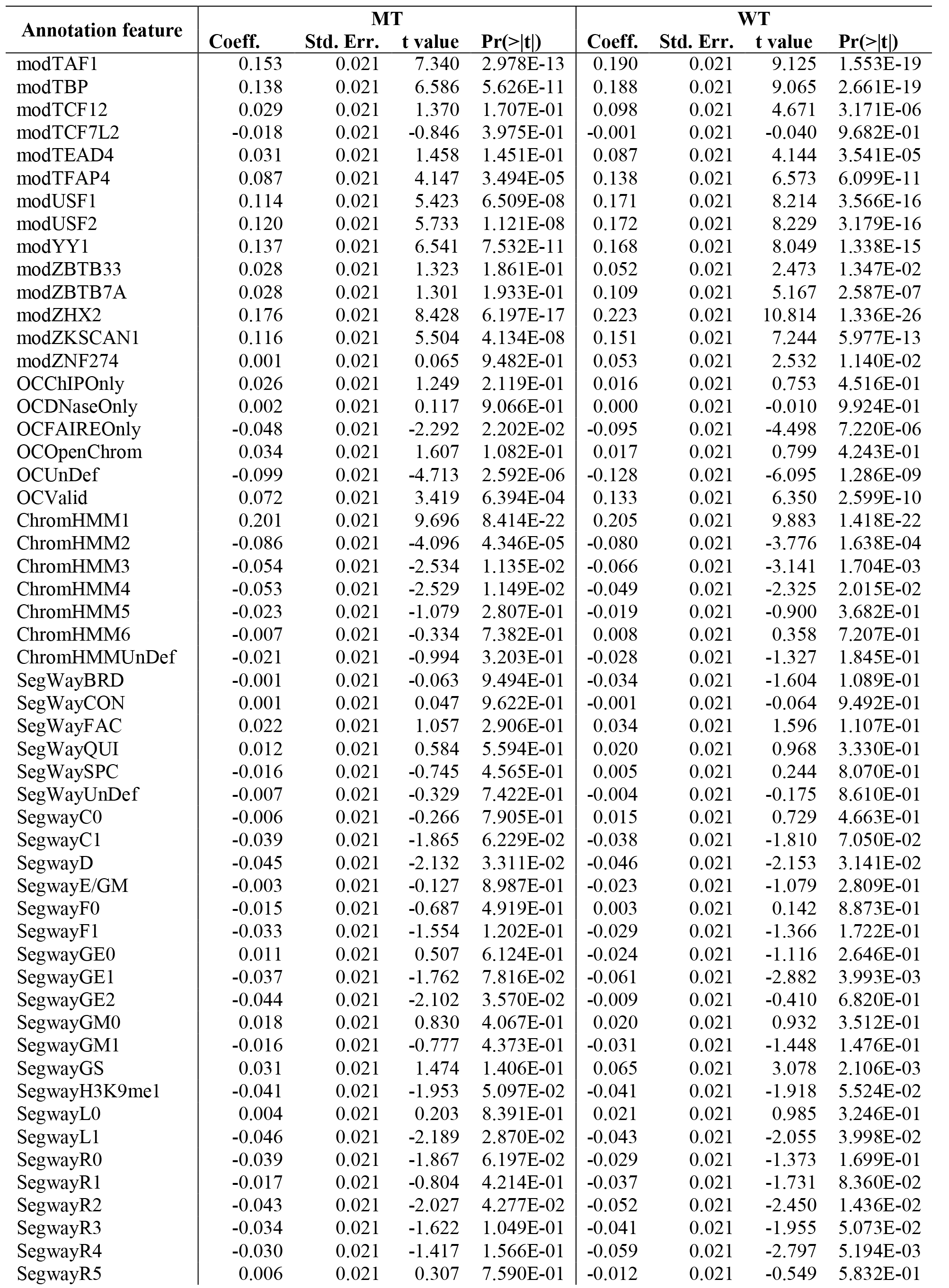

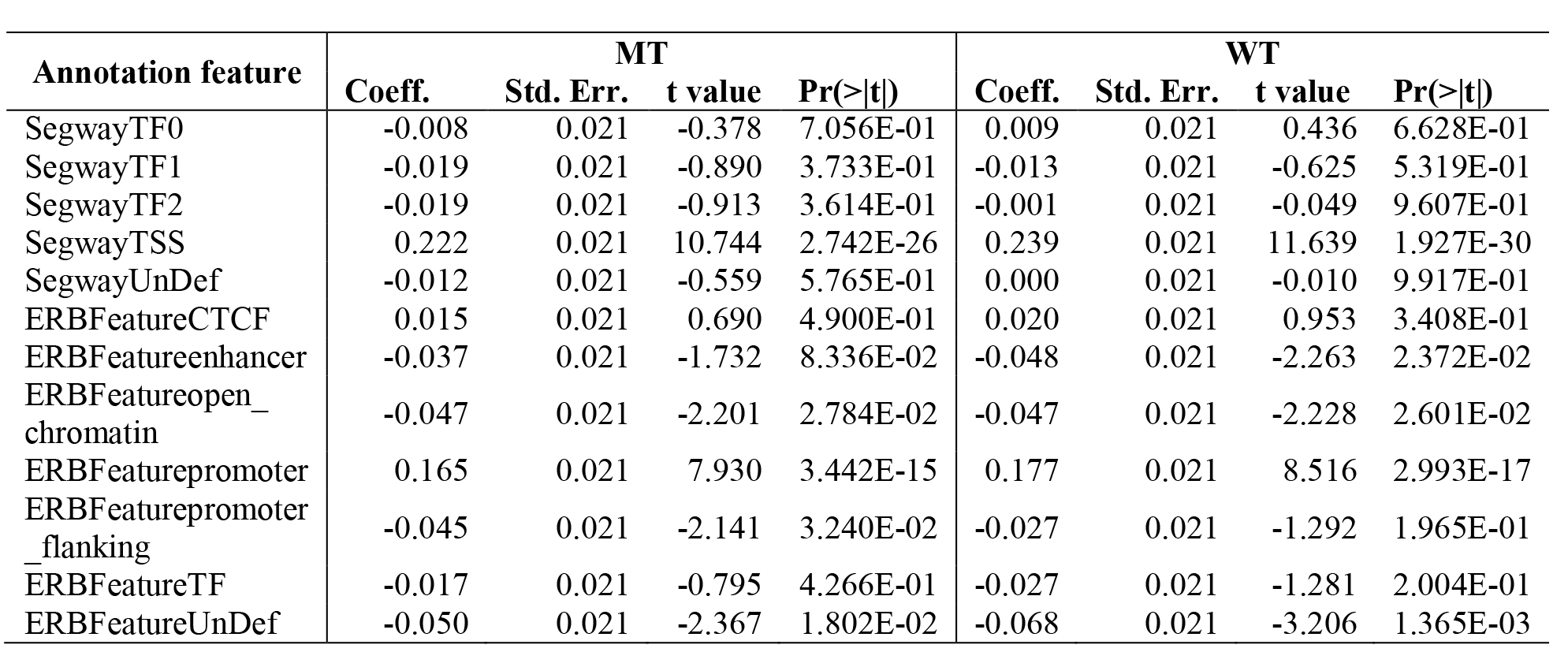

### Additional files

*File 1*: Annotated plasmid sequence file in GenBank Flat File Format.

*File 2*: Designed array oligo nucleotide sequences (gzip-compressed text file)

*File 3*: Text file listing out annotations used for prediction of observed RNA/DNA ratios.

